# Cancers modulate p53 truncal neoantigen display to evade T cell detection

**DOI:** 10.1101/2025.11.26.690562

**Authors:** Koji Haratani, Bruce Reinhold, Jonathan S. Duke-Cohan, Caroline G. Fahey, Kemin Tan, Robert J. Mallis, Alexander Gusev, Kenneth L. Kehl, Jia Luo, Elizabeth L. Holliday, Daniel J. Masi, Allyson Karmazyn, Katarzyna J. Zienkiewicz, Connor J. Hennessey, Rafael B. Blasco, Tran C. Thai, Grace M. Gibbons, Sophie Kivlehan, Patrick Lizotte, Cloud P. Paweletz, Andrew J. Aguirre, Keith L. Ligon, Roberto Chiarle, Matthew J. Lang, David A. Barbie, Ellis L. Reinherz

## Abstract

*TP53* mutations are early truncal events across cancers^1,2^. These are perceived to encode tumour-specific neoantigens representing prime cytotoxic T lymphocyte (CTL) targets^3,4^. However, studies systematically examining the physical cell surface display of p53 peptides bound to major histocompatibility complex molecules (pMHC), their relative antigenicity, and resultant immunogenicity have yet to be conducted. Here, we develop an epitope discovery platform using p53-reconstituted lung cancer cells as well as various tumour cells as pMHC sources. Combining data-independent acquisition mass spectrometry (MS), nanoscale chromatography, and peptide detection based on probabilistic measure and three-dimensional ion visualization techniques allows attomole sensitivity identification of pMHCs. This approach excluded ∼97% of algorithm-based virtual p53 immunopeptidomes, highlighting that only a few p53 pMHCs can be presented by common human MHC (human leukocyte antigen, [HLA]) alleles. Strikingly, surface expressed neoantigens are restricted to the corresponding set of such limited self-p53 peptide arrays and unaffected by enhancing p53 proteasomal turnover. Further curtailment of MS-validated, high affinity p53 neoepitopes that are structurally deviant from self-pMHC occurs in established tumours due to immune selection against the antigen presenting MHC allele or by a novel mechanism involving p53 neoepitope destruction by endoplasmic reticulum aminopeptidase 1 (ERAP1). In contrast, given the extremely weak MHC affinity and resultant short-lived cell surface pMHC expression, the common p53 neoepitope R175H/HLA-A*02:01 escapes immune selection despite CTL with high quality T-cell receptors. Rigorous tumour-protective immunoediting makes effective truncal neoepitope targeting a challenge, requiring attentive MS analysis and functional vetting to focus protective cytolytic responses.

## Main text

Inactivating mutations in *TP53*, the gene encoding the p53 tumour suppressor protein, are the most common genetic drivers observed across all forms of cancer^1,2^. Since p53 functions to protect cells from DNA damage and to promote apoptosis, its mutation often occurs early during malignant cell transformation, accounting for its presence in all tumour subclones and so called “truncal” nature^1,2,5^. Immunotherapeutic targeting of truncal neoantigens is viewed as particularly advantageous relative to non-truncal passenger mutations, as the latter can lead to tumour escape under immune pressure by selecting clonal variants that lack the neoantigen^5,6^. Furthermore, as *TP53* missense mutations characteristically increase p53 protein expression by a well described mechanism^1,2^, p53 neoantigens are thought to represent an attractive focus for cancer immunotherapies including T cell receptor–engineered T cells (TCR-T)^3^ or CD3/neoantigen bispecific T cell engagers (BiTEs)^4^. Indeed, over the past decade, potentially actionable tumour-derived peptides bound to major histocompatibility complex molecules [pMHCs, or pHLAs for the human leucocyte antigen complex] have been identified primarily using bioinformatic-based reverse immunology approaches for targeting in the clinic, such as an HLA-A*02:01–bound p53^R175H^ peptide (^168^HMTEVVR<u>H</u>C^176^)^3,4,7–10^. If these truncal neoantigens can be robustly displayed as high-quality epitopes, however, the fundamental question of how cancer cells could overcome initial immune surveillance is raised.

Computational prediction models using artificial neural networks in conjunction with tumour-derived DNA and RNA sequencing data are commonly used to define pHLA epitopes^11–14^. However, peptide binding and elution algorithms are useful but insufficient given the complexity of antigen presentation^13,15^. The endogenous peptide processing machinery of normal cells comprises protein proteasomal targeting and cleavage followed by transporter associated with antigen processing (TAP)-mediated delivery of peptides to the endoplasmic reticulum (ER)^11,15^. Attendant peptide trimming by ER aminopeptidases (ERAPs) occurs prior to surface pHLA transport and normal cell surface display^11,15^. While these components are operative in cancer cells, additional factors dysregulating this machinery may negatively impact tumour display of neoantigens^11,16,17^, especially under immune selection pressure during early tumourigenesis. Furthermore, dissimilarities related to neoepitope immunogenicity consequent to structural diversity of pHLA, variance in peptide-HLA binding affinities and divergent epitope copy numbers per cell, as well as TCR signaling performance in each patient’s repertoire will affect downstream T cell activation and anti-tumour responses^14,18–20^.

Therefore, we performed comprehensive profiling of both normal and tumour cells to interrogate the p53 immunopeptidome, utilizing a Poisson detection attomole-level ultrasensitive liquid-chromatography data independent acquisition mass spectrometry (LC-DIA-MS) platform^13,21,22^. Profiling was followed by interrogation of epitope structure and neoantigen-specific TCR generation for rigorous functional analyses relevant to antigenicity and immunogenicity. This conjoint approach identified novel p53 neoepitopes capable of being cell surface displayed but masked through precise mechanisms exploited by tumours to thwart effective T cell attack.

### p53 epitope characterization by Poisson LC-DIA-MS

Our approach mitigates certain limitations of current MS methods as described in Supplementary Discussion 1, including DIA-MS that systematically fragments all ions across defined mass-to-charge windows. Distinct from conventional feature extraction methods in DIA, we model peptide detection as sampling a stochastic Poisson process. Features derived from extracted ion chromatograms, Poisson chromatograms, ion surface maps and Poisson surfaces are used in our LC-DIA-MS detection. The approach leverages cost-effective synthetic peptide pools to provide both precise fragmentation and retention time references, improving peptide detection. Additionally, MS sensitivity is enhanced by deploying nano-LC with flow rates under 10 nanoliters/minute using monolithic 20-micron capillary columns and distal-coated tips pulled from 10-micron ID capillaries. Electrospray with very low flow rates and a small emitting aperture generates minute aerosol droplets which are widely recognized to significantly improve ionization efficiency^23–25^. This attomole-sensitive LC-DIA-MS can enable detection of a few or even one copy of a specific pHLA per cell as we showed previously^21^.

Utilizing this LC-DIA-MS technology, detailed wild-type (wt) and mutant (mut) p53 immunopeptidome data were acquired from a broad range of samples (Fig. 1a). All class-I HLA-presented peptides were isolated by pHLA immunoprecipitation followed by acid pH peptide release from patient-derived cancer cells or tumour tissues bearing mutant p53 (p53^mut^) and p53^mut^-transduced cell lines in addition to circulating immune cells as a normal counterpart. This comprehensive p53 immunopeptidome facilitated a subsequent immunogenicity profiling of p53 truncal neoepitopes based on large clinical genome databases, immunological studies using neoantigen-directed TCR-T, and pHLA structure determinations.

**Fig. 1.**
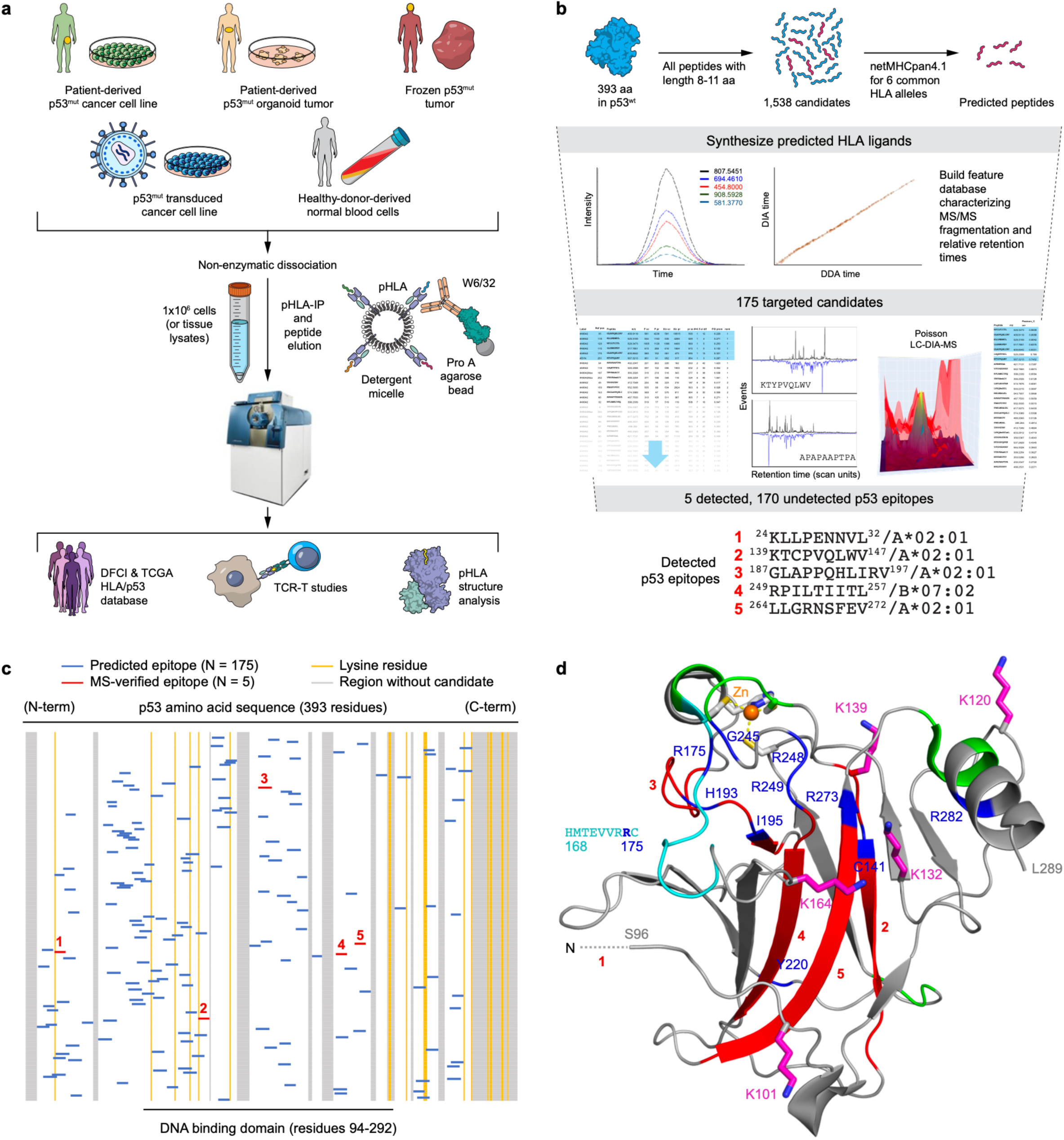
p53 immunopeptidome characterization platform identifies limited antigen presentation. **a**, Overview of the p53 immunopeptidome profiling platform involving cell sourcing, processing and analyses. **b**, All 8–11-mers from wt p53 and common p53 mutations that were predicted by netMHCpan4.1 to bind the HLA alleles of H2228 were synthesized in a pool and without purification. DDA/DIA analyses generated 363 reference sets combining fragmentation patterns (left curve) and elution positions (right curve) that are assembled into a target file for Poisson analyses. The full target file lists multiple charge states, methionine oxidation as well as some mutations. 175 distinct peptides from wt p53 were covered. Samples were analyzed against the full target set, generating 363 pairs of extracted ion chromatograms (XICs, black traces) for the precursor ions and Poisson chromatograms (inverted blue traces) for fragment sampling. Extracted features from these chromatograms were used to eliminate peptides with out-of-range features (for example, too far off the elution line) and remaining peptides were sorted by feature (list on left). Those peptides with the top ranking are correlated with detection (light blue highlight). The higher-ranking peptides are then also analyzed by extracting three-dimensional ion map surfaces (colored peak representations) and calculating a Poisson surface to compare with the precursor ion map. Some false positives are identified when these features are ranked (list on right). See Supplementary Discussion 1. These analyses identified five p53 peptides consistently observed with H2228 p53^wt^ transduction. **c**, Candidate peptides from p53 (top to bottom) and their positions in the protein (N- to C-terminus, left to right) shown in blue with the five in red physically detected by MS (Supplementary Data File 2). Lysine residues as potential ubiquitination and/or acetylation sites are in yellow, and regions without candidates are in grey. **d**, p53 DNA binding domain structure <u>(PDB code:2OCJ)</u> with indicated newly defined epitopes identified in red, R175H epitope in mint, and hot spot SNV mutant residues in blue. Green segments lack any candidate epitopes.

To characterize the HLA-restricted physical display of p53 epitopes, we initially analysed the H2228 non–small cell lung cancer (NSCLC) cell line in which we recently discovered anaplastic lymphoma kinase (ALK) epitopes^22^. Since endogenous p53 is truncated (p53^Q331*^) in H2228 cells, we first stably transduced them with full-length wt *TP53*. H2228 cells were previously reported to express six alleles, namely, HLA-A*02:01, HLA-A*03:01, HLA-B*07:02, HLA-B*38:01, HLA-C*07:02, and HLA-C*12:02^26^ which are commonly observed across races or ethnicities^27^. To generate MS/MS reference patterns for Poisson detection, all candidate 8–11-mer p53^wt^ peptides predicted by NetMHCpan-4.1^28^ to associate with one or more of the H2228 HLA alleles were synthesized, using liberal criteria including binding affinity (BA) or MS-eluted ligands (EL) scores of ≤2%. MS/MS reference fragmentation patterns and relative retention times were thus generated for 175 predicted p53^wt^ candidate peptides (Supplementary Data File 1).

Utilizing both two-dimensional chromatograms and three-dimensional surfaces related to the fragment and parent ions, features were extracted and used to rank the probabilistic presence of each peptide (Supplementary Discussion 1). As a result of this ultrasensitive Poisson detection LC-DIA-MS, we defined ^24^KLLPENNVL^32^/A*02:01, ^139^KTCPVQLWV^147^/A*02:01, ^187^GLAPPQHLIRV^197^/A*02:01, ^249^RPILTIITL^257^/B*07:02, and ^264^LLGRNSFEV^272^/A*02:01 as robust p53^wt^ epitopes (Fig. 1b). Four of these five p53^wt^ epitopes, except for the novel ^139^KTCPVQLWV^147^/A*02:01, have been independently identified by other MS studies (IEDB.org), supporting the validity of our LC-MS platform. Importantly, while the bioinformatically predicted peptides with high affinity extensively covered all functional p53 domains (Fig. 1c, and Supplementary Data File 2), only five of the 175 targeted candidates (2.86%) were validated being as cell-surface displayed epitopes.

To ensure display of all HLA alleles, we sequence-verified five of the six HLA alleles in H2228 cells and identified the remaining allele as a non-functional variant of A*03:01, confirming absence of HLA-A*03 protein expression on the cell surface (Supplementary Data File 3). Accordingly, none of a set of reference fragmentation patterns downloaded from SWATHAtlas.org and identified as HLA-A*03:01 binders by NetMHCpan-4.1 were detected in H2228 cells by Poisson LC-DIA-MS. We therefore examined the independent HLA-A*02:01^+^ HLA-A*03:01^+^ HLA-B*07:02^+^ tumour cell line COV318 bearing full-length endogenous p53^I195F^ and readily identified non-p53 HLA-A*03:01 binders (Supplementary Data File 1 and data not shown). While we did not identify display of any of the predicted HLA-A*03:01-restricted p53 candidates, the p53^wt^ epitopes restricted by HLA-A*02:01 and HLA-B*07:02 were detected and shared with H2228 cells, confirming the robustness of our approach (Supplementary Data File 1).

Since we consistently observed physical detection of specific p53^wt^ epitopes from transduced H2228 cells as well as from COV318 cells endogenously expressing mutant p53, we additionally sought to validate the display of these peptides in activated T cells as a proxy for normal cells. For this purpose, we obtained circulating T cells from a healthy donor (RG2684) expressing HLA-A*02:01 and HLA-B*07:02 as well as HLA-B*40:02, HLA-C*05:01, and HLA-C*07:02. Poisson LC-DIA-MS validated endogenous display of this exact same set of p53^wt^ peptides restricted by HLA-A*02:01 and HLA-B*07:02 (Extended Data Fig. 1 and Supplementary Discussion 1), in addition to a distinct HLA-B*40:02-restricted p53^wt^ epitope ^342^RELNEALEL^350^. Overall, the consistency between MS results and retrospective HLA genotyping, across primary cells and multiple cancer cells (*vide infra*) corroborates the accuracy of our Poisson LC-DIA-MS platform.

### Epitope mapping onto the p53 structure

We mapped the locations on the X-ray structural model of the p53 DNA-binding domain^29^ of four of the five identified epitopes along with that of ^168^HMTEVVRRC^176^ whose mutation to R175H is the most common p53 mutant and was previously MS identified^4^ (Fig. 1d). One peptide, ^24^KLLPENNVL^32^ derives from the transactivation domain and is thus not visualised in this model. Supplementary Movie 1 shows the 3D rendering of the DNA-binding domain, highlighting the positions of the β-strands comprising epitopes 2, 4, and 5 as well as the loops containing epitope 3 and ^168^HMTEVVRRC^175^.

The positions of key common “hotspot” p53 missense mutations^30^ in these regions are highlighted in blue (Fig. 1d). Those variants disrupt the normal function of p53 by altering hydrogen bonding that supports the H1 helix structure and Zn^2+^ coordination site (R175 in Extended Data Fig. 2a), perturb structures of hydrophobic or hydrophilic pockets located in the layer beneath the DNA interaction surface of the binding domain (C141, H193 and I195 in Extended Data Fig. 2b–d), mitigate DNA interaction through alteration of direct contact residues (R248 and R273 in Extended Data Fig. 2e,f), or disorder the L3 loop (R249 in Extended Data Fig. 2e) that in turn impacts DNA binding by altering the key double stranded DNA contact residue R248. As the p53^wt^ DNA binding domain is critical for tumour suppressor function, these mutations individually block vital anti-cancer protective mechanisms^1,2^. We note that vicinal lysine residues such as K101 and K132 as well as K139 and K164 could serve as acetylation sites^31^ or, alternatively, potential ubiquitination sites for proteasome 19S regulatory particle interaction, although the latter are thought to primarily reside at the C-terminal K370–386 segment^31,32^. Thus, structural features accounting for processing of these peptides and their preferential display is not immediately obvious, underscoring the importance of MS methods in defining the p53 immunopeptidome.

### Paucity of targetable common p53 neoantigens

We next expanded our pipeline to interrogate a broad set of p53 neoantigen candidates predicted to bind these same HLA alleles, by expressing each p53 mutant variant in H2228 cells and performing LC-DIA-MS following pHLA immunoprecipitation. Clinically well-recognized p53 variants as common “hotspot” mutations—such as p53^R175X^, p53^Y220X^, p53^G245X^, p53^R248X^, p53^R273X^, and p53^R282X^ as reported in IARC and GENIE database^30,33^—were assessed. p53^R158X^ and p53^E285X^ were also included due to their frequency in immune checkpoint inhibitor responsive cancers including NSCLC and urothelial carcinoma^33–35^. All potential 8–11-mers (418 peptides) from these hotspot mutations were screened for H2228 HLA alleles using the same liberal NetMHCpan-4.1 criteria (EL or BA rank of <2%), yielding 28 candidates. We also hypothesized that neoantigens from the corresponding p53^wt^ peptides we identified would have a high probability of presentation. Therefore, predicted high affinity peptides from p53^C141X^, p53^H193X^, p53^I195X^, and p53^R249X^ variants were also included despite their relative lower frequency in the clinic. Confident fragmentation patterns and elution positions were successfully generated for most of these candidates (Supplementary Data File 1). However, the putative p53^R175H^ neoantigen ^168^HMTEVVR<u>H</u>C^176^ (as well as its wt counterpart ^168^HMTEVVRRC^176^) could not be included in our LC-MS, due to poor chromatographic behavior of the alkane-modified polystyrene-divinylbenzene monolith for hydrophilic peptides, and thus precluding confident detection (Supplementary Discussion 2). Nevertheless, presentation of this 9-mer neoantigen on HLA-A*02:01 was previously confirmed by LC-MS using a different chromatographic column coupled with selective reaction monitoring (SRM) and heavy label yielding ∼1–2 copies per tumour cell^4^. Methionine oxidation during elution also limited definitive detection of several other peptide candidates detailed in Supplementary Discussion 2.

In contrast to positive detection of neoantigen variants derived from p53^C141X^, p53^H193X^, p53^I195X^ and p53^R249X^, we were unable to detect neoepitopes from any of these other candidates (Extended Data Fig. 3). Panel a therein shows positions of detected epitopes, Panel b summarizes results of wt and mutant peptide concordance and Panel c gives the ranked 2-D chromatographic parameters of the various detections. The findings reveal a highly significant correlation between p53 peptides found in our immunopeptidome analysis and their corresponding point mutant neoantigens in the context of the alleles examined (*P* = 0.0039). For example, both H193Y and I195X neoantigens are detected upon variant p53 transduction and reside in the same peptide segment matching the displayed normal counterpart ^187^GLAPPQHLIRV^197^, confirming that wild-type protein immunopeptidome analysis can inform more efficient screening of true positive cancer neoantigens. Taken together, these findings suggest that tumours avoid displaying “hotspot” neoepitopes on common HLA alleles, as the explanation for their clinical frequency. An exception is ^168^HMTEVVR<u>H</u>C^176^/A*02:01, the most common clinical variant^30^ p53^R175H^, that becomes a focus of our further investigation below.

### Degraders do not enhance p53 immunopeptidome display

Given the restricted nature of p53 epitopes, we assessed whether we could broaden and/or enhance surface display using the cdTAG fusion protein system to enforce robust p53 proteasomal degradation^36^ (Extended Data Fig. 4a). We treated p53-cdTAG–transduced cells with the bifunctional degrader molecule dTAG47, which led to significant cdTAG-p53 turnover as compared with the control (Extended Data Fig. 4a). Enhancing p53 protein turnover, however, does not impact the abundance of any of the displayed p53 peptides (Extended Data Fig. 4b) nor create *de novo* additional p53 peptides searched for in our analysis (Fig. 1c).

To exclude the possibility that failure of the cdTAG47 degrader system to induce peptides was a consequence of the chimeric protein, the drug employed, and/or unique to p53, we examined a different degrader in an orthogonal system, utilizing an inhibitor that degrades endogenous EML4-ALK in H2228 cells. Having previously identified that H2228 cells present one ALK epitope, RPRPSQPSSL/B*07:02, whereas lymphoma cells additionally present three other ALK epitopes (VPRKNITLI/B*07:02, IVRCIGVSL/B*07:02, and AMLDLLHVA/A*02:01)^22^, we utilised the ALK-specific degrader TL-13-112^37^ to assess the effect upon ALK peptide display in H2228 cells (Extended Data Fig. 4c–g and Supplementary Data File 4). TL-13-112 treatment did not significantly impact quantitative augmentation of RPRPSQPSSL or induce any lymphoma-related epitopes (Extended Data Fig. 4g). Together, these data are consistent with the already markedly high turnover rate of proteins in the cytosol relative to pMHC complexes (on average 10,000-fold greater)^38,39^ amongst other parameters, likely contributing to the inability to manipulate this further pharmacologically.

### Further p53 neoepitope winnowing on tumours

Despite their clinical uncommonness, our newly identified p53 neoepitopes were investigated as targetable truncal neoantigens. To that end, we obtained clinically established tumours with p53^I195X^/HLA-A*02:01 that had been fresh frozen in the DFCI tumour bank for our LC-DIA-MS. We identified an ovarian cancer brain metastasis (10-417-2654) with p53^I195F^ mutation reported as *A*02:01^+^* by a high-resolution targeted HLA exome sequencing (Supplementary Data File 5). Surprisingly, however, MS detection of ^187^GLAPPQHL<u>F</u>RV^197^ was negative (Extended Data Fig. 5a). Moreover, all the other p53-derived A*02:01 ligands as well as A*02:01-restricted non-p53 positive control peptides were absent. To interrogate the mechanism underlying the global absence of A*02:01-binding peptides, the tumour’s coding genome and transcriptome were examined by whole exome sequencing and RNA-Seq, respectively. This revealed minimal exon reads and transcript reads for exons 1–3, implicating a locus-specific genetic defect that impacts intact HLA-A*02:01 expression and dependent immunopeptidome display (Extended Data Fig. 5b). Here it appears that the ovarian cancer exploited *HLA-A*02:01* gene deficiency in the context of the p53^I195F^ driver mutation to prevent surface display of a potentially immunogenic truncal neoepitope.

We next examined neoantigen display by LC-DIA-MS in two patient-derived p53^I195X^ cancer cell lines, PANFR0583 which is a pancreatic ductal adenocarcinoma (PDAC) organoid from a DFCI PDAC organoid bank^40^, as well as the ovarian epithelial serous carcinoma COV318 cell line whose p53^wt^ epitopes were already identified (Supplementary Data File 1). Sanger sequencing of cDNA extracted from these two cell lines confirmed the *TP53^I195X^* mutations (*TP53^I195T^*for the PANFR0583 and *TP53^I195F^* for the COV318) (Supplementary Data File 6a). *TP53* gene and p53 protein expression were also confirmed by quantitative RT-PCR and immunoblotting (Supplementary Data File 6b,c), showing substantial p53 expression in comparison with the primary T cells from the healthy donor RG2684 from which ^187^GLAPPQHLIRV^197^/A*02:01 p53^wt^ antigen was clearly detected. Importantly, flow cytometry combined with targeted-exome sequencing also verified A*02 expression on the cancer cell surface as comparable to that of H2228 cells (Supplementary Data File 6d). Strikingly, however, ^187^GLAPPQHL<u>T</u>RV^197^ pHLA neoepitopes were not detected by LC-DIA-MS in PANFR0583 (Fig. 2a), despite ready detection of^187^GLAPPQHL<u>T</u>RV^197^ in H2228 p53^I195T^ transductant (Fig.2b).

**Fig. 2.**
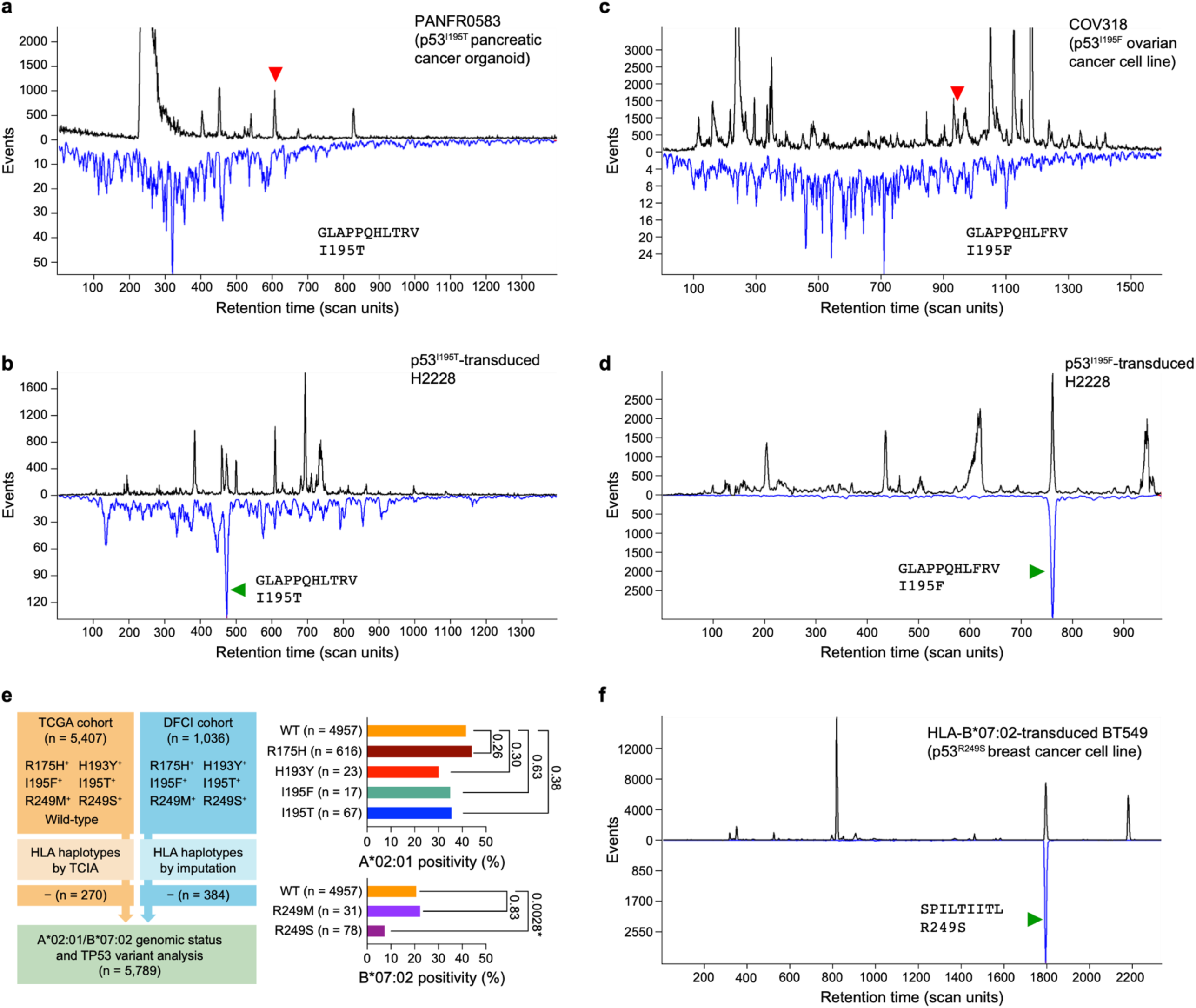
Mutant p53 ^I195T^ and p53 ^I195F^ peptide processing and surface expression differs in H2228 transductants versus cancer cells. **a–d**, Poisson detection plots for the indicated mutant p53 peptides. The top black trace is the extracted ion chromatogram (XIC), measuring precursor ion abundance. The inverted blue trace measures the number of fragments events that can be embedded in the corresponding DIA window (see Supplementary Discussion 1). Green and red arrowheads indicate MS detection or lack of detection, respectively, throughout figure panels. Genotyped p53^I195X^ peptides are not observed in the HLA-A2 supertype positive tumour samples (Panels a, c) but are observed in H2228 cells transduced with the corresponding p53^mut^ (Panels b, d). **e**, On the left, the flow diagram is shown for identification of patient samples bearing *TP53* variants and appropriate HLA-presenting alleles (patient recruitment details described in Methods). On the right, the bar graphs of HLA-A*02:01 (top) and HLA-B*07:02 (bottom) depict patient expression by percentage and sample numbers related to p53^mut^ peptides potentially presented by the indicated alleles. Tumours expressing wild-type p53 (WT) are used as controls. *P* values are calculated using Fisher’s exact test. **f**, BT549 breast cancer cell line genotyped with mutant *TP53^R249S^* expressed that neoantigen after transduction with cDNA encoding the restricting HLA-B*07:02 allele.

Likewise, COV318 did not present ^187^GLAPPQHL<u>F</u>RV^197^ (Fig. 2c), whereas H2228 p53^I195F^ transductant did present ^187^GLAPPQHL<u>F</u>RV^197^ (Fig. 2d). However, the other A*02:01 ligands from p53 protein including ^24^KLLPENNVL^32^ or ^139^KTCPVQLWV^147^, and ^264^LLGRNSFEV^272^ as well as the B*07:02-restricted ^249^RPILTIITL^257^ were expectedly presented by COV318, suggesting a p53^I195X^-specific neoepitope exclusion mechanism. Together, these patient-derived p53^I195X^ LC-MS data imply that host immune selective pressure could impact physical display of truncal p53 neoantigens as part of tumour evolution in the host.

### Negative selection of immunogenic neoepitopes

To further study the notion of immune selective pressure on clinical tumours harbouring truncal p53 neoepitopes, we obtained genotypic data on *HLA-A*02:01* and *HLA-B*07:02* alleles in cohorts of 5,879 patients with solid malignancies to compare allele frequency based on the specific p53 tumour mutation status (Fig. 2e). Clinical tumours with A*02:01-related p53 mutations generally did not appear to select against *HLA-A*02:01*. Likewise, p53^R249M^ tumours developed equivalently to p53^wt^ tumours in *HLA-B*07:02*^+^ host. Of note, however, p53^R249S^ tumours were significantly less frequent amongst *HLA-B*07:02*^+^ individuals.

To investigate this disparity further, we obtained the p53^R249S^ but *HLA-B*07:02*^negative^ breast cancer cell line BT549 and found by LC-DIA-MS that ^249^<u>S</u>PILTIITL^257^ is readily presented from the endogenous p53^R249S^ protein upon *HLA-B*07:02* transduction (Supplementary Data File 7 and Fig. 2f). Apparently, p53^R249S^ tumours predominantly rely on *HLA-B*07:02* genomic negativity to evolve clinically, whereas the p53^I195X^ tumours fail to present endogenous p53 neoepitope despite HLA-A*02:01 positivity. Additionally, unlike p53^R249S^, p53^R249M^ tumour evasion is impervious to neoantigen display even when restricted by the same *HLA-B*07:02* allele. Hence, a single amino acid difference may regulate antigenicity and/or immunogenicity, consistent with CTL detection of single amino acid changes in peptides^41–44^.

### Structural correlates of immunogenicity

Structural studies could offer insights into neoepitope evasion mechanisms, explaining why only a subset of epitopes are eliminated by a cancer. We generated recombinant HLA-A*02:01 and HLA-B*07:02 proteins in E. coli, refolded them with their p53^wt^- or p53^mut^-derived ligands, purified and crystallised the resultant complexes, and then carried out structural determinations (Extended Data Table 1 and Supplementary Discussion 3). Fig. 3a overlays HLA-B*07:02-related p53^wt^ epitope ^249^RPILTIITL^257^ with its related neoepitopes ^249^<u>M</u>PILTIITL^257^ and ^249^<u>S</u>PILTIITL^257^, revealing that the only difference in these peptides is observed at the p1 position. In this side view of the pHLA complexes, the α2 helix has been removed to reveal both buried and exposed residues. The Fig. 3b surface representation shows that while the N-terminal end of the peptide groove is similar for the p53^wt^ (1R) and p53^R249M^ (1M) complexes due to the upward pointing large side chains of both p1 residues, that of p53^R249S^ (1S) is distinct. Its small serine side chain is buried in the peptide-binding groove, allowing for a long bidentate salt bridge to form between R62 and E163 on α1 and α2 helices, respectively (Extended Data Fig. 6a,b). The local HLA-B*07:02 structure surrounding p1S creates a pocket readily able to host a charged or polar sidechain possibly from CDR3α or elsewhere in the approaching TCR Vα domain (Fig. 3c). The resulting singular focal structural change on the pMHC surface confers high immunogenicity on the p53^R249S^ neoepitope, likely disfavoring tumour development when displayed by the host in HLA-B*07:02. Contrast this neoantigen with p53^R249M^ that is so similar in antigenicity to the corresponding p53^wt^ peptide bound to HLA-B*07:02 that it would remain invisible to the immune system. Given self-tolerance mechanisms, expression of p53^R249M^ in patients even if presented by HLA-B*07:02 would be unlikely subject to immune editing. This proposition is based on the structural differences between p53^R249S^ and p53^R249M^ and is consistent with the contrasting HLA-B*07:02 selectivity observed in the clinical data (Fig. 2e).

**Fig. 3.**
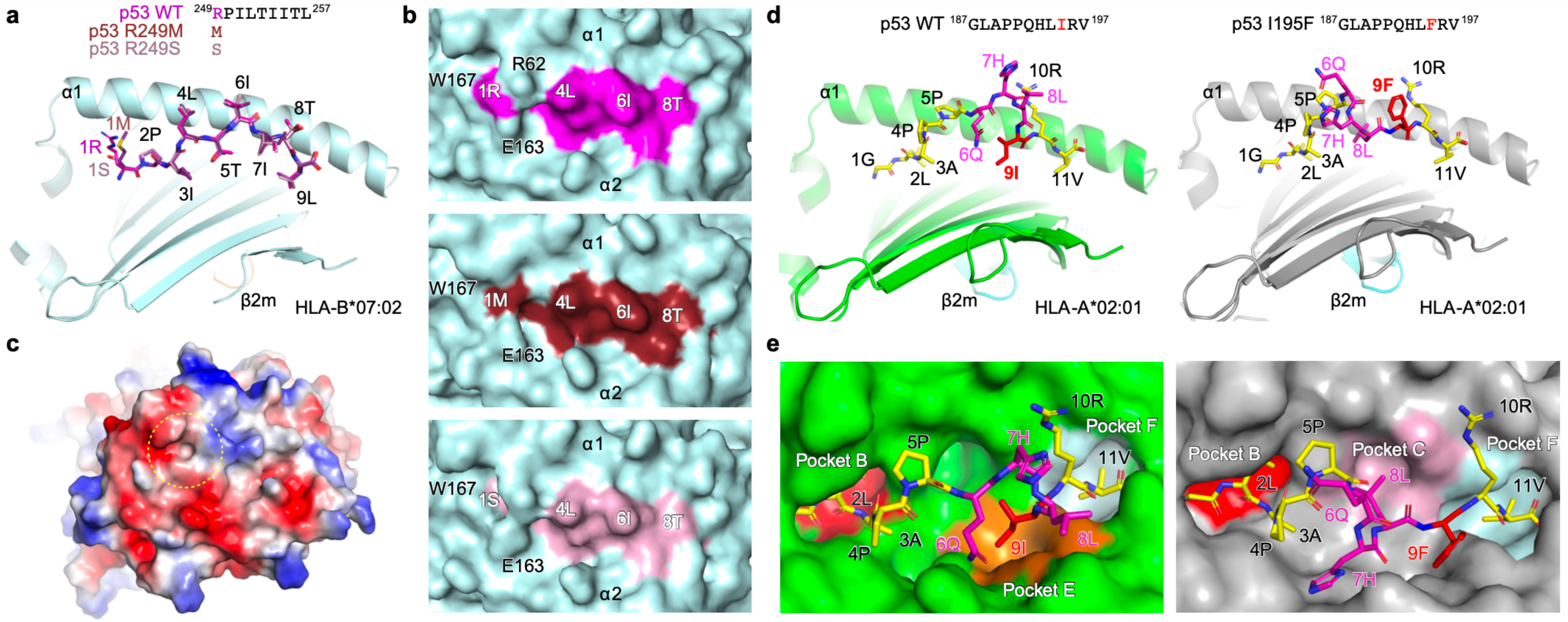
Distinct structural bases for strong immunogenicity of I195F/HLA-A*02:01 and R249S/HLA-B*07:02 pMHC complexes. **a**, Structural comparison of ^249^RPILTIITL^257^ (WT, magenta) peptide bound to HLA-B*07:02 with ^249^<u>M</u>PILTIITL^257^ (R249M, brown) and ^249^<u>S</u>PILTIITL^257^ (R249S, salmon) in the overlay showing nearly identical backbone conformations of the peptides. The α1 helix and β2m are labeled for orientation in this side view, and the α2 helix is removed for clarity. **b**, Top views display the p53 WT ^249^RPILTIITL^257^ peptide, and its R249M and R249S mutants (from top to bottom) within the HLA-B7 peptide binding groove, shown in a molecular surface representation. The surfaces contributed by HLA-B7, the WT peptide, the R249M peptide, and the R249S peptide are colored cyan, magenta, brown and pink, respectively. The peptide 1S residue in R249S is recessed below and to the left side of the salt link, forming a cavity as highlighted in panel c. **c,** Top view of the electrostatic potential surface representation of the p53 R249S ^249^SPILTIITL^257^/HLA-B7 complex. The colors on the molecular surface illustrate the distribution of electrical charges: red indicates negatively charged areas, blue represents positively charged regions, and white shows neutral zones. The dashed yellow circle highlights the cavity formed by the serine mutation in R249S. **d,e,** Comparison of X-ray crystallographic structures of p53^wt^ and p53^I195F^ 11-mer peptides (^187^GLAPPQHLIRV^197^ and ^187G^LAPPQHL<u>F</u>RV^197^). **d**, p53^wt^ ^187^GLAPPQHLIRV^197^ and p53^I195F^ ^187^GLAPPQHL<u>F</u>RV^197^ peptide/A2 complexes are on left and right, respectively, viewed from the side with the α2 helix of HLA-A*02:01 removed for clarity. Each of the 11 residues shown in stick format is labeled with the wild-type I and mutant F residues colored in red, and residues whose positions are structurally altered in magenta, respectively. The five N-terminal (GLAPP) and two C-terminal (RV) residues of the peptide with only minor conformational changes are in yellow. **e**, View from the TCR binding perspective onto the MHC groove with the α1 on top and α2 on the bottom in molecular surface representation.

As shown in Fig. 3d and Extended Data Fig. 6c,d, where ^187^GLAPPQHLIRV^197^ and ^187^GLAPPQHL<u>F</u>RV^197^ complexes are directly compared, despite just a single amino acid change in peptide sequence (I to F at the p9 position), there are conformational rearrangements in several side chains of exposed residues as well as the main chain. This broad impact of the p53^I195F^ mutation contrasts with those of the HLA-B*07:02 series above. The two p1-p5 and p10-p11 segments are virtually identical in both p53^wt^ and p53^I195F^ structures while, in contrast, the p6-p9 residues are very different. The 9F side chain of the neoantigen is too large to fit into the HLA-A*02:01 E pocket, unlike the 9I residue (Fig. 3d). Instead of being buried, the phenylalanine is surface exposed and featured for TCR recognition. In addition, the 8L residue of the neoantigen assumes an anchor position, entering the C pocket rather than the E pocket, and, in turn, rendering the main chain less proturberant, elevating 6Q and shifting 7H to a more lateral position. The view down onto the pMHC in Fig. 3e shows these anchor residue pockets as well as the exposed residues and mainchain that would be interrogated by the TCR. Thus, while both p53^wt^ and p53^I195F^ 11-mers bound to HLA-A*02:01 are canonical TCR ligands in structural terms and hence potentially antigenic, the former is a “self” ligand subjecting I195wt peptide-reactive thymocytes to undergo deletion, generating central immune tolerance. In contrast, I195F neoantigen is a foreign ligand whose distinct structure creates substantial potential immunogenicity.

### Truncal p53 neoepitope detection by T-cells

Since the mechanism of selective p53^I195X^ neoepitope concealment (^187^GLAPPQHL<u>X</u>RV^197^) despite intact HLA-A*02:01 expression was particularly enigmatic, we utilised a reverse immunologic approach to investigate TCR repertoires targeting these neoepitopes (Fig. 4a). After a 2-week coculture of naïve CD8^+^ T cells from an HLA-A*02:01 healthy donor (888299768) with autologous matured dendritic cells pulsed either with ^187^GLAPPQHLIRV^197^, ^187^GLAPPQHL<u>T</u>RV^197^, or ^187^GLAPPQHL<u>F</u>RV^197^ peptides, we readily expanded naïve CD8^+^ T cells recognizing ^187^GLAPPQHL<u>F</u>RV^197^/A*02:01 but not ^187^GLAPPQHL<u>T</u>RV^197^/A*02:01 (Fig. 4b), highlighting the antigenicity of p53^I195F^. Not unexpectedly, ^187^GLAPPQHLIRV^197^ induced no expansion. Subsequently, ^187G^LAPPQHL<u>F</u>RV^197^/A*02:01-specific T cells were sorted out by pHLA-dextramer staining, and subjected to single-cell RNA sequencing to determine full V(D)J sequences of paired TCR α/β chains and corresponding transcriptome. ^187^GLAPPQHL<u>F</u>RV^197^/A*02:01-dextramer–stained T cells comprised one dominant clonotype composed of TRAV17/TRBV28 coexpressing *IL2RA, HLA-DRB1* and *PCNA* genes consistent with activation through the prior TCR stimulation, in contrast to the polyclonal TCR repertoire from non-expanded autologous T cells derived from the ^187^GLAPPQHLIRV^197^ cultures (Fig. 4c, and Supplementary Data File 8).

**Fig. 4.**
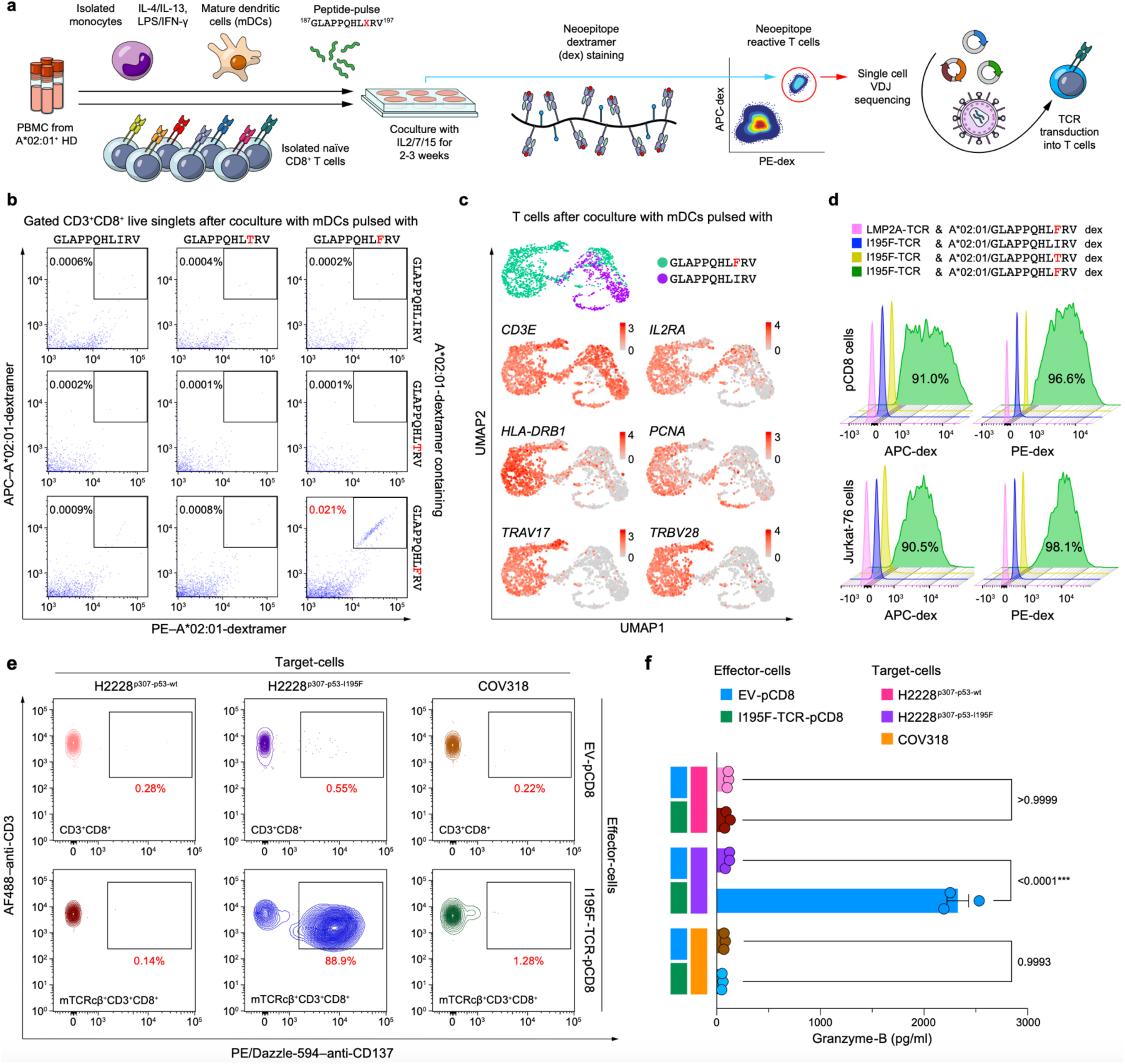
Identification of a I195F/HLA-A*02:01-specific TCR. **a**, Schematic for discovery of novel TCR clonotypes recognising p53 neoepitopes. Identified TCR clonotypes were lentivirally transduced into primary CD8^+^ T (pCD8) cells or Jurkat-76 cells to verify recognition of target neoepitopes. TCR constant regions were replaced with modified mouse constant regions (mTCRc). **b**, ^187G^LAPPQHL<u>X</u>RV^197^/A*02:01-dextramer binding positivity in CD8^+^ T cells after coculture with ^187^GLAPPQHL<u>X</u>RV^197^-pulsed mDCs as described in (**a**). Frequency of dextramer double–positive (CD3^+^CD8^+^) live singlets after dendritic cell peptide pulsing. **c**, Uniform Manifold Approximation and Projection (UMAP) plots of RNA-sequenced ^187^GLAPPQHL<u>F</u>RV^197^(I195F)/A*02:01-dextramer–sorted T cells (green) are shown with a negative T-cell control (purple), as indicated. Expression levels of *CD3E* are shown as pan-T cell markers, and those of *IL2RA*, *HLA-DRB1*, and *PCNA* are indicated as evidences of TCR pathway activation. **d**, Neoepitope reactivity by transduced human pCD8 (top) and Jurkat-76 (bottom) cells. Binding of PE- and APC-dextramers to TCR-transduced pCD8 or Jurkat-76 cells detected by flow cytometry. TCR recognizing CLGGLLTMV 9-mer from Epstein-Barr virus latent membrane protein 2A (EBV-LMP2A) was used as a negative control. mTCRcβ^+C^D3^+^CD8^+^ or mTCRcβ^+C^D3^+^ live singlets were gated to obtain dextramer-staining pattern for pCD8 or Jurkat-76 cells, respectively. Percentage binding to I195F-reactive T cells is indicated while for all other conditions, dextramer binding was <2.3%. Data representative of two independent biological replicates. **e**,**f**, Flow cytometry for CD137 expression (**e**) and ELISA quantitation of granzyme B (**f**) as T cell activation markers. Empty-vector (EV)-pCD8 or I195F-TCR-pCD8 cells were cocultured with target-cells as indicated for 24 h beforehand. Data representative of two independent biological replicates. **e**, The CD3^+^CD8^+^, or mTCRcβ^+C^D3^+^CD8^+^, live singlets were gated as indicated for CD137 expression depicted by contour plots with outliers as dots with CD137-positivity quantitation in red. **f**, Granzyme release quantitated by ELISA. Mean, standard errors of the mean (bars) and individual values from three technical replicates (filled circles) are shown. *P* values are calculated by one-way analysis of variance (ANOVA) with Tukey’s multiple-comparison test.

To validate this ^187^GLAPPQHL<u>F</u>RV^197^/A*02:01-stimulated TCR profile, the TCR was transduced into primary CD8^+^ T (pCD8) cells as well as into Jurkat-76 (J76) cells, a CD8^negative^ Jurkat subline lacking endogenous TCR α/β chains^45^. We utilised constructs where the TCR subunit constant regions were replaced by modified mouse orthologs to avoid mispairing with endogenous TCR repertoires and to mark the transduced cells (Supplementary Data File 9). The engineered TCR-pCD8 cells (termed I195F-TCR-pCD8) and comparably engineered J76 cells (termed I195F-TCR-J76) specifically distinguished the ^187^GLAPPQHL<u>F</u>RV^197^/A*02:01 neoepitope from wt or I195T counterparts as shown by selective dextramer binding (Fig. 4d). Activation of I195F-TCR-pCD8 cells by H2228^p53-I195F^ but not H2228^p53-wt^ target stimulator cells when assessed by CD137 upregulation or granzyme B secretion verified ^187^GLAPPQHL<u>F</u>RV^197^ presentation by HLA-A*02:01^+^ cells (Fig. 4e,f). Importantly, COV318 cells were unable to stimulate I195F-TCR-pCD8 cell activation, consistent with the negative LC-MS detection, and providing a functional tool with which to explore neoepitope suppression below.

### Tumour ERAP1 ablates a neoepitope

Since PANFR0583 and COV318 presented myriad peptides including those derived from p53^wt^, disruption of essential components of the antigen-presentation machinery (such as TAP1/2 or immunoproteasome subunits) could not be the basis for ^187^GLAPPQHL<u>X</u>RV^197^ neoepitope concealment. However, as all the p53^wt^-derived epitopes detected by MS analysis were 8–9-mers whereas the undetectable ^187^GLAPPQHL<u>X</u>RV^197^ was a 11-mer, we reasoned that endoplasmic reticulum aminopeptidase 1 (ERAP1) might be involved. Prior reports examining non-neoantigens revealed that ERAP1 activity increase can selectively attenuate long peptide (10–11-mers) epitope presentation^46,47^. On comparison with the H2228 cells, both PANFR0583 and COV318 expressed high levels of ERAP1 (Fig. 5a). Single nucleotide polymorphism genotyping of PCR amplicons, moreover, suggests that these tumours have hyperactive ERAP1 phenotypes (Supplementary Data File 10)^48^.

**Fig. 5.**
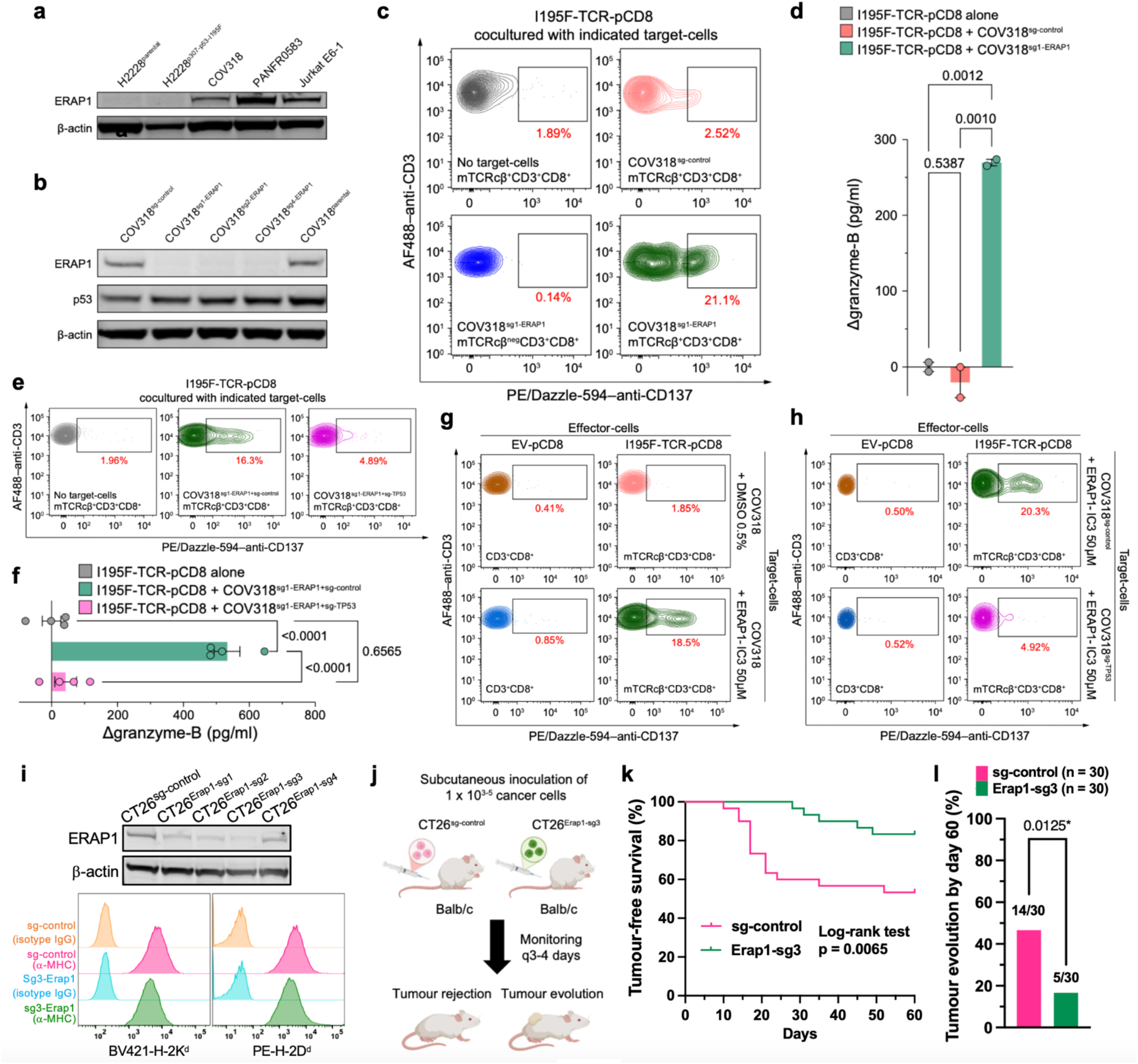
ERAP1 inhibition restores immunogenic truncal 11-mer neoepitope presentation. **a**, Immunoblotting of ERAP1 protein in COV318 and PANFR0583 versus parental and ^187G^LAPPQHLXRV^197^-expressing H2228 cells. Jurkat E6-1 cells were used as positive control given substantial ERAP1 expression and with β-actin as internal control throughout. **b**, Immunoblotting of ERAP1 and p53 proteins in COV318 with *ERAP1*-KO by three different sgRNAs (1, 2, and 4). p53 protein levels were not affected by *ERAP1*-KO. COV318 with sg-control (left) and parental COV318 (right) are negative controls. **c–h**, Flow cytometry data of CD137 expression (**c**,**e**,**g**,**h**) and ELISA data of granzyme B (**d**,**f**) as T cell activation markers. Empty-vector (EV)-pCD8 or I195F-TCR-pCD8 cells were cocultured with unmodified- or *ERAP1*-KO COV318 target cells as indicated for 24 h. **c**,**e**,**g**,**h**, CD137 expression by CD3^+^CD8^+^, mTCRcβ^+C^D3^+^CD8^+^, or mTCRcβ^−C^D3^+^CD8^+^ live singlets were gated and plotted/quantitated as described in Fig. 4e. Data are representative of two independent biological replicates in **a–f** and **h**. **d**,**f**, The results are presented as Δgranzyme-B ([value with target cells] – [value without target cells]). Mean values and standard error of the mean (SEM) (bars) and individual values from two to four technical replicates (filled circles) are shown. *P* values are calculated by one-way analysis of variance (ANOVA) with Tukey’s multiple-comparison test. **i**–**l,** *in vivo* studies**. i**, Immunoblotting of ERAP1 protein (top) and flow cytometry data of MHC-class I expression in CT26 cell lines using isotype controls to set backgrounds (bottom). **j**, Schematic of the CT26 *in vivo* study. Mice were subcutaneously inoculated with either CT26^sg-control^ cells or CT26^Erap1-sg3^ cells. Ten mice were used for each of three different cell concentrations (1 x 10^3^, 1 x 10^4^, or 1 x 10^5^), pooled from two independent experiments (five mice per concentration from each experiment), totaling 30 mice per CT26 cell line. **k**, Kaplan–Meier curves of tumour-free survival are shown. Events were defined as the first appearance of visible tumours. Vertical bars indicate censored data. **l**, Frequency of mice that developed tumours by day 60. The *P* value was calculated using the chi-squared test. sg, single guide RNA; DMSO, dimethyl sulfoxide.

To test whether ERAP1 function mediates this epitope concealment, we generated a CRISPR-Cas9-mediated *ERAP1*-knockout (KO) in COV318 cells by lentivirus transduction (Fig. 5b). Strikingly, as compared with COV318 cells expressing a control guide, COV318*^ERAP1^*^-KO^ cells were detected by I195F-TCR-pCD8 T cells while autologous non-engineered pCD8 T cell controls remained incapable of recognizing COV318*^ERAP1^*^-KO^ (Fig. 5c,d). Restoration of ^187^GLAPPQHL<u>F</u>RV^197^ neoepitope presentation by *ERAP1*-KO was confirmed by a second guide RNA (Extended Data Fig. 7a,b). Such I195F-TCR-pCD8 activation by *ERAP1*-KO was diminished by concurrent CRISPR-Cas9-mediated *TP53*-KO ensuring p53 specificity of TCR activation in the single *ERAP1*-KO (Fig. 5e,f and Extended Data Fig. 7c). These KOs did not alter HLA-A*02 cell-surface expression, reinforcing the direct effect of ERAP1 on peptide processing per se (Extended Data Fig. 7d). Furthermore, orthogonal treatment of COV318^parental^ cells with a chemical ERAP1 inhibitor (ERAP1-IC3)^49^ recapitulated the findings with COV318*^ERAP1^*^-KO^ (Fig. 5g,h).

Thus, cancer cells can exploit aminopeptidase trimming to avoid CTL targeting against certain immunogenic truncal neoepitopes of greater than average peptide length, perhaps providing a therapeutic mechanism to restore immunogenicity in those established tumours. Consistent with this proposition, we performed syngeneic mouse studies *in vivo* using the BALB/c-derived colon cancer cell line CT26 with or without *Erap1*-KO for implantation and observed that intact *Erap1* facilitates tumour formation even at very low numbers of injected tumour cells, relative to its absence (Fig. 5i–l and Extended Data Fig. 7e).

### Divergent p53^R175H^ versus p53^I195F^ antigenicity

p53^R175H^ is a clinically dominant missense variant whose physical and functional epitope display as well as TCR-based cognate recognition through X-ray crystallographic analysis are well-documented^4,43^. Yet, paradoxically, this neoepitope neither exploited negative HLA selection nor can invoke aminopeptidase cleavage (given its length) to escape immune pressure. Our independent experiments with KLE cells endogenously expressing p53^R175H^ and HLA-A*02:01 confirmed that the ^168^HMTEVVR<u>H</u>C^176^/A*02:01 neoepitope can be naturally presented for CTL recognition (Extended Data Fig. 8a).

To investigate how p53^R175H^/A*02:01^+^ cancer cells escape initial immune surveillance to foster clinical tumour development, we conducted parallel functional studies directly comparing the neoepitope quality of p53^R175H^ with that of p53^I195F^ since both are restricted by HLA-A*02:01. The binding affinity is poor between the p53^R175H^ 9-mer and HLA-A*02:01 (IC50, 4543.17 nM) relative to that for ^187^GLAPPQHL<u>F</u>RV^197^ (37.40 nM) as estimated by the NetMHCpan-4.1^28^. Accordingly, a previous MS analysis detected maximally only a few copies of the p53^R175H^ neoepitope per cell^4^.

We screened three published ^168^HMTEVVR<u>H</u>C^176^/A*02:01-reactive TCR clonotypes^43^, selecting R175H-TCR3 as the best functionally (Extended Data Fig. 8b). The HLA-A*02:01 surface expression levels and p53 protein expression levels were matched between H2228 cells transduced with p53^R175H^ or p53^I195F^ (Extended Data Fig. 8c). TCR expression levels were also coordinated between the two neoepitopes, with that of R175H-TCR3-pCD8 slightly greater for p53^R175H^ (Extended Data Fig. 8d). Nonetheless, antigen-specific T cell activation by the respective TCRs transduced into the same donor’s CD8^+^ T cells was inferior against H2228^p53-R175H^ compared with H2228^p53-I195F^ by T cell granzyme B and IFNγ secretion (Fig. 6a) and direct target-cell killing (Fig. 6b). In addition, *TAP1/2*-deficient HLA-A*02:01^+^ T2 cells lacking endogenous peptide presentation capability were used to normalise ^168^HMTEVVR<u>H</u>C^176^ and ^187^GLAPPQHL<u>F</u>RV^197^ loading onto HLA-A*02:01 by exogenous peptide pulsing at defined molar concentrations. A TCR specific for the Epstein-Barr virus latent membrane protein 2A (EBV-LMP2A) peptide CLGGLLTMV served as a positive control. The CTL activation by ^187^GLAPPQHL<u>F</u>RV^197^ was as good or better than established immunogenic viral epitope from EBV-LMP2A (Fig. 6c). In contrast, ^168^HMTEVVR<u>H</u>C^176^ required 100–1,000 fold higher concentrations to achieve a comparable level of IFNγ or granzyme B release, as confirmed using TCR-J76^CD8αβ^ despite higher expression levels of TCR, CD3, and CD8β for ^168^HMTEVVR<u>H</u>C^176^ (Extended Data Fig. 8e,f). Furthermore, T2 cells were killed less efficiently by R175H-TCR3-CD8 when they were pulsed with ^168^HMTEVVR<u>H</u>C^176^ at a low (1 nM) peptide-pulse concentration, whereas ^187^GLAPPQHL<u>F</u>RV^197^-pulsed T2 cells were readily lysed by I195F-TCR-CD8 from the same donor (Fig. 6d).

**Fig. 6.**
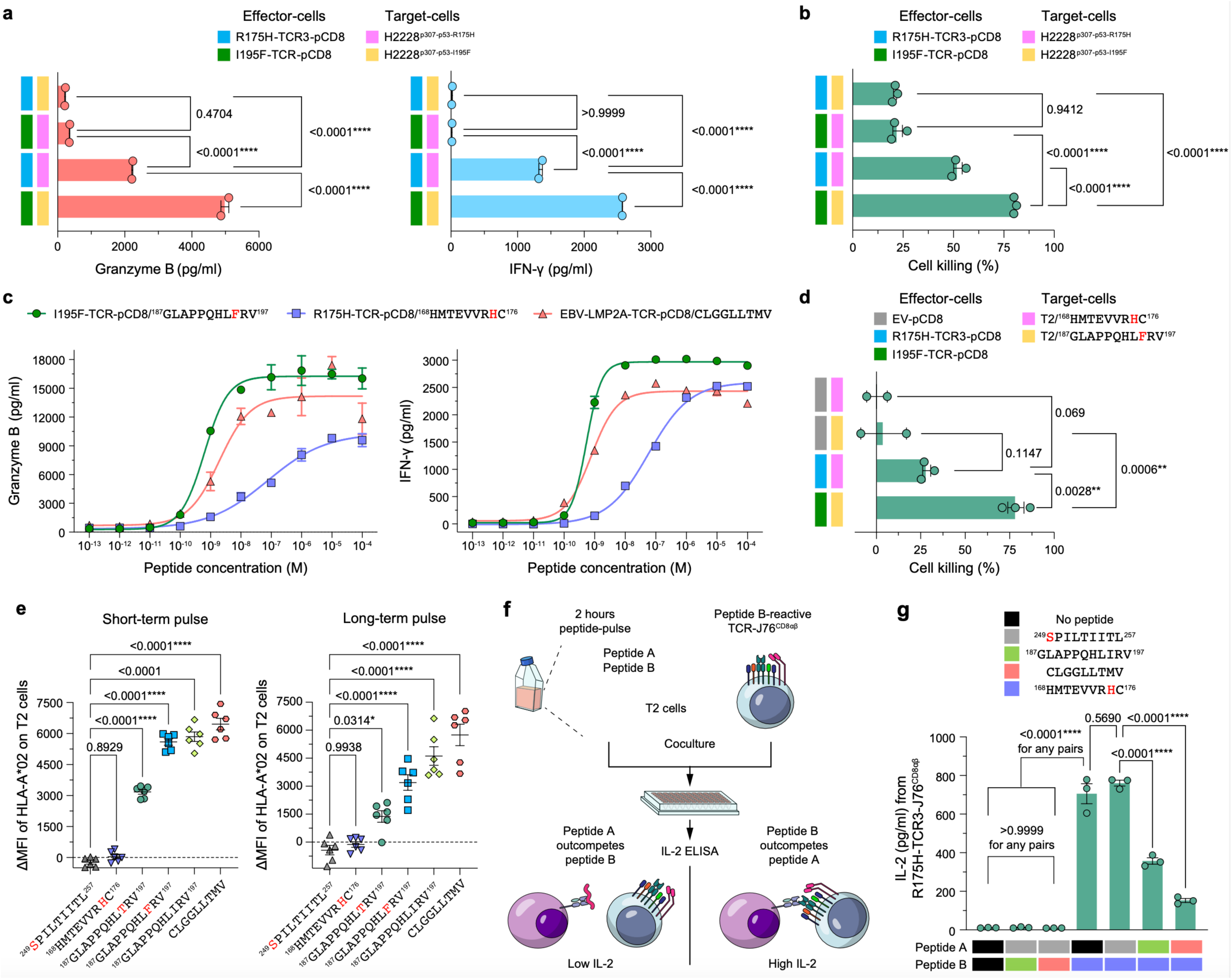
Unstable R175H/HLA-A*02:01 neoepitope expression degrades antigenicity and diminishes cancer cell susceptibility to adaptive T cell attack. **a**, Granzyme B (left) and interferon-γ (IFNγ; right) release as markers of T cell activation. Effector pCD8 T cells were cocultured with the indicated target cells for 24 h. Data are representative of two independent biological replicates. Mean values and standard error of the mean (SEM; bars) and individual values from two technical replicates (filled circles) are shown. **b**, Cell killing by p53 neoepitope**–**reactive pCD8 cells on p53-neoepitope**–**expressing H2228 cells. Effector cells and target-cells were cocultured as indicated for 24 h. Data representative of two independent biological replicates. Mean values and SEM (bars) and individual values from three technical replicates (closed circles) are shown. **c**, Dose-response curves for release of granzyme B and IFNγ by the indicated pairs of TCR-pCD8 cells and peptides pulsed T2 cells. Peptides were added at the indicated concentrations for 2 h, then cocultured with TCR-pCD8 cells for 24 h. Data representative of two independent biological replicates. Mean values and SEM (bars) and individual values from two to three technical replicates with indicated symbols are shown. **d**, Cell killing effect of p53 neoepitope**–**reactive pCD8 cells on peptide-pulsed (1 nM) T2 cells. Effector cells and target-cells were cocultured for 24 h. Data representative of two independent biological replicates. Mean values and SEM (bars) and individual values from two to three technical replicates (filled circles) are shown. **e**, HLA-A*02 expression levels on T2 cells after short-term (2 h, left) or long-term (6 h, right) pulse with indicated peptides at 10 μM. ΔMFI ([mean fluorescence intensity (MFI) of HLA-A*02 with indicated peptides] – [MFI of HLA-A*02 without any peptides]) reflects HLA-A*02:01 surface stabilisation by peptide binding. Data representative of three independent biological replicates. Mean values and SEM (bars) and individual values from six technical replicates (closed symbols) are shown. **f**, Schematic (left) and **g**, results (right) of HLA-A*02:01 competition assay for R175H-TCR3-J76^CD8αβ^ are shown. Data representative of two independent biological replicates. Mean values and SEM (bars) and individual values from three technical replicates (circles) are shown. All *P* values are calculated by one-way analysis of variance (ANOVA) with Tukey’s multiple-comparison test.

Because peptide affinity for MHC is critical in preserving surface expression of a pMHC and hence its half-life^50,51^, we compared HLA-A*02:01 binding capability between these two mutant-p53 peptides and others by T2 A*02:01 stabilization assay^51^. While ^187^GLAPPQHL<u>X</u>RV^197^ stabilised HLA-A*02:01 as well as the EBV-LMP2A positive control, ^168^HMTEVVR<u>H</u>C^176^ did not, yielding a mean fluorescence intensity (MFI) by quantitative immunofluorescence analysis equivalent to the non–A*02:01-binding 9-mer ^249^<u>S</u>PILTIITL^257^, results unmitigated by longer peptide-pulse exposure time (Fig. 6e). Thus, poor stabilization is unlikely a consequence of a slow p53R175H on-rate, instead underscoring neoepitope instability in the MHC antigen presenting groove, with dissociation on the cell surface at 37°C leading to loss of HLA-A*02:01 expression. Competition assay using empty surface MHC molecules function as a rough surrogate for peptide loading to activate their corresponding TCR-J76^CD8αβ^ confirmed this assertion (Fig. 6f,g). T cell activation by ^168^HMTEVVR<u>H</u>C^176^/A*02:01 was significantly undermined by another p53^wt^ peptide ^187^GLAPPQHLIRV^197^ as well as the unrelated high affinity EBV-derived A*02:01 binder CLGGLLTMV. Meanwhile, ^187^GLAPPQHL<u>F</u>RV^197^ outcompeted even the robust competitor CLGGLLTMV, emphasizing that cancer cells must develop mechanisms to exclude this 11-mer peptide prior to HLA-A*02:01 association (Extended Data Fig. 8g).

Importantly, we found that both R175H-TCR3 and I195F-TCR manifest high quality (digital) performance when optical tweezers are used to place physical load (piconewton [pN] level forces) on their respective TCR-pMHC bonds, as would occur during T cell immune surveillance in tissues^20^ (Extended Data Fig. 9). This force-bond lifetime analysis provides the best metric to assessment of TCR performance^20^. Such biophysical analysis confirms that both TCRs can detect as few as 1–2 copies of neoepitopes per cell. Thus, instability of the ^168^HMTEVVR<u>H</u>C^176^/A*02:01 degrades antigenicity despite neoepitope immunogenicity^43^, thereby diminishing TCR activation and cytolysis against p53^R175H^ neoantigen–expressing tumours to facilitate their escape from CTL without requiring immune editing.

## Discussion

Here we have established a pipeline that refines aspects of p53 neoantigen presentation, emphasizing the critical importance of determining physical neoepitope display. We further highlight a deceptive feature of cancer cells to suppress robust immunogenic truncal neoepitope display through divergent mechanisms during clinical tumour development. Such tumour chicanery involving immune editing that impacts truncal drivers has been understudied, perhaps because continual oncogenic driver expression fueling tumour growth (i.e., positive selection) implied a persistence of those surface arrayed neoantigens. In retrospect, given myriad strategies of immune evasion used by infectious pathogens^52^ and tumours^53^, our results are not unexpected.

To achieve the requisite attomole detection sensitivity necessary to arrive at these conclusions, our MS methodology reduced LC physical dimensions and leveraged information provided by LC-MS analyses of synthetic peptide candidates. The target recognition in our LC-DIA-MS uses a probabilistic or entropic measure of stochastic event sampling for both chromatographic traces and ion map surfaces, and, as evidenced by p53 neoepitope analysis, is favorable relative to other current metrics of detection (Supplementary Discussion 1). This detection methodology is also applicable to neoantigens generated from insertion/deletions, gene fusions, noncoding genomic regions, RNA splicing or editing, and endogenous retroviruses. With respect to single nucleotide variants (SNVs), a correspondence between p53^wt^ and p53^mut^ display is generally expected unless the point mutation alters a residue impacting HLA binding. That said, such a case has been described for an uncommon *PIK3CA* mutation^41^.

The discerning nuance of tumour editing is astounding even involving truncal antigens that differ at a single amino acid residue. For instance, R249M is a structural mimic of R249wt that foils immunogenicity while R249S in the same HLA-B*07:02 creates a self-deviant immunogenic pMHC ligand appearing different from self (Fig. 3a–c) that potently stimulates anti-tumour immunity to prevent *in vivo* clinical development (Fig. 2e). In the case of the HLA-A*02:01 restricted 11-mer, the single residue I195F markedly impacts the peptide structure (Fig. 3d,e), changing antigenicity dramatically and allowing for a potent immune response (i.e., strong immunogenicity) that clinical tumours must suppress by overexpressing ERAP1 in order to cleave ^187^GLAPPQHL<u>F</u>RV^197^ as a countermeasure to avoid CTL targeting (Fig. 5). In this context, immunopeptidomic analysis suggests that ongoing immune pressure in CD8^+^ T cell-infiltrated tumour locales diminishes MHC class I neoepitope expression relative to T cell-excluded tumour zones of the same lung cancer patients^54^.

The prevailing view is that truncal oncogenic driver neoantigens are the highest value cancer immunotherapy targets being expressed on virtually all tumour stem cell progeny in each patient. Moreover, drivers are shared by different patients with the same HLA haplotype even across multiple distinct cancer types in contrast to passenger mutations that are typically subclonal and patient-unique^5^. Not surprisingly, initial clinical trial results utilizing 20 neoantigen drivers including from p53 in a vaccination and checkpoint blockade dual therapy for advanced cancer have been disappointing^55^. Hence additional insights are required to discern correct response targeting. Elegant analysis of p53 has suggested an evolutionary trade-off between neoantigen immunogenicity and oncogenic potential^56^. In this light, immunogenicity is permitted if oncogenic potential is high. However, our results demonstrate that oncogenic driver function can be maintained when neoepitope expression is excluded from the tumour immunopeptidome via the tumour’s adaptive response (negative selection) towards CD8 T cell immune selection pressure. Furthermore, among the 50 most frequent p53 missense mutations across human cancers, all mapping to the DNA binding region^30^, none of the hotspot mutations (those accounting for more than 25% of all p53 SNVs across cancers) were detected in the immunopeptidome of the tumour cells by MS, aside from R175H defined previously^30^. Thus, almost all hotspot SNV-encoded protein segments are stealth at the start while others become so, as evidenced by I195F. Exceptions are R175H whose weak HLA-A*02:01 binding degrades its antigenicity, circumventing effective immunity even though high quality TCRs can be generated to this ligand, and R249S for which immunogenicity is so potent as to disfavor clinical tumour evolution in HLA-B*07:02 individuals overall. Only a small subset of R249S tumours can survive through an escape mechanism(s) yet to be elucidated.

Given the paucity of driver neoantigen display, how can we generate protective immunity? Detailed antigenicity and immunogenicity analyses of individual epitopes identified by MS is a worthy but arduous pursuit. Alternatively, although chemical degrader-based efforts to increase tumour antigen flux in the immunopeptidome were ineffective, other approaches to enhance immunogenic antigen display on tumours are more generic and worth consideration. Explicitly, ERAP1 inhibitors^49^, small molecules binding to HLA anchor pockets^57^, or RNA splicing modifiers^58^ can shift the immunopeptidome to reveal new specificities that have not undergone central tolerance. The consequent dramatic shifts should induce endogenous polyclonal T cells expansion, thereby making tumour immune editing challenging for a cancer to escape adaptive immunity. Such an approach may require selective delivery to tumours (for example using antibody-drug conjugates) to prevent off-tumour effects. Alternatively, adoptive TCR-based cellular immunotherapies for less robust targets such as R175H might overcome impediments of poor antigenicity (Fig. 6) if large numbers of autologous lentivirus transduced digital CD8 T cells are provided, greatly exceeding those generated during natural endogenous expansion because of unstable R175H/HLA-A*02:01 tumour expression. Additional efforts to enhance T-cell performance through hybrid TCR-like CARs integrating TCR detection sensitivity and signaling potency with single chain Fv high binding affinity against neoepitopes are worthwhile^59–61^. Diabodies directed at this pMHC are another consideration^4^. The search for other truncal antigens and their subsequent targeting efforts shall benefit from robust physical detection.

## Supporting information

Supplmentary Discussion 1

Supplmentary Discussion 2

Supplmentary Discussion 3

## Methods

### Mass spectrometry

#### Affinity capture of peptide-HLA complexes

Adherent cells were released into suspension by incubation in Non-Enzymatic Cell Dissociation Medium (NECDM; phosphate-buffered saline, pH7.2; Fetal Bovine Serum, 1%, EDTA,10 mM; EGTA, 10 mM) for 1 h at 37°C. Cells were washed in NECDM (500g, 5 mins at 4°C) and resuspended to 10^6^/mL in NECDM. For each sample, 10^6^ cells were pelleted by centrifugation and the pellet gently resuspended in 1 mL digitonin extraction buffer (digitonin, 0.15 mM; sucrose, 75 mM; sodium chloride, 25 mM; PIPES, 2.5 mM, pH6.8; EDTA, 1 mM; magnesium chloride, 0.8 mM; protease inhibitor cocktail (cØmplete; Roche)). If using tumour fragments (5–30 mg), upon addition of the digitonin buffer to the sample in Eppendorf 1.5 mL LoBind tubes, the samples were gently homogenized on ice using a micropestle for 2–3 mins. For all samples, after incubation on ice for exactly 10 mins, the permeabilized cells/tissue samples were pelleted by centrifugation (480g, 10 mins at 4°C) and resuspended in Triton X-100/alkylation buffer (Triton X-100, 0.5%; sucrose, 75 mM; sodium chloride, 25 mM; PIPES, 2.5 mM, pH7.4; EDTA, 4 mM; magnesium chloride, 0.8 mM; protease inhibitor cocktail (cØmplete, Roche); cysteines were alkylated with iodoacetamide added to 10 mM final just before use). After incubation for 30 mins on ice in the dark, the nuclei (together with stromal material for tumour samples) were pelleted by centrifugation (5000g, 10 mins at 4°C) and the clarified supernatant transferred to new tubes (1.5 mL, LoBind, Eppendorf). Further Triton X-100 was added to bring the proportion to 1%, incubated for a further 20 mins on ice and centrifuged at 21,600 g for 20 min to pellet microsomal material, subcellular organelles and remaining stromal debris. The clarified lysates were added to new 1.5 mL tubes together with Protein A-agarose beads (10 µL packed volume) and anti–HLA-A, B, C monomorphic determinant (4 µg; clone W6/32, Biolegend) followed by rotation for 3 h at 4°C. After centrifugation (300g for 2 mins at 4°C and used in all subsequent steps), the agarose bead pellet was washed x6 in octyl β-D-glucopyranoside wash buffer (1 mL; octyl β-D-glucopyranoside, 1.75%; sodium chloride, 400 mM; Tris-HCl, 40 mM, pH7.6; EDTA, 1 mM) followed by 2 washes in salt wash buffer (sodium chloride, 400 mM; Tris-HCl, 40 mM, pH7.6; EDTA, 1 mM). After removal of supernatant, pelleted beads with bound immunoprecipitate were stored at −80°C prior to peptide elution for mass spectrometry.

#### Sample preparation for LC-MS

Beads were washed with 200 μL of 5% acetonitrile (Fisher Chemical, HPLC grade) in water (LC-MS grade; Thermo Scientific) (x3) and transferred to a clean 0.5 mL tube (LoBind, Eppendorf) and all fluid aspirated leaving wet beads. 15 μL of 0.2% TFA (Optima LC/MS, Fisher Chemical) along with a set of 40 retention time peptides (JPT Peptide Technologies) at 250 attomoles/peptide were added to beads. The bead/acid mixture was set at 65°C for 5 mins and then extracted with a C18 tip (ZipTip, Millipore Sigma). The tip was washed with 0.2% TFA in water (x5) followed with 0.1% formic acid (>99%, Thermo Scientific) in water (x3). Peptides were eluted into 2–3 μL 60% MeOH (HPLC grade, Fisher Chemical) in water, and the volume was reduced under an N_2 s_tream following which 0.1% formic acid was added to form 1 μL for loading the trapping column with an He-driven pressure bomb.

#### Chromatography

Alkane-modified polystyrene-divinylbenzene monoliths in 20-and 50-micron ID silica capillaries were synthesized in house for analytical and trapping columns, respectively^62^. 90-minute segmented linear gradients from 0–40% acetonitrile in water (both solvents 0.1% formic acid) were employed with the segmentation varying somewhat depending on the column in use. Flow rates varied with columns but were under 10 nanoliters/minute.

#### DDA and DIA

A Sciex 6600+ quadrupole-oTOF was used for all experiments. For extracting reference fragmentation patterns from synthetic peptide sets (JPT Peptide Technologies) the instrument was operated in a data dependent acquisition (DDA) mode. The synthetic sets were simple, consisting of orders for 250–400 pooled peptides and were analyzed at a nominal 150 or 300 attomoles per peptide in a 0.5 or 1 μL loading. For elution mapping of the synthetic set and all sample runs the instrument was operated in a data independent acquisition (DIA) mode. The instrument collects a full range mass spectrum followed with 11 MS/MS spectra in which the quadrupole filter is set to transmit an m/z window (the width varies over the 11) such that the union of these windows cover the m/z range of interest. In this way all precursor molecular ions are fragmented, but each fragmentation pattern is embedded in a complex background of other co-selected molecular ions. The sequence of MS and 11 MS/MS collections is repeated through the LC elution.

#### MS Data Analysis

The general theory of the Poisson/relative entropy detection, the three-dimensional extension and visualization methodology and some studied applications are reviewed in Supplementary Discussion 1.

### RNA sequencing of patient-derived tumour

RNA was isolated from biopsy samples (∼25 mg tissue) using the RNeasy Mini kit (Qiagen), according to the manufacturer’s directions. Libraries were prepared for RNA-Seq analysis using the SMART-Seq v4 Ultra low Input RNA kit (Takara), followed by addition of Illumina adapters and 150 bp paired end (PE150) sequencing on the Illumina NovaSeq 6000 platform (MedGenome Inc.). The paired output fasta files for each sample were checked for quality using FastQC, and adapters trimmed using FastqMcf and Cutadapt^63^. Contaminating non-polyA-tailed RNA sequence, mitochondrial genome sequences, ribosomal RNAs and transfer RNAs were then removed using Bowtie2^64^ and the resulting filtered reads aligned to the human reference genome assembly GRCh38 using STAR (v2.7.3a)^65^. HLA display from the aligned BAM assembly file was then determined using the consensus calls from 3 tools (OptiType^66^, xHLA^67^, and HLA-LA^68^) thus minimizing implementation-specific errors.

### Whole exome sequencing of patient-derived tumour

Genomic DNA (gDNA) was isolated from ∼25 mg tumour tissue using the QIAamp DNA Mini kit with initial tissue disruption in ATL/proteinase K buffer using a micropestle in Eppendorf Lo-Bind 1.5 mL tubes. Following complete tissue dissolution at 56°C for 2 h, viscous DNA was sheared into uniform 23 kb fragments by passage through a QIAshredder column and centrifugation for 2 min at 15,000 g. The manufacturer’s protocol for gDNA purification was subsequently followed for all steps. Libraries were prepared from the purified gDNA at MedGenome for Whole Exome Sequencing using the Agilent SureSelect Human All Exon v7 kit and the manufacturer’s protocol. Quality control for the library included Qubit fluorometric quantification, Agilent 4200 tapestation electropherogram, and qPCR. Following sequencing (PE150) on the NovaSeq platform to a depth of x300, the output fastq files were examined for quality control by checking base quality, nucleotide distribution, %GC and PCR duplicates. Preprocessing included read trimming using Cutadapt ^63^ ^w^here appropriate. Following alignment of reads to the GRCh38 human reference genome using Bowtie^64^, genomic variants were called using GATK Haplotypecaller^69^ and output in VCF file format. Data was visualised and manipulated for genomic data exploration using Golden Helix GenomeBrowse (Golden Helix, Nagoya, Japan).

### HLA-A*02:01 and HLA-B*07:02 complexes: expression, refolding, and purification

Peptide/MHC complexes were generated as previously published for pMHC production^70,71^, and peptides were chemically synthesized (United Biosystems, Inc). Briefly, HLA-A*02:01, codon optimized HLA-B*07:02 and human β2m (hβ2m) were expressed separately using BL21-DE3 cells transduced with the HLA or β2m-encoded pET vector. HLA-A*02:01 and hβ2m were expressed in 1 L LB medium at 37°C to OD_600 =_ 0.6 prior to induction with 0.5 mM IPTG for 3 h after induction. HLA-B*07:02 was expressed in 1L MagicMedia^TM^ (Invitrogen) by autoinduction overnight at 37°C according to manufacturer recommendations. Cells were harvested by centrifugation, resuspended in 30 mL Tris buffered saline (TBS) pH 8.0 and lysed by incubation with 1 mg/mL lysozyme, 0.3 mg/mL DNAase I, and protease inhibitor (Roche) and repeated freeze/thaw. Inclusion bodies were purified by centrifugation/resuspension using two washes of TBS, two of TBS + 1% TritonX-100 and one of TBS. Purified inclusion bodies were dissolved in 10 mL 5.4 M Guanidine HCl + 0.1 M Tris-HCl pH 8.0 + 2 mM dithiothreitol and stored in aliquots at –80°C. To refold pMHC, HLA-A*02:01 or HLA-B*07:02, hβ2m and peptide were diluted at a ratio of 60:20:10 w/w into 300 mL 8 M Urea + 20 mM Tris-HCl (final concentration ∼0.25 mg/mL) and sequentially dialyzed against 4 L of 2, 1, 0.5, and 0 M Urea + 20 mM Tris-HCl serially with each step greater than 2 h or overnight for a total of at least 48 hours dialysis/refold time. Proteins were purified to homogeneity by size exclusion chromatography followed by anion exchange chromatography, with the final product eluting as a single peak and two bands present (heavy chain and β2m) when separated by non-reducing SDS-PAGE. The protein was then buffer exchanged into 10 mM Tris-HCl, pH 8.0 + 100 mM NaCl and concentrated to approximately 10 mg/mL (I195 WT/ HLA-A*02:01) or 20 mg/mL (I195F/ HLA-A*02:01 and all HLA-B*07:02 complexes) for crystallization trials.

### Crystallization, Data collection and Structural determination

The crystallization trials of WTp53_^187^GLAPPQHLIRV^197^/HLA-A*02:01 (10.3 mg/ml), p53_I195F/HLA-A*02:01(19.6mg/ml), WTp53_^249^RPILTIITL^257^/HLA-B*07:02 (20.8mg/ml), p53_R249M/HLA-B*07:02 (17.6mg/ml) and p53_R249S/HLA-B*07:02 (17.5mg/ml) were set up with a Mosquito nanoliter liquid handler (TTP LabTech) using the sitting drop vapor diffusion technique in 96-well CrystalQuick plates (Greiner). For each condition, 0.4 ml of pMHC sample and 0.4 ml of crystallization formulation were mixed; the mixture was equilibrated against 140 ml of precipitating solution in each reservoir well. The crystallization screens used were MCSG-1–4 (Anatrace) at 16°C. Crystals of both WTp53_^187^GLAPPQHLIRV^197^/HLA-A*02:01 and p53_I195F/HLA-A*02:01 appeared under the same condition of 0.1 M Bis-Tris Propane:HCl pH 7 and 60% (v/v) Microlytic Mix. Crystals of WTp53_^249^RPILTIITL^257^/HLA-B*07:02, p53_R249M/HLA-B*07:02 and p53_R249S/HLA-B*07:02 all appeared under the same condition of 0.2 M ammonium sulfate, 0.1 M Tris:HCl pH8.5, 25% (w/v) PEG 3350. The crystallization of p53_R249S/HLA-B*07:02 was further improved by using additive screening (Hampton Research). The diffraction quality crystals of p53_R249S/HLA-B*07:02 were obtained from the buffer that contains 10% ZnCl_2 a_nd 90% initial crystallization screening buffer mentioned above. All of these crystals were harvested and treated with a cryoprotectant solution (25% glycerol or 25% ethylene glycol in mother liquor) and then flash-frozen in liquid nitrogen before X-ray diffraction data collection.

X-ray diffraction data were collected at 100 K from the cryocooled crystals at the 19-ID (NYX) beamline or 17-ID-2 (FMX) beamline of National Synchrotron Light Source II at Brookhaven National Laboratory. The intensities of each data set were integrated, scaled, and merged with the HKL-3000 program suite^72^. The structure of p53_^187^GLAPPQHLIRV^197^WT/HLA-A*02:01 was first determined using the molecular replacement (MR) method^73^ with the HLA-A*02:01 structure (PDB code: 5D9S) as a search template. There are two p53_WT/HLA-A*02:01 complexes in one asymmetric unit. The pMHC complex packs in a honey-comb-like fashion within crystals, resulting an unusually high solvent content of ∼82%. The model rebuild, including building the peptide was performed using the program Coot ^74^. The final model was refined using the program phenix.refine^75^. The structure of p53_I195F/HLA-A*02:01 was subsequently determined and refined in the same procedure. The structural determination of p53_^249^RPILTIITL^257^WT/HLA-B*07:02 was similarly done by using MR method with a known HLA-B*07:02 structure (PDB code: 4U1H) as a search template. Structural determination of p53_R249M/HLA-B*07:02 and p53_R249S/HLA-B*07:02 followed. In the presence of Zn^2+^ ion in the additive screening condition, p53_R249S/HLA-B*07:02 formed a different crystal form, in which Zn^2+^ ions bridge neighboring molecules and presumably stabilised molecular packing.

### Culture of cell lines and patient-derived organoids

NCI-H2228 (CRL-5935), CaSki (CRL-1550), HEK293T (CRL-3216), TAP1/2-deficient T2 (CRL-1992), KLE (CRL-1622), and CT26 (CRL-2638) cells were obtained from the American Type Culture Collection. Lenti-X 293T (Cat.# 632180) cells were obtained from Takara Bio Inc. COV-318 (Cat.# 07071903) cells were obtained from Sigma-Aldrich. TCR-null/CD8-deficient Jurkat-76 cells were kindly gifted by Dr. Mirjam H.M. Heemskerk (Leiden University Medical Center, Netherlands)^45^ under a material transfer agreement. PANFR-0583 patient-derived organoids were established from a liver metastasis legion of pancreatic ductal adenocarcinoma as previously described^40^. Cells were maintained in a humidified atmosphere of 5% CO_2 a_t 37°C. NCI-H2228, CaSki, T2, Jurkat-76, and CT26 cells were cultured in RPMI-1640 medium (11875-119, Gibco). HEK293T, Lenti-X 293T, and COV318 cells were cultured in DMEM (11965118, Gibco; 10-013-CV, Corning). KLE cells were cultured in DMEM/F-12 (11320033, Gibco). All medium was supplemented with 10% fetal bovine serum (FBS) (100-106, GeminiBio products) and 1% penicillin-streptomycin (10378016, Gibco). For passage of adherent cells, cells were washed once by phosphate buffer saline (PBS) (10010023, Gibco), and then collected by Trypsin-EDTA (25200114, Gibco). PANFR-0583 cells were cultured and handled as previously described^40^. Cells were regularly confirmed negative for mycoplasma contamination using the Universal Mycoplasma detection kit (30-1012K, ATCC). Cells were cryopreserved in Bambanker serum-free cryopreservation media (50-999-554, Fisher Scientific) as necessary. Passage numbers were limited 10 times or less throughout this research project with fresh cultures established from frozen stocks as required.

### Isolation of primary human peripheral blood mononuclear cells

De-identified human healthy donors–derived whole blood byproducts of platelet donation were obtained from the Crimson Core Biomaterials Collection Core in Brigham and Women’s Hospital (Boston, USA), and then transferred into BD Vacutainer CPT Mononuclear Cell Preparation Tubes (362760, BD) to isolate peripheral blood mononuclear cells (PBMC). After centrifugation at 1,500 g for 20 mins, PBMC in buffy coat were collected and washed x2 with PBS at 300 g for 10 mins before use. Excess PBMC were frozen in CELLBANKER 2 serum-free freezing media (11891, amsbio). Any primary cells derived from human PBMC were cultured in human T cell medium consisting of RPMI-1640 medium, 10% heat-inactivated human serum AB (Sigma-Aldrich, H5667), 2 mM L-glutamine (25030149, Gibco), and 1% penicillin-streptomycin. DNA/RNA sequencing was not permitted for PBMC obtained from the Crimson Core. Consequently, and untyped PBMC subsequently typed in this study were purchased from STEMCELL Technologies (Cat.# 70025). In each comparison, PBMC were obtained from the same donors.

### Preparation and culture of primary human T cells for LC-MS

Frozen HLA-A*02:01^+^ HLA-B*07:02^+^ PBMC from the donor RG2684 (70025, STEMCELL Technologies) were thawed, washed x2 in human T cell medium (at 400 g for 6 mins), and with human T cell medium for 45 mins at 37°C incubator. T cells were then isolated from the PBMC using the EasySep Human T Cell Isolation Kit (17951, STEMCELL Technologies) with EasySep Buffer (20144, STEMCELL Technologies). The isolated T cells were activated using human T Cell TransAct (130-111-160, Miltenyi Biotec) at 1:100 in the T cell medium (1 x 10^6^ cells/mL) with hIL-2 (100 IU/mL; PHC0021, Gibco), hIL-7 (25 ng/mL; PHC0075, Gibco), and hIL-15 (25 ng/mL; PHC9154, Gibco). Human T cell medium and cytokines were refreshed every three days. After 6 d (144 h) of activation, the cells were processed for pHLA immunoprecipitation for LC-MS as described above, for protein extraction for western immunoblotting, for RNA extraction, and for qRT-PCR.

### *In vitro* cell line treatment with chemical reagents

dTAG47 (HY-147098, MedChemExpress), TL-13-112 (HY-123919, MedChemExpress), and ERAP1-IC3 (HY-133125, MedChemExpress) were dissolved in DMSO, according to the manufacturer’s instructions. Treatments with dTAG47 and TL-13-112 were performed as indicated in the figure legends. For ERAP1-IC3 treatment, the reagent was added to culture medium at 50 μM overnight and was continued during subsequent coculture with T cells at the same final concentration. DMSO was used as a negative control at concentrations equivalent to corresponding treatment in all experiments.

### Cloning of neoepitope-reactive human CD8^+^ T cell clonotypes

Frozen HLA-A*02:01 PBMC (70025, STEMCELL Technologies) were thawed, washed x2 using the human T cell medium (at 400g for 6 mins), and incubated in 15 mL polypropylene tubes with the T cell medium diluted with EasySep buffer at 1:1 ratio for 45 mins at 37°C. DNase (07900, STEMCELL Technologies) was added to the medium at 100 μg/mL. These PBMC were incubated in 6-well plates (3516, Corning) at 1 x 10^7^ cells per well with 2 mL of T cell medium for 2 h allowing monocytes to adhere to the plate (∼1 x 10^6^ cells/well). Monocytes were differentiated into dendritic cells (DCs) in T cell medium supplemented with hIL-4 (200 IU/mL; 200-04, Peptotech) and granulocyte-macrophage colony-stimulating factor (GM-CSF) (800 IU/mL; 300-03, Peprotech) for 6 d (from day −6 to day 0), during which the medium was also supplemented with IFNγ (100 IU/mL; 300-02, Peprotech) and lipopolysaccharides (10 ng/mL; L4391-1MG-PW, Sigma-Aldrich) for the last 16–24 hours. Subsequently, ^187^GLAPPQHLIRV^197^, ^187^GLAPPQHL**<u>T</u>**RV^197^, or ^187^GLAPPQHL**<u>F</u>**RV^197^ peptides (MBL International) were pulsed individually onto the induced dendritic cells for an additional 2 h (1 μM). At this point, additional frozen PBMC from the same donor were processed in the same manner to provide the second batch of mature DCs at d 6. Naïve CD8^+^ T cells were also isolated from the same PBMC using EasySep Human Naïve CD8^+^ T Cell Isolation Kit (19258, STEMCELL Technologies) and EasySep Buffer. After washing x2 the peptide-pulsed DCs with PBS (3 mL), 7.0 x 10^6^ isolated naïve CD8^+^ T cells were added to the DCs at 3.5 x 10^6^ cells/well in T cell medium along with hIL-2 (100 IU/mL), hIL-7 (25 ng/mL), and hIL-15 (25 ng/mL). At d 6, the naïve CD8^+^ T cells were combined into one tube for each peptide condition, and then split into 2 wells of a new 6-well plate already seeded with the second batch of peptide-pulsed DCs. Culture was continued until d 15 with T cell medium and cytokines refreshed every three days. During the 15 days, T cell medium and cytokines were refreshed every three days. At d 15, cocultured naïve CD8^+^ T cells were collected from one well for each peptide condition, and then stained as described below with Zombie-NIR (423106, BioLegend), FITC–anti-CD3ε (11-0036-42, Invitrogen), BV510–anti-CD8a (563919, BD Biosciences), and PE– or APC– peptide-MHC-dextramers (Immudex, Virum, Denmark) to determine neoantigen-reactive T cell expansion by flow cytometry (LSRFortessa X-20, BD Biosciences).

At d 17, all non-adherent cells were collected for isolation of neoepitope-reactive T cells. Cells were stained as described below with Zombie-NIR, FITC–anti-CD3ε, BV510–anti-CD8a, and PE– or APC– peptide-MHC-dextramers. All double-colour-positive dextramer-positive viable T cells were isolated on a FACSAria II Flow Cytometer (BD Biosciences). Non-expanded T cells were used as a negative control to ensure true positive cells.

Isolated dextramers-positive T cells were washed with the cold T cell medium once, and resuspended to 1,000 cells/μL for single-cell RNA sequencing, aiming for a properly-sequenced final count of ∼10,000 cells per condition. The same number of non-expanded autologous T cells derived from the wells loaded with ^187^GLAPPQHLIRV^197^ were also subjected to the same pipeline as a negative control. Low isolated cell numbers were supplemented with irrelevant non-immune cells as a carrier. Cells were were loaded onto a Chromium instrument (10X Genomics) for processing to barcoded single-cell cDNA libraries using the Chromium Next GEM Single Cell 5ʹ Kit (PN-1000263, 10X Genomics), and the Chromium Single Cell Human TCR Amplification kit (PN-1000252, 10X Genomics). Quality control of the cDNA libraries was performed using the Bioanalyzer High Sensitivity DNA Kit (5067–4626, Agilent Technologies). Sequencing using the Illumina NovaSeq platform, and conversion of resultant base call files into fastq files were performed by Novogene Co, Ltd.

The raw fastq files were processed by CellRangerCount 7.0.1 and CellRanger VDJ 7.0.1 using the human reference GRCh38 for alignment and assembly to generate filtered gene-barcode matrix files and vdjloupe files. Filtered gene-barcode matrix files were analyzed by Seurat pipeline (v.5.0.3)^76^. Seurat objects for ^187^GLAPPQHL**<u>F</u>**RV^197^/A*02:01-dextramers–sorted T cells and the negative control were merged into a single object for downstream analysis. Cells which were containing fewer than 250 unique molecular identifiers (UMIs), expressing fewer than 250 genes, or with more than 5% of mitochondrial gene content were excluded as debris or dying cells. Cells were filtered out as doublets if the number of UMIs were ≥20,000 or the number of expressed genes were ≥4,500. Remaining UMI counts were normalised and log_2-_transformed by a global-scaling method. 2,000 variable features were then found using ‘FindVariableFeatures’ with ‘vst’ option. After scaling all data, principal component analysis (PCA) was performed with the 2,000 variable features for linear dimensional reduction. Based on JackStraw plots, ElbowPlots, and heatmaps, the top 10 principal components were used for Uniform Manifold Approximation and Projection (UMAP) and clustering. Clusters positive for *CD3E* but negative for *EPCAM* and *EGFR* were extracted as a pure T cell subset for subsequent analyses. PCA was reperformed in the same manner, and the top 15 principal components were used for a final UMAP visualization. To obtain consensus VDJ sequences, the vdjloupe files were processed by Loupe VDJ Browser 5. Whereas fastq files provided by Chromium Single Cell Human TCR Amplification kit were not capable of creating vdjloupe files due to insufficient cDNA library quality, durable vdjloupe files were successfully recovered with fastq files using the Chromium Next GEM Single Cell 5ʹ Kit (Supplementary Data File 8).

### Plasmid construction

DNA fragments encoding p53^wt^, p53^mut^, HLA-B*07:02, firefly luciferase, and CD8αβ were synthetized using the gBlocks platform (Integrated DNA Technologies). These DNA fragments were designed for assembly using the Gateway system (ThermoFisher) using a 5’ attB1 sequence followed by a Kozak sequence and a 3’ attB2 sequence. The DNA fragment for CD8αβ was designed as a single linear fragment encoding CD8β, a cleavable furin-GSG-P2A linker, followed by CD8α isoform M4. The inserts were introduced into destination lentivirus vectors including pLX304 (25890, Addgene), pLEX_305-C-dTAG (91798, Addgene), or pLEX_307 (41392, Addgene) through pDONR223 (no longer available, originally from Invitrogen) using the Gateway Cloning system with LR Clonase (11791-020, Invitrogen) and BP Clonase (11789-020, Invitrogen). DNA fragments encoding single-guide RNAs (sgRNAs) targeting *TP53*, *ERAP1*, and *Erap1* were synthesized for LentiCRISPRv2 system^77^ by Integrated DNA Technologies Inc. The sgRNA sequences were determined for SpyoCas9 by CRISPick (https://portals.broadinstitute.org/gppx/crispick/public). A non-targeting sgRNA from the Gecko library v2 was used as a scrambled sgRNA as previously described^77^. These sgRNA-encoding DNA fragments were introduced into lentiCRISPR v2 (52961, Addgene) or lentiCRISPR v2-Blast (83480, Addgene) backbone plasmids. Plasmids were amplified by One Shot Stbl3 Chemically Competent E. coli (C737303, Invitrogen) with relevant antibiotics following manufacturer’s instructions. Amplified plasmids were purified using the HiSpeed Plasmid Midi Kit (12643, Qiagen). All TCR plasmids were built by VectorBuilder Inc, where DNA sequences encoding TCRs following an EF1α promoter region and a Kozak sequence were put into the same lenti-virus backbone plasmids. VDJ-coding DNA sequences for R175H-TCRs were determined based on amino acid sequences previously published. cDNA information about EBV-LMP2A-TCR was kindly provided by Dr. Nicholas R.J. Gascoigne (National University of Singapore)^78^. For I195F-TCR, VDJ-coding DNA was obtained by our experiments as described above. In TCR-encoding DNA, β chain was followed by α chain via a cleavable linker furin-RAKR-SGSG-P2A, and constant region domains were replaced by mouse homologues (murine *Trac* and murine *Trbc1*) with a hydrophobic modification in *Trac* as well as cysteine replacement of the 47^th^ residue in *Trac* and the 57^th^ residue in *Trbc1*. Final plasmid sequences were verified by Sanger sequencing (Azenta US) or full plasmid sequencing (Primordium Labs). Information about the coding DNA sequences and the guide RNA sequences are provided in Supplementary Data File 11

### Preparation of lentivirus supernatant for non-TCR transduction

HEK293T cells (4 x 10^6^) were plated in 60-mm dish 1 d prior to transfection. For transfection, the HEK293T were incubated with 3 mL of complete medium with 300 μl of Opti-MEM Reduced Serum Media (31-985-070, Gibco) containing 9 μl of X-tremeGENE 9 DNA Transfection Reagent (6365779001, Roche), together with 1 μg of the lentiviral plasmids of interest, 1 μg of pCMV-VSV-G^79^, and 1 μg of pCMV-dR8.91^79^ for 12–24 hours, after which the medium was refreshed with 3 ml of fresh complete medium. After 48 and 72 hours of the transfection, the lentiviral particle–containing supernatant were collected and filtered through 0.45 μm surfactant-free cellulose acetate membrane filters (431220, Corning). The lentivirus supernatant was concentrated by Lenti-X Concentrator (631232, Takara Bio) as needed.

### Preparation of lentivirus supernatant for TCR transduction

Lenti-X 293T cells (1.5 x 10^6^) were plated in 150 mm dishes (229651, CELLTREAT Scientific Products). At ∼90% confluence (usually after 5 d), the medium was replaced with 20 mL of fresh complete medium. Cells were then incubated with 2.4mL of Opti-MEM Reduced Serum Media containing 72 μL of X-tremeGENE 9 DNA Transfection Reagent, together with 12 μg of the lentiviral TCR plasmids, 9 μg of psPAX2 (12260, Addgene), and 3 μg of pMD2.G (12259, Addgene) for 6–8 h, after which the medium was refreshed with 15 mL of fresh complete medium containing ViralBoost Reagent (VB100, ALSTEM cell advances) at 1:500. After 48 h of the transfection, the lentiviral particle–containing supernatant was collected and stored at 4°C, and a further 15 mL of fresh complete medium was added to the transduced cells. At 72 h, supernatant was collected. Collected viral particle–containing supernatant (30 mL in total) was filtered through 0.45 μm surfactant-free cellulose acetate membrane filters, and then concentrated by Lenti-X Concentrator into Opti-MEM Reduced Serum Media or human T cell medium. For empty-vector transduction as a negative control, all procedures were performed consistently in the same manner, except that the lentiviral TCR plasmids were not used.

### Lentiviral transduction of cell lines

For transduction of adherent cancer cell lines, cells were plated into 6-well plates at 2 x 10^5^ cells/well with 2 mL of complete medium. The next day, the complete medium was replaced with 1 mL of a fresh complete medium. For transduction of non-adherent cancer cell lines, cells were plated into 6-well plates at 2 x 10^5^ cells/well with 1 mL of complete medium. 20–1,000 μL of the non-concentrated or concentrated lentiviral supernatant were added to each well. Spinfection was performed at 2,000 g for 1–2 h at 32°C if required. After 12 h, the cells were washed with the complete medium once, and cultured with the fresh complete medium for a further 24 h. Selection of transduced cells was achieved by addition of 7 d treatment with puromycin (Gibco, A1113803) at 0.2–1.8 μg/mL or 10 d treatment with blasticidin (Gibco, A1113903) at 2–18 μg/mL in the complete medium with fresh antibiotic-containing media added every 3–4 days. The antibiotic concentrations used for selection were previously calibrated to kill non-transduced cells within 3 d.

### Lentiviral transduction of primary T cells with TCR

Primary human T cells were isolated from fresh PBMCs derived from the Crimson Core using the EasySep Human T Cell Isolation Kit with EasySep Buffer. Isolated T cells were activated by human T Cell TransAct at 1:100 in the T cell medium (1 x 10^6^ cells/mL) with hIL-2 (100 IU/mL), hIL-7 (25 ng/mL), and hIL-15 (25 ng/mL). After 36–48 h activation, non–tissue-culture–treated 24-well plates (3738, Corning) were precoated by RetroNectin Recombinant Human Fibronectin Fragment (T100B, Takara Bio) at 18 μg/cm^2^ following manufacturer’s instruction. The concentrated lentivirus particle solution for TCRs of interest were loaded into the precoated plates at 1,500 μL/well, and then was spun down at 2,000 g at 32°C for 2 h. After carefully removing the supernatant, activated T cells were transferred into the virus-coated 24-well plates at 5 x 10^5^ cells/well in 400 μL of the T cell medium containing the same cytokine cocktail containing LentiBOOST (4 μl; SB-P-LV-101-10, Mayflower Bioscience). The plate was spun down at 800 g, 32°C, for 10 mins. A further 500 μL of the concentrated lentivirus particle solution was added with 24 h later containing LentiBOOST (15 μL). Human T cell medium (600 μL) was added to adjust the IL-2 concentration to 100 IU/mL and IL-7/15 concentrations to 25 ng/mL. The plate was spun down at 2,000 g, 32°C, for 2 h. TCR expression level was confirmed at d 8 by flow cytometry. At day 15, CD8^+^ cells were isolated for subsequent functional assay using the EasySep Human T Cell Isolation Kit. Fresh hIL-2/7/15 was added to the cells every 3 days throughout the experiments.

### *HLA* genotyping

Cells were washed with PBS once (500g, 5 mins), and genomic DNA was extracted using DNeasy Blood & Tissue Kit (69504, Qiagen). DNA quantity was determined by NanoDrop (ND2000, Thermo Scientific), and 2 μg of the genomic DNA was submitted to HistoGenetics LLC (https://www.histogenetics.com/) for 4x high-resolution HLA genotyping. HLA haplotypes were determined by HistoGenetics LLC (https://www.histogenetics.com/) based on IMGT/HLA version 3.48.0.

### *TP53* and *ERAP1* genotyping

Cells were washed in PBS once (500g, 5 mins), and genomic DNA was extracted using DNeasy Blood & Tissue Kit. DNA quantity was determined by NanoDrop. PCR was performed using Q5 High-Fidelity DNA Polymerase kit (M0491S, New England Biolabs) according to the manufacturer’s instructions. The PCR amplicon was visualised by agarose gel electrophoresis, and then purified by NucleoSpin Gel and PCR Clean-up kit (740609.50, Takara Bio). Purified PCR amplicon was sequenced by Sanger sequencing (Azenta US), and visualised by SnapGene Viewer (version 7.0.3). The sequences of the primers are listed in Supplementary Data File 11.

### Quantitative RT-PCR

Cells were washed in PBS once (500g, 5 mins), and RNA was extracted using RNeasy Mini Kit (74104, Qiagen) according to the manufacturer’s instructions. RNA quantity was determined by NanoDrop. First-strand cDNA was generated using SuperScript III First-Strand Synthesis SuperMix for qRT-PCR kit (11752250, Invitrogen). Quantitative real-time PCR was performed using Power SYBR Green PCR Master Mix kit (4367659, Applied Biosystems) on a CFX96 Touch Real-Time PCR Detection System instrument (Bio-Rad Laboratories) following the manufacturer’s instructions. Human *36B4(RPLP0)* was used as a house keeping gene. Results were reported as 2^−ΔΔCt^. The sequences of the primers used for qRT-PCR are listed in Supplementary Data File 11.

### Immunoblotting

Cells were washed in PBS once (500g, 5 mins), and resuspended in the protein lysis buffer which consisted of RIPA Lysis and Extraction Buffer (89901, Thermo Scientific), phosphatase inhibitors (4906845001, Sigma-Aldrich), and protease inhibitors (11836170001, Roche), according to the manufacturer’s instructions. To facilitate protein extraction, cell lysates were sonicated using a Branson Digital Sonifier (101-135-066R, Branson Ultrasonics) on ice (10%, 3 s, x3). Thereafter, cell lysates were spun down at 14,000 g for 10 mins at 4°C, and protein concentration in the clarified supernatant was determined by Pierce BCA Protein Assay Kits (23225, Thermo Scientific) using an Infinite M Plex instrument (30190085, Tecan). Reducing agent (NP0009, Invitrogen) and Sample Loading Buffer (NP0007, Invitrogen) were added to the protein solution following the manufacturer’s instructions, and protein concentrations were then normalised between samples using the protein lysis buffer. After heating the mixtures at 95°C for 5 mins, samples were cooled down on ice and loaded into NuPAGE Bis-Tris Mini Protein Gels (Invitrogen) for sodium dodecyl sulfate (SDS)-polyacrylamide gel electrophoresis with SDS running buffer (NP0001, Invitrogen). Gel-separated proteins were transferred onto nitrocellulose membranes (IB23001, Invitrogen) using the iBlot2 instrument (IB21001, Invitrogen). Membranes were blocked for 1 h at RT in Odyssey blocking buffer (927-60001, LI-COR Biotech), followed by overnight incubation at 4°C with primary antibodies for the indicated proteins. Membranes were then washed x4 for 10 mins in washing buffer containing 0.05% Tween-20 (IBB-181X, Boston BioProducts), and subsequently incubated with fluorescent secodary antibodies for 1 h at RT. Following further washes x4 in the same washing buffer in the same manner, protein bands were visualised using the LI-COR Odyssey 9120 system instrument (LI-COR Biotech). Primary and Secondary antibodies were diluted in Signal Enhancer HIKARI Solution 1 and 2 (NU00102-1 and NU00102-2, Nakarai USA), respectively. The following antibodies were used: mouse–anti-p53 (1:1000, sc-126, Santa Cruz Biotechnology), mouse–anti-HSP90 (1:1000, sc-13119, Santa Cruz Biotechnology), mouse–anti–β-actin (1:2500, 3700, Cell Signaling Technology), rabbit–anti-ERAP1 (1:2500, 99979, Cell Signaling Technology), rabbit–anti-ARTS1/ERAP1 antibody (1:1000, EPR6069, Abcam), IRDye 680RD Goat anti–mouse IgG secondary antibody (926-68070, LI-COR Biotech), and IRDye 800CW Goat anti–rabbit IgG secondary antibody (926-32211, LI-COR Biotech).

### Flow cytometry

Cells were transferred to a 96-well round-bottom plate (3799, Corning) and washed with 200 μL of PBS once (450g, 6 mins), followed by incubation in PBS with Zombie-NIR (423106, BioLegend), Zombie-Aqua (423102, BioLegend), or Zombie-Green (423112, BioLegend) at 0.1:50 at RT for 5 mins. The plates were then spun down at 450 g for 6 mins at 4°C, and supernatant were discarded. After incubation of the cells with human Fc Blocker (422302, BioLegend) in FBS-containing Stain Buffer (554656, BD Pharmingen) at 1–2.5:100 (1:100 for non-immune cells, and 2.5:100 for immune cells) for 10 mins at 4°C, supernatant was removed following centrifugation (450 g, 6 mins). Thereafter, the cells were stained by the indicated antibodies or markers diluted in FBS-containing Stain Buffer for 30 mins at 4°C, and then washed twice with FBS-containing Stain Buffer prior to flow cytometry analyses. The following antibodies or markers were used: BV510–anti–human CD8a (1:100, 301048, BioLegend; 1:100, 563919, BD Biosciences), PE–anti–human CD8a (1:100, 301007, BioLegend), AF488–anti–human CD3ε (1.5:100, 344810, BioLegend), FITC–anti–human CD3ε (1.5:100, 11-0036-42, Invitrogen), BV605–anti–human CD4 (1:100, 317437, BioLegend), PE/Dazzle-594–anti–human CD137/4-1BB (1.5:100, 309825, BioLegend), APC–anti–mouse TCRβ constant region (1.5:100, 109212, BioLegend), PerCP/Cy5.5–anti–mouse TCRβ constant region (1.5:100, 109227, BioLegend), BV510–anti–HLA-ABC (2.5:100, 311436, BioLegend), APC–anti–HLA-ABC (2.5:100, 311409, BioLegend), AF700– anti–HLA-A2 (2.5:100, 343318, BioLegend), PE–anti–HLA-B7 (1:100, 372403, BioLegend), BV-421–anti–mouse H-2k^d^ (1:100, 116623, BioLegend), PE–anti–mouse H-2D^d^ (1:100, 110607, BioLegend), PE–peptide-MHC-dextramers (5:100, Immudex), and APC–peptide-MHC-dextramers (5:100, Immudex). For peptide-MHC-dextramer staining, cells were preincubated with the dextramers in FBS-containing Stain Buffer at 5:50 for 10 mins at RT. Flow cytometry was performed on an LSRFortessa X-20 instrument (BD Biosciences) or FACSAria II Flow Cytometer (BD Biosciences). Spectral compensation was performed in the flow cytometer machines using UltraComp eBeads (01-2222-42, Invitrogen) or human PBMC stained with a signle fluorophore. Fluorescence minus one (FMO) controls and corresponding isotype controls were used as negative controls. Obtained data were analysed and visualised using FlowJo software (v10.10.0).

### Single-cell clone expansion of TCR-Jurkat-76^CD8αβ^

After the lentiviral transduction of TCR-null/CD8-deficient Jurkat-76 cells with TCRs of interest and CD8αβ, the cells were stained by Zombie-Aqua, PE–anti-CD8a, and APC–anti-mouse-TCRcβ as described above. Viable human CD8- and mouse-TCRcβ–expressing singlet cells were sorted using a FACSAria II Flow Cytometer (BD Biosciences), and placed into 96-well round-bottom plates at one cell/well with 250 μL of complete medium. After several weeks, cell surface expression of CD8 and TCRs on expanded single-cell clones were confirmed. For a functional comparison of different TCRs, expression levels of CD8 and TCR were matched between single-cell clones.

### T2 peptide-pulse assay

TAP1/2-deficient T2 cells were incubated with indicated peptides (MBL International) and concentrations for 2 or 6 h at 1 x 10^6^ cells/mL of complete medium. The cells were then washed with complete medium once (450g for 6 mins at 4°C). Thereafter, peptide-pulsed cells were used for HLA-A*02 flow cytometry and coculture with TCR-pCD8 cells for ELISA or cell killing assays.

### ELISA of cytokines from TCR-pCD8 cells after coculture with target-cells

Target cells (1 x 10^5^ cells in 100 μL complete medium per well) were incubated overnight in 96-well flat-bottom plates (3596, Corning). The next day, the medium was replaced with fresh human T cell medium. In case of peptide-pulsed T2 cells, 1 x 10^4^ cells were seeded into 96-well flat-bottom plates with human T cell medium on the day of coculture. Subsequently, TCR-pCD8 cells (1 x 10^4^ per well) were added to the plates with human T cell medium. The final number of required pCD8 cells were determined by mouse-TCRcβ positivity ([1 x 10^4^] x 100/[mouse-TCRcβ positivity (%)] of the total pCD8 cells). The number of empty-vector–transduced pCD8 cells, used as a negative control, was matched to the total number of pCD8 cells, leading to final volume of 250 μL per well. After 24 h of coculture, the plate was spun down at 450 g for 6 mins at 4°C, and the supernatant was carefully collected into new 96-well round-bottom plates. The plate was then spun down at 4000 g for 10 mins at 4°C, and the supernatant was transferred into new 96-well round-bottom plates. Thereafter, supernatant cytokines were quantified on an Infinite M Plex instrument using Human Granzyme B or IFN-gamma ELISA Kit (DGZB0 or SIF-50C, R&D) according to the manufacturer’s instructions.

### ELISA of cytokines from TCR-Jurkat-76^CD8αβ^ cells after coculture with peptide-pulsed T2 cells

1 x 10^5^ peptide-pulsed T2 cells and 1 x 10^5^ TCR-J76^CD8αβ^ cells were placed in each well of a 96-well flat-bottom plate with 250 μL of complete medium. After 72 h of coculture, supernatant was collected as performed in the TCR-pCD8 experiment. Afterwards, hIL-2 in the supernatant were quantified by Infinite M Plex instrument using Human IL-2 ELISA Kit (D2050, R&D) according to the manufacturer’s instruction.

### Cell killing assay

Firefly luciferase–transduced target cells (1 x 10^4^ per well) were incubated overnight in 96-well flat clear bottom black plates (3904, Corning) with 100 μL of complete medium. The next day, the medium was replaced with fresh human T cell medium. In case of peptide-pulsed T2 cells, 5 x 10^4^ firefly luciferase–transduced cells were seeded into 96-well flat clear bottom black plates with human T cell medium on the day of coculture. Subsequently, 1 x 10^4^ TCR-pCD8 cells/well were added to the plates with human T cell medium. The final number of required pCD8 cells were determined by mouse-TCRcβ positivity ([1 x 10^4^) x 100/[mouse-TCRcβ positivity (%)] of total pCD8 cells). The number of empty-vector– transduced pCD8 cells, used as a negative control, was matched to the total number of pCD8 cells, leading to a final volume of 100 μL/well. After 24 h of coculture, cell viability was quantified on a FLUOstar Omega Microplate Reader (BMG LABTECH) using the ONE-Glo EX luciferase assay system (E8110, Promega) according to the manufacturer’s instructions. In case of T2 cells, the Bright-Glo Luciferase Assay System (E2610, Promega) was used. Luminescence data were obtained two times serially from the same plates to minimize time-dependent signal variability, and mean values from the 2 serial records were used for precise cell killing calculation. Percentage of cell killing was calculated using the following formula: % target-cell death = 100 × [(viability of coculture well − viability of target-cell only well)/(viability of 1% Triton-X-100–treated well − viability of target-cell only well)].

### HLA-A*02 competition assay

TAP1/2-deficient T2 cells were pulsed for 2 h with combinations of peptides as indicated (10 μM) at 1 x 10^6^/mL in complete medium followed by washing with complete medium (450g, 6 min, 4°C). Thereafter, the double-peptide–pulsed T2 cells were cocultured with TCR-J76^CD8αβ^ for 72 h. Conditioned media were then collected for IL-2 determination by ELISA, using the Human IL-2 ELISA Kit on an Infinite M Plex instrument. If the TCR-reactive peptides were less competent for HLA-A*02 binding than non-reactive peptides, IL-2 secretion from the TCR-J76^CD8αβ^ was abrogated because of failure to activate the T cells.

### MHC ligand prediction

NetMHCpan-4.1 (https://services.healthtech.dtu.dk/services/NetMHCpan-4.1/)^28^ was mostly used to list putative pHLAs for all possible 8–11-mers against indicated HLA-ABC alleles, and peptides were ranked based on %Rank_EL (EL rank), %Rank_BA (BA rank), and Aff(nM) (IC_50 v_alue of binding affinity). For the EL and BA ranks, 2% was used as a cut-off unless otherwise specified. In the initiation of our efforts, NetMHCpan4.0 and HLAthena^80^ were also used.

### Syngeneic mouse study

Seven-week-old female BALB/c mice (Cat.# 028, Charles River Laboratories) were inoculated subcutaneously on their right flanks with 1 x 10^3-5^ CT26^sg-control^ or CT26^Erap1-sg3^ cells resuspended in 100 μl of cold PBS. The number of mice was normalised between each condition. Tumour volume and body weight were monitored every 3–4 days until day 60. Tumour volume was defined as 1/2 × length × width^2^. Mice were euthanized when tumours became necrotic or grew to a length of 2 cm or a volume of 2000 mm^3^. All experiments were conducted according to a Dana-Farber Cancer Institute–approved protocol (22-013).

### Clinical database of HLA haplotypes in patients with cancer

HLA haplotype and p53 mutation status were obtained from The Cancer Genome Atlas (TCGA) program cohort and DFCI cohort.

For the TCGA cohort, p53 mutation status was obtained from TCGA PanCancer Atlas cohort (generated by the TCGA Research Network [https://www.cancer.gov/tcga]) for 20 tumour types which were also used in The Cancer Immunome Atlas (TCIA) project^81^ via cBioportal (https://www.cbioportal.org/)^82–84^. Data were available from 8,966 tumours in 8,955 patients. After integrating 11 replicated tumour samples biopsied from the same patients and excluding 513 patients for whom p53 status was not available (such as “NS”), 8,442 patients with a defined p53 mutation status remained. This core data was then filtered to yield a cohort of 5,407 patients with tumours harbouring p53^wt^, p53^R175H^, p53^H193Y^, p53^I195F^, p53^I195T^, p53^R249S^, or p53^R249M^. Full HLA-ABC haplotypes were available for 8,507 patients from the same cohort through TCIA project database (https://tcia.at/home)^81^ under NCBI dbGaP Data Access Request Approval (#125398-2 for the project 35503), where HLA haplotypes were not provided for 270 patients with tumours for the p53^wt^ and p53^mut^ listed above. This resulted in a final cohort with p53 status and full HLA haplotype for 5,137 patients.

For the DFCI cohort, patient-derived OncoPanel data were obtained under a clinical research protocol #22-556, which was approved by the Dana-Farber/Harvard Cancer Center Institutional Review Board. This protocol allowed secondary anonymous use of a preexisting clinical database built under protocol #16-360^85^. The #22-556 was designed to collect raw OncoPanel data anonymously from tumours positive for p53^R175H^, p53^H193Y^, p53^I195F^, p53^I195T^, p53^R249S^, or p53^R249M^, specifically for computational imputation of HLA-ABC haplotypes. Patient consent was thus waived given the minimal privacy risk. Consequently, 1,036 patients with tumours positive for the six selected p53 variants were included from the DFCI Profile Release Latest cohort^86^. Full HLA-ABC haplotypes for fields 1&2 (formerly defined as four-digits) were imputed with off-target reads from OncoPanel data using the computational pipeline as previously published^87^. After excluding 384 patients for whom sufficient OncoPanel data were not available, paired data of p53 status and full HLA haplotype were available for 652 patients. Procedures were performed in accordance with the Declaration of Helsinki.

For the final analysis for association between certain types of p53 mutations and HLA-A*02:01 or HLA-B*07:02, tumours with indel or frameshift p53 mutations causing ≥2 amino acids alteration within the corresponding peptide arrays were not counted as those positive for p53 mutations of interest.

### Single molecule (SM) and single cell activation requirement (SCAR) assays

Protein production of the two sets of TCRαβ heterodimers and pMHC ligands for SM assays used the corresponding sequences of the molecules described herein. Both SM and SCAR assays followed methods detailed previously^19,20^. For SCAR, R175H-TCR3-J76^CD8αβ^ and I195F-TCR-J76^CD8αβ^ cells were employed.

*(SCAR):* The SCAR assay was performed to monitor calcium flux within a cell upon presentation of pMHC through an optically trapped bead. This assay monitors early T cell activation and pMHC threshold requirements with and without force. Briefly, Quest-Rhod-4, AM (AAT Bioquest Inc.) was used to fluorescently label intracellular Ca^2+^ in the TCR-J76 cells. 1.18 µm polystyrene beads (Spherotech Inc.) were decorated with either ^187G^LAPPQHL<u>F</u>RV^197^- or ^168^HMTEVVR<u>H</u>C^176^- bound biotinylated MHC at varying interfacial copy numbers. Beads were manipulated via an optical trap and placed on a coverslip bound T cell. Cells were displaced relative to the fixed trap to apply force across the TCR-pMHC bonds. After pMHC bead attachment and force application, the T cell is monitored for 10 mins with fluorescent images captured every 5 s to track the calcium flux of the cell. Detailed protocols for the SCAR assay can be found in previously published works^19,20^. Adaptations made to the previous protocol include spinning down cells at 400 g for 3 mins and using RPMI colorless media (Gibco).

*(SM):* Single molecule measurements were collected via the optical trap surface assay as detailed previously^88^ and diagrammed in Extended Data Fig. 9a. The assay was constructed with a biotinylated-PEG coverslip functionalized with streptavidin and biotinylated HLA. The remaining non-functionalized surface was blocked with casein in PBS to minimize non-specific interactions (not shown in cartoon). An anti-leucine zipper antibody (2H11) conjugated to one 5’ end and biotin on the opposite 5’ end of a 1010-bp dsDNA was incubated for 20 mins with 1.18 µm streptavidin-coated beads (Spherotech). The given TCRαβ is then incubated with DNA coated beads for one hour in the presence of 0.5% w/v casein. Bond lifetime measurements were performed on both I195F-TCRαβ-pMHC (I195F/HLA-A*02:01) and R175H-TCR3αβ-pMHC (R175H/HLA-A*02:01) by holding a bead in the optical trap and scanning the surface with piezo-stage displacements to achieve surface bead tethers under load. Individual force loaded events were analyzed for their bond lifetime and binned every 5 pN and plotted as mean bond lifetime per bin ± SEM. Forces were corrected for the assay geometry by the factor 1/sin(Ø) which includes a ∼50-degree angle of the tether relative to a vector normal to the coverslip. Reversible transition distances in Extended Data Fig. 9d were determined via a position distribution representing the populations of points in each respective dwell in the compact or extended state of transition. This distribution was then fit to the sum of two Gaussian distributions as described in Das et al^88^. The difference in the peaks of the fits reveals the distance of the conformational transition.

## Statistical analysis

Statistical tests applied to compare variables and provide *P* values are described in main text or figure legends. Data from independent experiments are presented individually or combined, with indicated number of technical and biological replicates, as outlined in each figure legend. All *P* values are two-sided. *P* values less than 0.05 were considered statistically significant. Asterisks used to indicate significance correspond to: *p<0.05, **p<0.01, ***p<0.001, ****p<0.0001. Statistical analyses and data visualization were performed using GraphPad Prism 10.2.2 (GraphPad Software), unless otherwise specified.

## Data availability

Atomic coordinates and structure factors for the reported crystal structures have been deposited in the Protein Data Bank under accession numbers 9OBF (^187^GLAPPQHLIRV^197^/A*02:01), 9OBG (^187^GLAPPQHLFRV^197^/A*02:01), 9EIQ (^249^RPILTIITL^257^/B*07:02), 9EK4 (^249^MPILTIITL^257^/B*07:02), and 9EJD (^249^SPILTIITL^257^/B*07:02).

All data generated or analyzed during this study are included in this article and its supplementary information file. Source data are provided with this paper.

## Code availability

Implementation in individual MS user-unique environments is possible using data provided in the manuscript. Specific queries should be sent to B.R.

## Acknowledgements

This research was supported by NCI RO1CA265928 (E.L.R.), NIAID PO1AI143565 (E.L.R. and M.J.L), Novartis Global Scholars Program (D.A.B. and E.L.R.), NIH Lung SPORE P50CA265826 (D.A.B.), NIH R01CA190294 (D.A.B.), the Ludwig Center at Harvard Medical School (D.A.B.), the Mark Foundation (D.A.B), the Heerwagen, Candice Bagby, and Ming and Polly Tsai Funds for Lung Cancer Research (D.A.B.), the Takeda Science Foundation (K.H.), the Osaka Medical Research Foundation for Intractable Diseases (K.H.), the Mochida Memorial Foundation for Medical and Pharmaceutical Research (K.H.), the Nakatomi Foundation (K.H.), and NIH training grant T32 DK101003 (E.L.H.). Use of the NYX beamline 19ID at NSLS-II was supported by NYSBC. NYX detector instrumentation was supported by NIH grant S10OD030394. This research used resources of the National Synchrotron Light Source II, a U.S. Department of Energy (DOE) Office of Science User Facility operated for the DOE Office of Science by Brookhaven National Laboratory under Contract No. DE-SC0012704. We acknowledge Evan Kirkpatrick and Hannah Stephens for assay contributions and helpful advice, and Jonathan Lee and Kaveri Uberoy for expressing and purifying proteins for SM assays. We also thank Yanan Kuang, Elliott Brea, Tetsuo Tani, Cedric Louvet, Shunsuke Kitajima, Atsuko Ogino, and William Feng for their technical support. We appreciate Nicholas R.J. Gascoigne for providing cDNA information about EBV-LMP2A-TCR. We thank Eric Smith for graphic design of figures.

## Author contributions

Conceptualization, K.H., B.R., D.A.B., and E.L.R.; Methodology, K.H., B.R., J.S.D.-C, R.J.M., A.G., K.L.K., J.L., C.J.H., R.B.B, P.L., C.P.P., A.J.A., K.L.L., R.C., M.J.L., D.A.B., and E.L.R.; Investigation, K.H., B.R., J.S.D.-C, C.G.F., K.T., R.J.M., A.G., R.B.B, T.C.T., G.M.G., S.K., E.L.H., D.J.M., A.K., K.J.Z., and E.L.R.; Writing-Original Draft, K.H., B.R., J.S.D.-C, D.A.B., and E.L.R.; Writing-Review and Editing, all authors; Funding Acquisition, K.H., D.A.B., and E.L.R.; Supervision, D.A.B., and E.L.R.

## Additional information

Supplementary Information is available for this paper.

Correspondence and requests for materials should be addressed to

---

**Extended Data Fig. 1.**
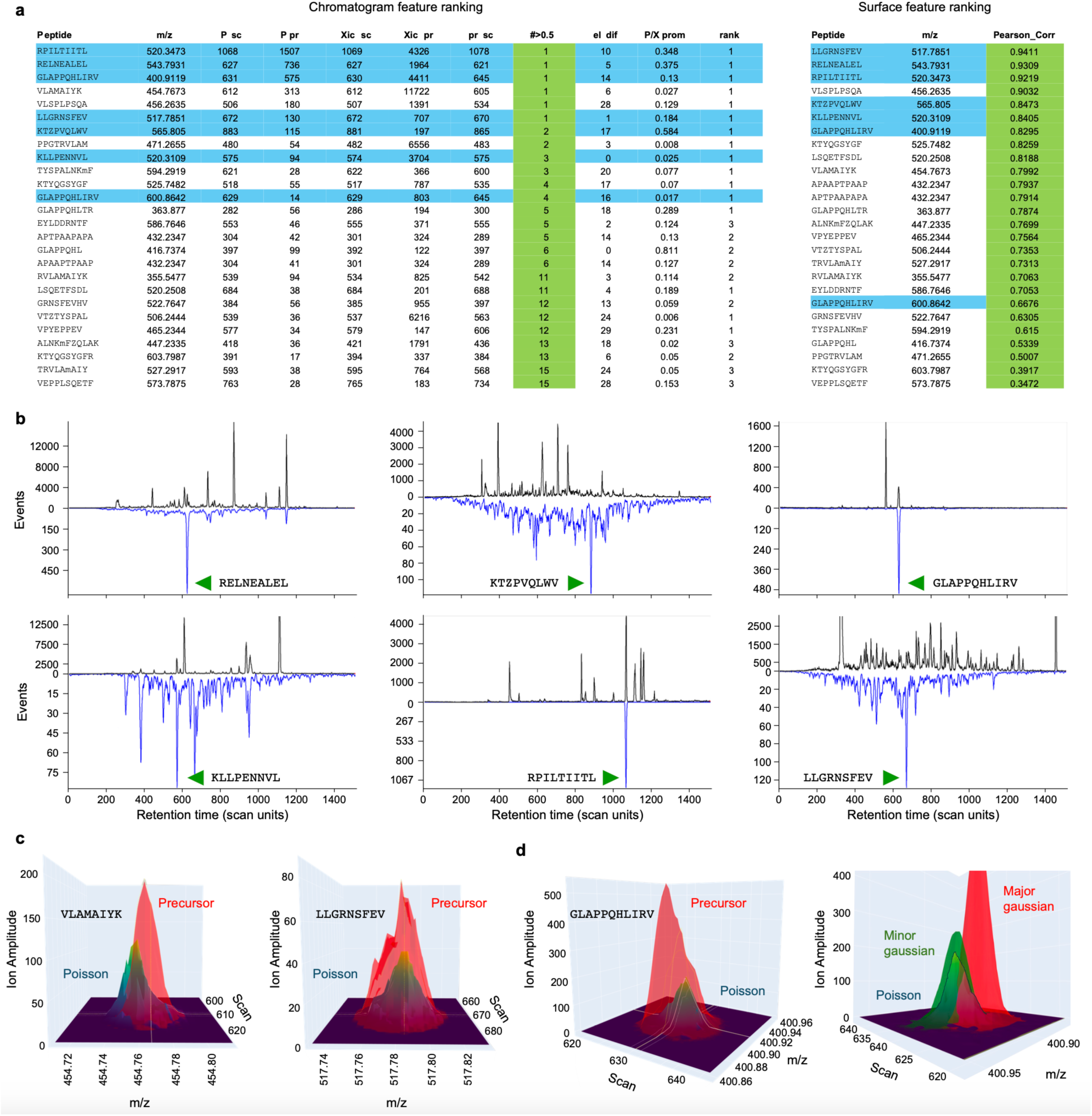
Poisson LC-DIA-MS detection of p53 peptides expressed by HLA in proliferating T cells. **a,** *Left panel*. Top ranking p53 peptides detected in a sample of 1 million stimulated T cells (see Methods). Features are extracted from precursor XICs and Poisson chromatograms (examples, Fig. 1b) corresponding to 251 reference patterns from 212 distinct p53 peptides (39 of 251 peptides are targeted in both +2 and +3 charge state). The rows are ranked by column features and cutoffs for ‘out of bound’ features are applied, with a final ranking along the green significance column ‘#>0.5’. This is the number of other peaks in the Poisson chromatogram with a peak prominence (baseline corrected amplitude) greater than half the prominence of the Poisson peak due to elution of the peptide. ‘rank’ describes the target prominence relative to the height of the other prominences in the Poisson chromatogram. The ‘el dif’ column shows the distance off the elution line for the target’s elution. The elution times of the 251 synthetic peptides (Syn) with a set of added retention time (RT) peptides is measured (in scan units). The RT peptides are also added to the sample. A polynomial map fits Syn RT to sample RT scans (elution line), and the same mapping predicts the elution times of peptides from the Syn set in the sample run. See Supplementary Discussion 1 for further details. *Right panel*. For each of the peptides displayed in the left panel, ion map surfaces in a local [scan, m/z] domain around the elution of the precursor and expected fragments were extracted. Poisson surfaces were calculated from the set of fragment surfaces (Supplementary Discussion 1) and the Poisson surface compared with the precursor ion surface by Pearson correlation. The peptides in this table were then ranked (0-1, green column). The HLA alleles for the T cell sample were identified as A*02:01(homozygous), B*07:02, B*40:02, C*02:07 and C*07:02. VLSPLPSQA is calculated to be a strong binder to HLA-A*02:01 (%EL rank 0.183) but is not detected in the H2228 system. RELNEALEL was predicted a weak HLA-B*38:01 binder for H2228 cells (hence synthesized) but was not observed there. However, it is predicted to be a strong binder for T cell sample’s HLA-B*40:02 (%EL rank 0.019) and is present. **b,** Paired precursor XIC (black trace, top) and Poisson chromatogram (inverted blue trace) for the 6 detected peptides. The features listed in ED Fig. 1a, left panel, are extracted from these traces (and RT elution positions). **c**, Precursor and Poisson surfaces for VLAMAIYK and LLGRNSFEV visually illustrating the ranking by Pearson correlations (poorer to better) as listed in ED Fig. 1a, right panel. **d**. GLAPPQHLIRV’s lowering of rank position by Pearson correlation relative to chromatographic rank position reflects a precursor ion surface that is a composite peak. The data processing chain fits a pair of Gaussian surfaces to precursor ion surfaces for all the peptides in the chromatogram list (ED Fig. 1a). Pearson correlation between the major and minor components and other parameters characterizing the fit are compiled. High Pearson correlation between the minor fit component and the Poisson surface are confirmed by 3D visual inspection (see Supplementary Discussion 1).

**Extended Data Fig. 2.**
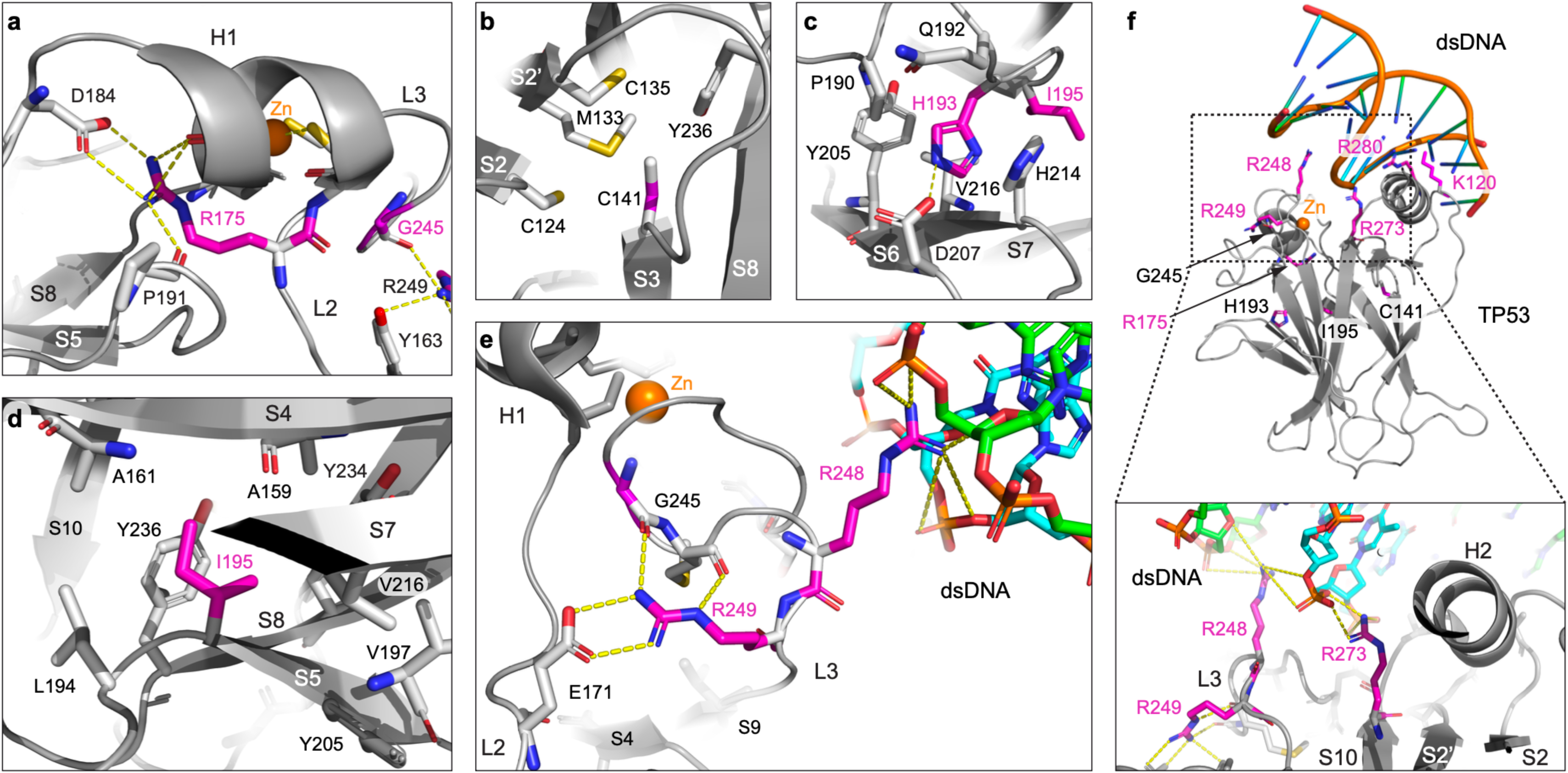
Location of hotspot missense mutations on the p53 DNA binding domain defining the structural basis for their capacity to mediate functional disruptions. **a,** A zoomed-in view of the residue R175 on the L2 loop (L2) of the DNA binding domain (DBD) of p53 that is drawn in grey ribbon. Through a bidentate salt-bridge to D184 and hydrogen bonds to other neighboring residues, R175 provides critical structural support for the L2 loop, H1 helix (H1), the Zn coordination (involving residues C176, H179, C238, and C242) and other structural elements shown. R175 and its interacting residues as well as other relevant residues are drawn in stick format. Salt bridge and hydrogen bonds are displayed in dashed lines. The Zn ion is shown as an orange sphere. Two β-strands, S5 and S8, are labelled. The details of the interactions involved G245 on L3 loop are best viewed in Fig. 1d. **b,** The residue C141 is at the beginning of the β-strand S3. This cysteine is important for the proper packing of the hydrophobic core of the DBD, especially with its neighboring residues that include two other cysteines, a methionine and a tyrosine. **c,** Residue H193 on the loop between H1 and the β-strand S5 (not shown) is sandwiched by P190 and H214. It also forms a displaced T-type stacking with Y205 and a hydrogen bond with D207. It helps stabilise one edge of the DBD β-sandwich structure, especially the two edge β-strands, S6 and S7. **d,** The residue I195 located at the beginning of the central S5 β-strand contributes to the tightly packed hydrophobic core of DBD that would be disrupted by a polar residue or a larger hydrophobic residue. **e,** The L3 loop harbours two consecutive arginine residues, R248 and R249. R248 is one of key residues in the interaction of p53 with DNA while R249 provides structural support. In this regard, R249 forms two hydrogen bonds to the two carbonyl groups on the same loop and a bidentate salt-bridge to E171 on the L2 loop. It also forms a cation-π interaction with Y163 at the end of S4 β-strand (not shown), apparently stabilising the conformation of the important long L3 loop. **f,** The interaction between p53 and dsDNA. The entire DBD is displayed in grey ribbon with those hot spot residues described in **a**–**e** as well as four DNA binding residues K120, R248, R273 and R280 being mapped on the domain. All hot spot residues and DNA-binding residues are drawn in magenta sticks for highlight. The bottom insert is a zoomed-in view of the interactions contributed by R248 and R273. R248 points into the minor DNA groove, forming multiple salt links to backbone phosphates (orange) of both DNA chains. R273 interacts with the backbone phosphate of a single DNA chain. All figures were prepared based on the p53 structure (PDB code:2OCJ) and p53/DNA complex structure (PDB code: 1TUP).

**Extended Data Fig. 3.**
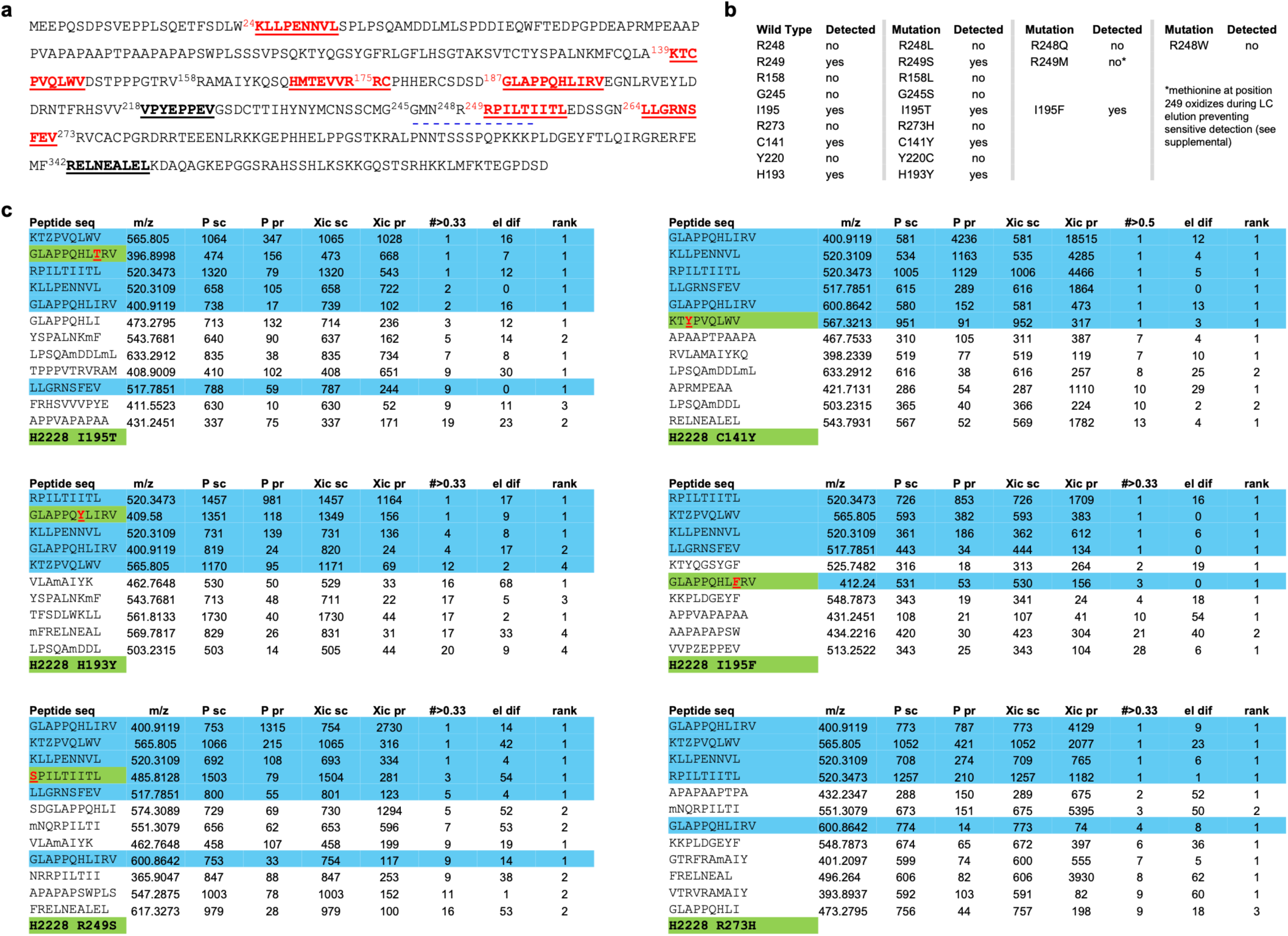
Positive detection of additional p53 neoepitopes and concordance of wt and mutant-related peptide presentation. **a**, The sequence of wild type p53 with epitopes detected in the H2228 system marked in **<u>bold</u> <u>red underline</u>**. The unmutated peptide segment corresponding to the R175H neoantigen is also designated. <u>Bold black underline</u> marks epitopes detected in other cells with different HLA restrictions (RELNEALEL B*40:02, VPYEPPEV B*51:01). <u>The blue dashed underline</u> shows position of GMNRRPILTI 10-mer and its conflict with the generation of the RPILTIITL epitope. **b**, Sequences containing the indicated amino acid/position in wild type p53 and whether any epitopes incorporating that position were detected in the H2228 system are listed (left two columns). In the middle two columns H2228 cell were transduced with mutant *TP53* and detection status of the corresponding epitopes indicated. The last 2 columns correspond to transduction of multiple mutants at the 195, 248, and 249 amino acid position. **c**, Top-ranked Poisson chromatogram features for mutant *TP53* transfections. The mutation and the mutated epitope when detected are highlighted in green. Mutations of sites contained in wild type epitopes were detected except for R249M, a problem associated with the methionine oxidizing during the chromatography (see Supplementary Discussion 2). The R273H mutation modifies a site adjacent to the carboxy terminus of the ^264^LLGRNSFEV^272^ epitope and is correlated with loss of detection.

**Extended Data Fig. 4.**
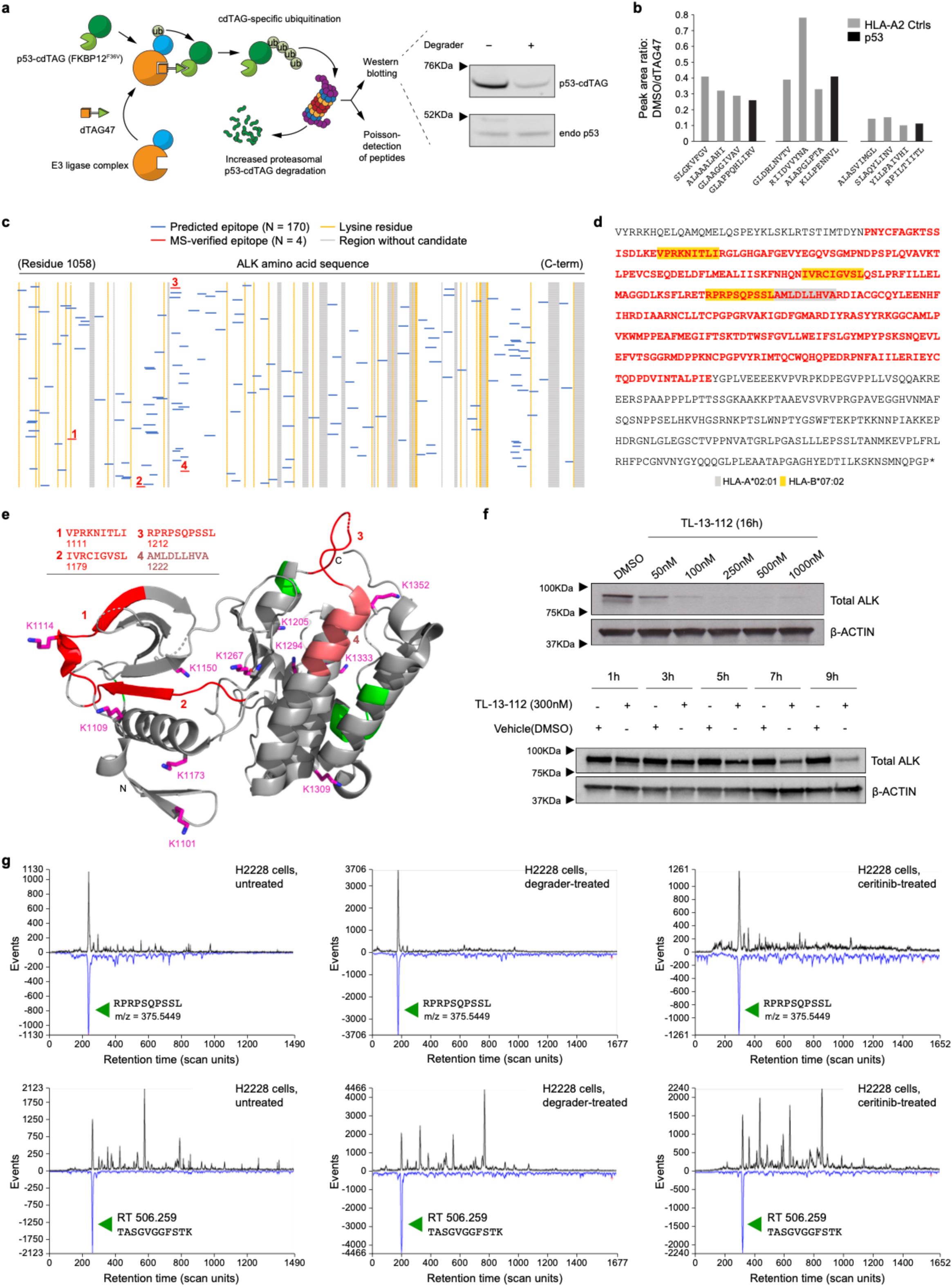
Facile degradation of both p53 and anaplastic lymphoma kinase (ALK) without detectable quantitative enhancement or qualitative shift in their tumour antigen peptidome displays by MS. **a**, Schematic of p53 detection and immunopeptidome profiling after specifically targeted proteasomal degradation using the dTAG system with accompanying Western blot. **b,** The ratio of peak areas for the p53 peptides between DMSO control and TAG47 treated cells for three p53 peptides and nine HLA-A*02:01 sample peptides unrelated to p53 (‘HLA-A2 Ctrls’). To normalise the differential expression of p53 epitopes between the control and TAG47 treated cells, sets of three endogenous HLA-A2 peptides (each set eluting around a different p53 peptide) were monitored. In the TAG47 samples the intensities of p53 and A2 controls were reduced compared with the DMSO treated cells, especially late in the gradient where RPILTIITL elutes. The p53 peptides largely track the same as the A2 controls. **c**, position of candidate epitopes within ALK region from the EML4-ALK fusion protein including the entire kinase domain. Color coding and schema as in Fig. 1c. **d**, An amino acid sequence of ALK region from the EML4-ALK fusion protein with kinase domain in red font and position of HLA-A*02:01 (grey) and HLA-B*07:02 (yellow) epitopes. **e**, structure of kinase domain and position of four epitopes found collectively in lymphoid and epithelial tumours^22^ with only epitope 3 found in H2228. **f**, Immunoblot analysis demonstrating ALK degradation in the ALK-rearranged H2228 cell line. Cells were seeded at a density of 1 x 10^6^ per 10 cm plate and after 12 hours treated with the ALK-specific degrader TL-13-112. The top panel cells were treated for 16 hours at indicated concentrations and the bottom panel harvested at the specified time points flowing treatment with degrader (at 300 nM concentration) or vehicle control. **g**, MS detection reveals no significant biological increase 9 hours after TL-13-112 (the kinase inhibitor ceritinib conjugated to cereblon E3 ligase ligand) treatment of the HLA-B*07:02 epitope 3 (top row) relative to the retention time peptide used as elution position marker. Untreated or treatment with ceritinib kinase inhibitor alone results are shown as additional controls. Poisson detection chromatograms for H2228 cells are shown for untreated, treated with ALK degrader TL-13-112 and ceritinib. For all the panels the bottom blue trace identifies the elution of a target’s fragmentation pattern, and the top black trace is the extracted ion chromatogram (XIC) for the m/z of the target. The amplitude of the XIC peak corresponds to peptide abundance. The top 3 panels follow the ALK-sourced peptide RPRPSQPSSL while the bottom three track a retention time peptide in the same sample as in the above panel. A fixed amount of retention time peptides are added to all sample runs with the TASGVGGFSTK peptide eluting near RPRPSQPSSL. The XIC peaks normalise the ion signals between the different runs: 1130/1250 = 0.9 (untreated), 3706/2000 = 1.9 (degrader), 1261/1500 = 0.8 (ceritinib) with less than two-fold change considered small.

**Extended Data Fig. 5.**
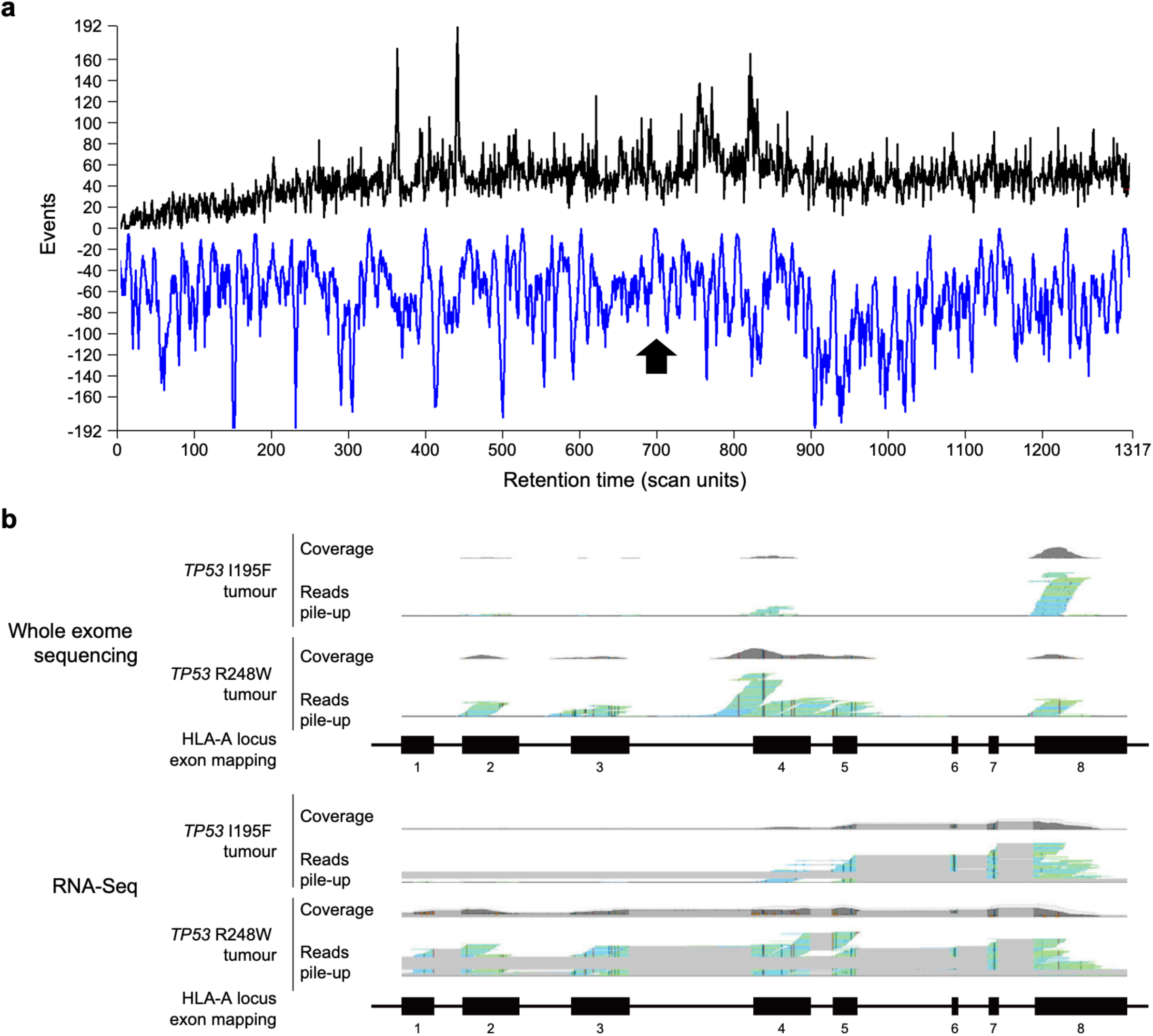
An aberrant HLA-A locus leads to loss of allele expression and that of the p53 driver neoepitope I195F in the immunopeptidome. **a**, Poisson chromatogram showing absence of the epitope (arrow). **b**, Both whole exome sequencing (upper lanes) and RNA-Seq (lower lanes) revealed minimal reads covering HLA-A locus exons 1–3 from a brain metastasis biopsy of a patient with p53^I195F^ HLA-A*02:01/A*29:02 ovarian cancer on comparison with a p53^R248W^ HLA-A*24:02/A*32:01 glioblastoma biopsy sample prepared and sequenced simultaneously. For each sample, reads are presented as exon coverage (upper grey distributions) and as a pile-up where blue lines represent 5’-3’ (forward) reads and green lines represent 3’-5’ (reverse) reads aligned to the HLA-A locus exon mapping depicted below. The grey connecting regions in the RNA-Seq read pile-up indicate transcript reads that span splice junctions.

**Extended Data Fig. 6.**
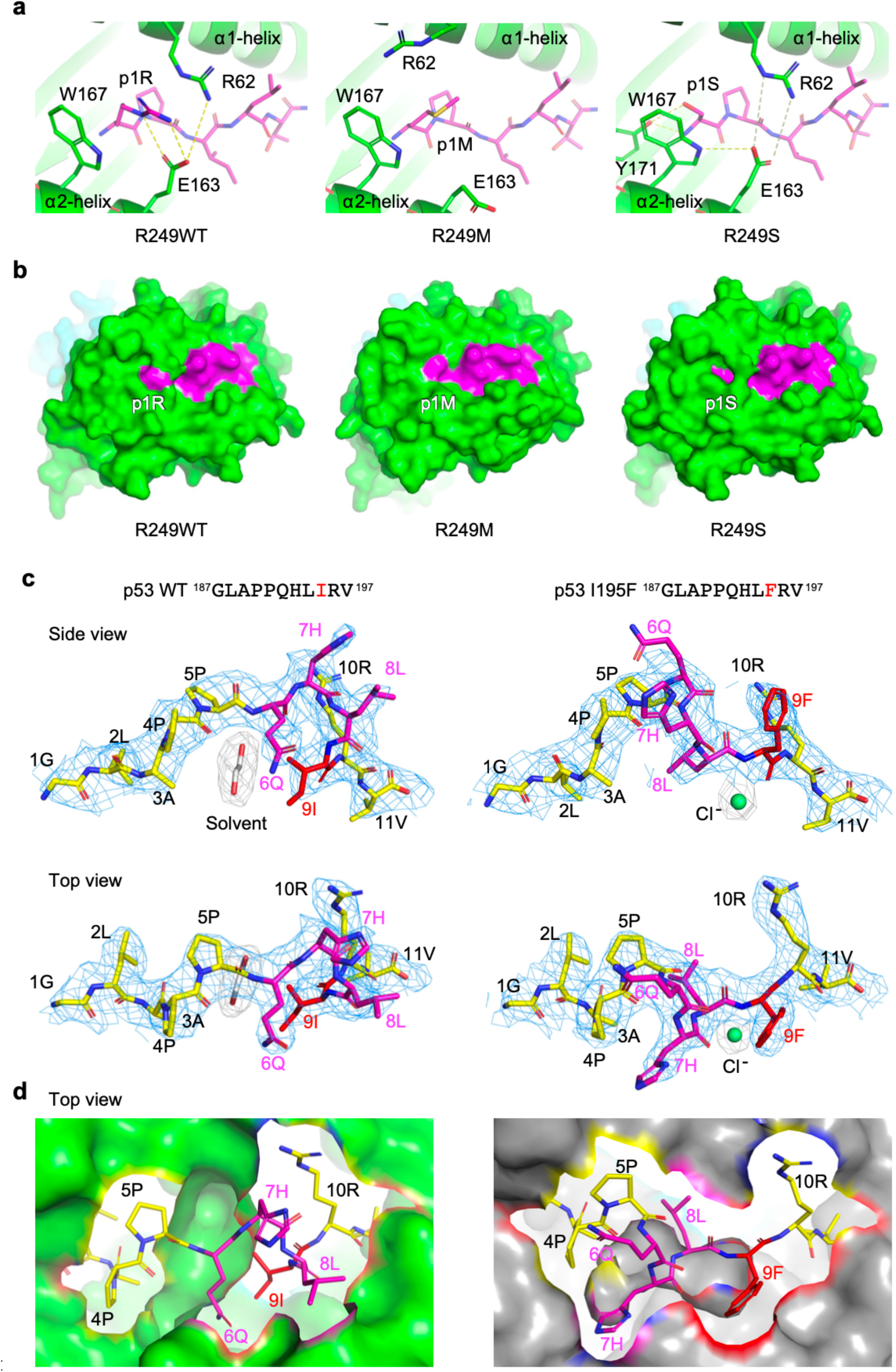
Structural analysis of immunodominant neoantigens and their normal p53 immunopeptidome counterparts. **a**, Ribbon diagrams illustrate the interactions between HLA-B*07:02 and the p53 WT ^249R^PILTIITL^257^ peptide, as well as the R249M and R249S mutants (from left to right), focusing on the N-terminal region of the bound peptide. The peptides are drawn in sticks with carbon atoms colored magenta. HLA-B*07:02 is colored green, with the key interacting residues, drawn in green sticks. Salt bridges and hydrogen bonds are indicated by dashed lines. **b,** Space filled rendering of MHC in green and indicated peptides in pink, highlighting the recessed p1S residue of R249S and the bidentate salt link (a) between the MHC R62 and E163 residues of the alpha 1 and alpha 2 helices, respectively, creating the side wall of the cavity/crevice. **c**,**d**, Structural analysis of p53^wt^ and p53^I195F^ 11-mer peptides bound to HLA-A*02:01. In c, the wt and mutant peptides are displayed in stick format with unaltered segments colored yellow. For wt peptide, the light cyan mesh represents the σA-weighted 2Fo–Fc electron density map associated with the peptide using a contour level of 1. A density between the bulged part of the peptide and the peptide binding cleft of HLA-A*02:01 below (not shown) is presumably contributed by a solvent or solvents from buffers. An ethylene glycol molecule was built into the density for structural refinement purpose. For p53^I195F^, an identical contour level is shown. A globular density attached to the amide group of 9F is interpreted as a Cl^−^ anion, which also forms interactions to the side chains of R97 and H114 of HLA-A*02:01 from its peptide binding cleft (not shown). Both side and top views are shown. The view in panel d is with MHC surface in green and grey for p53^wt^ and p53^I195F^ 11-mer complexes, respectively and looking from the TCR perspective.

**Extended Data Fig. 7.**
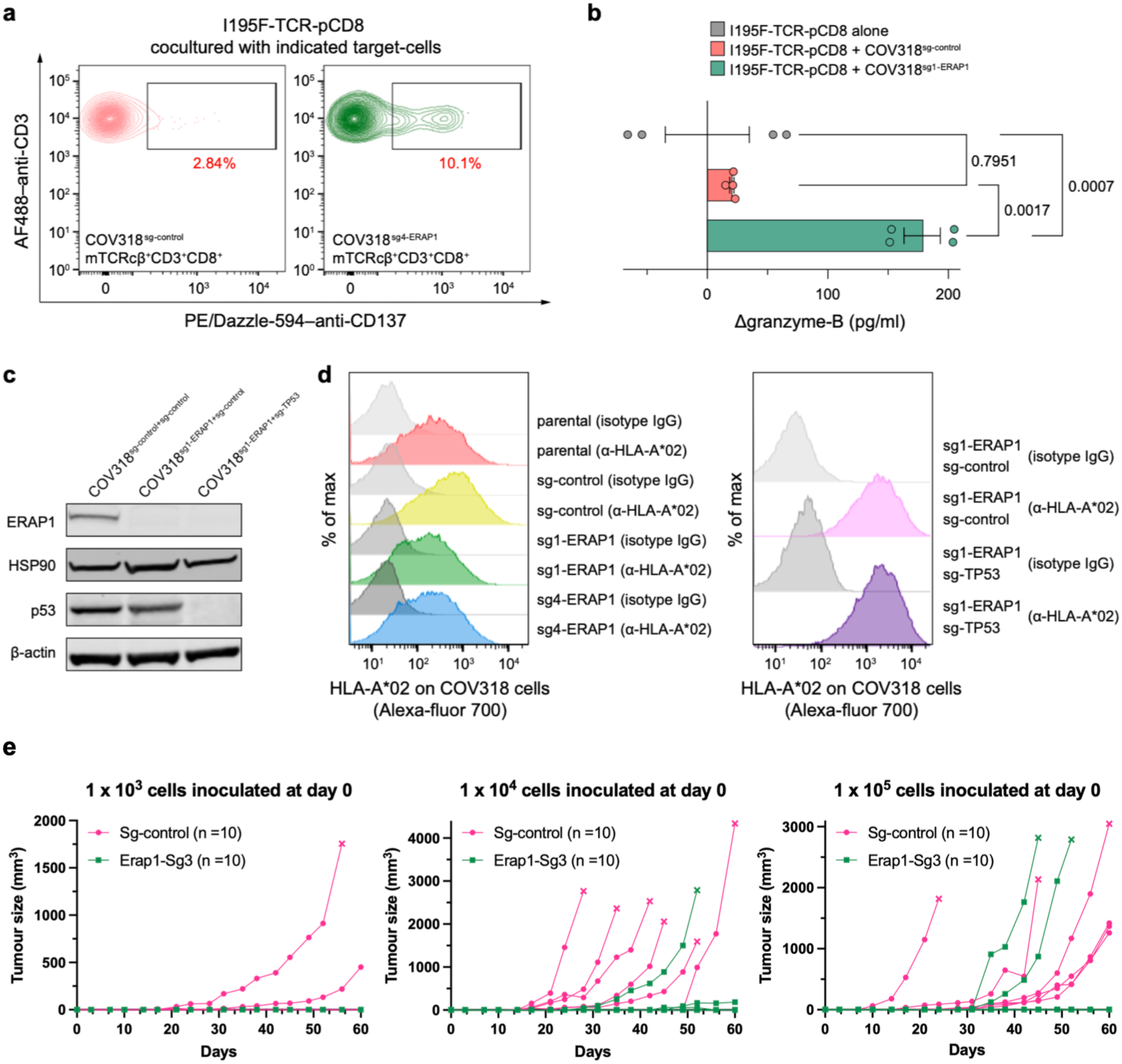
Abrogation of tumour ERAP-1 *in vitro* and *in vivo* restores induction of anti-cancer T cell immunity. **a-d**, *in vitro* studies. Flow cytometry data of CD137 expression (**a**) and ELISA data of granzyme B (**b**) as T cell activation markers. I195F-TCR-pCD8 cells were cocultured with target-cells as indicated for 24 hours beforehand. **a**, mTCRcβ^+C^D3^+^CD8^+^ live singlets were gated as indicated to obtain the CD137 expression data. Data are shown by contour plots with outliers as individual dots. Frequency of CD137^+^ cells are shown in red letters. **b**, The results are presented by Δgranzyme-B ([granzyme B value of each sample] – [the mean value of granzyme B of technical replicates in an I195F-TCR-pCD8 alone sample]). Mean values and standard error of the mean (SEM) (bars) and individual values from four technical replicates (circles) are shown. *P* values are calculated by one-way analysis of variance (ANOVA) with Tukey’s multiple-comparison test. **c**, Immunoblotting of p53 proteins in COV318 with *TP53*-KO. COV318 with sg-control (left) or sg1-ERAP1 (middle) are positive control for p53 protein. **d**, Flow cytometry data of HLA-A2 expression on COV318 cells with *ERAP1*-KO (left) or *TP53*-KO (right). COV318^parental^ cells, COV318^sg-control^ cells, and isotype control were used as negative control. Data are representative of two independent biological replicates throughout. **e**, Individual tumour growth curves in mice used in Fig. 5j–l are shown. “X” on the curves indicate that the tumours met euthanisation endpoints.

**Extended Data Fig. 8.**
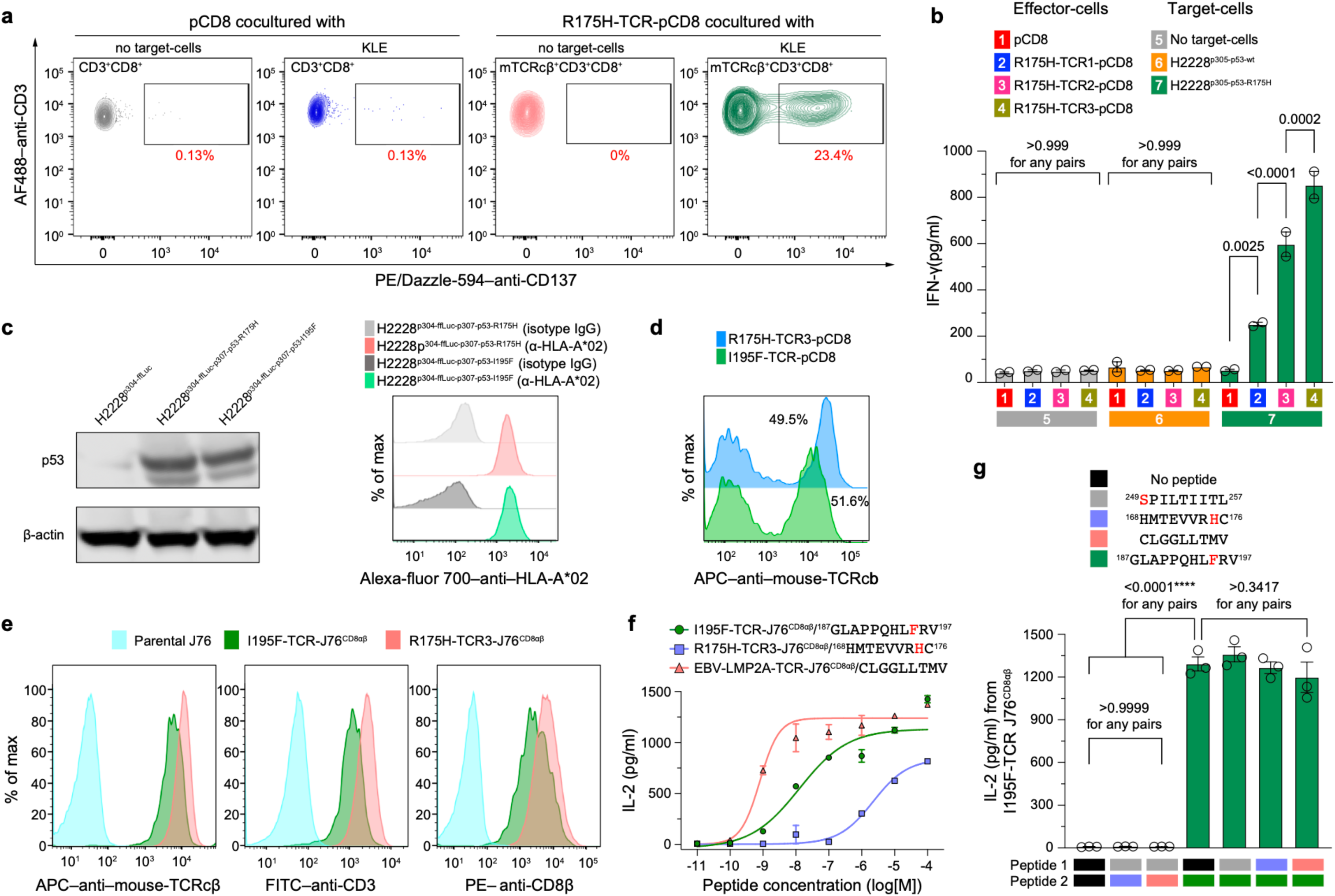
Data for comparison of p53 neoepitope quality. **a**,**b**, Flow cytometry data of CD137 expression (**a**) and ELISA data of IFNγ (**b**) as T cell activation markers. R175H-TCR-pCD8 cells were cocultured with target-cells as indicated for 24 h beforehand. **a**, CD3^+^CD8^+^ or mTCRcβ^+C^D3^+^CD8^+^ live singlets were gated as indicated to obtain the CD137 expression data. Data are shown by contour plots with outliers as individual dots. Frequency of CD137^+^ cells are shown in red numbers. **b**, Mean values and standard error of the mean (SEM) (bars) and individual values from two technical replicates (open circles) are shown. *P* values are calculated by one-way analysis of variance (ANOVA) with Tukey’s multiple-comparison test. **c**, Immunoblotting of p53 protein (left) and flow cytometry data of HLA-A2 expression (right) in H2228^p304ffluc^ cells transduced with p53^R175H^ or p53^I195F^. H2228^p304ffluc^ is negative control for full-length p53 protein (left). Isotype control was used for negative control (right). **d**,**e**, Flow cytometry data of transduced TCR cell surface expression, as labeled by mTCRcβ, on TCR-pCD8 cells (**d**) and TCR-J76^CD8αβ^ cells (**e**). Expression levels of CD3 and CD8 on cell surface are also demonstrated for J76 cells (**e**). **f**, Dose-response curves of IL-2 for the indicated pairs of TCR-J76^CD8αβ^ cells and peptides pulsed on T2 cells. T2 cells were pulsed with peptides at indicated concentrations for 2 h, and then cocultured with TCR-J76^CD8αβ^ cells for 72 h. Mean values and SEM (bars) and individual values from three technical replicates are shown. **g**, Results of HLA-A*02:01 competition assay for I195F-TCR-J76^CD8αβ^ are shown. Mean values and SEM (bars) and individual values from three technical replicates (open circles) are shown. Data are representative of two independent biological replicates throughout. *P* values are calculated by one-way ANOVA with Tukey’s multiple-comparison test.

**Extended Data Fig. 9.**
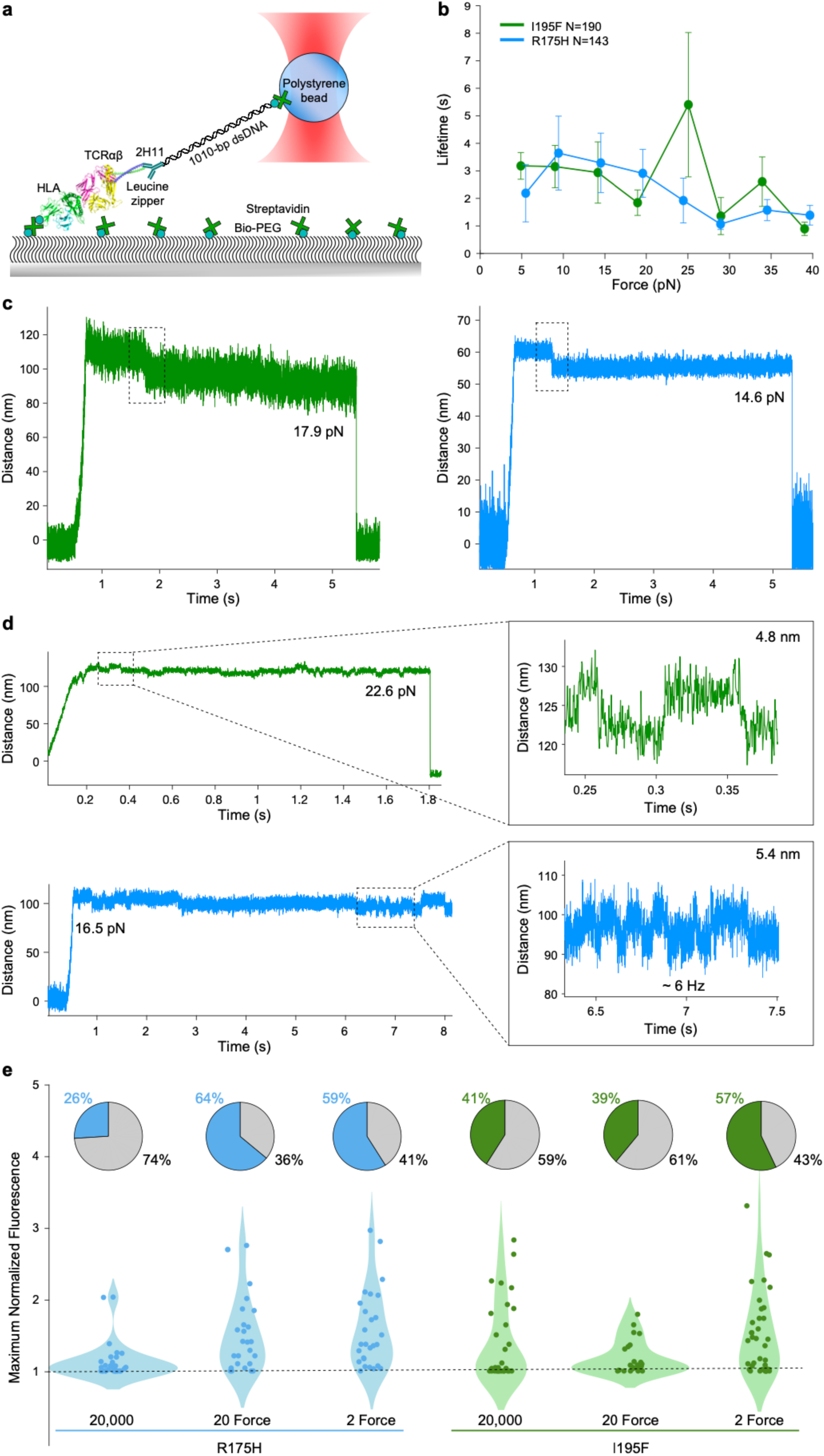
Both R175H-TCR3 and I195F-TCR exhibit digital performance characteristics. **a**, Single molecule (SM) system cartoon using a pHLA coated surface and TCRαβ heterodimer tethered to an optically trapped polystyrene bead. **b,** Bond lifetime vs. force plots for I195F (green, N = 190) and R175H (blue, N = 143) TCRs interacting with their respective ligands. The color convention is consistent through all of figure. Lifetime is plotted as mean bond lifetime ± SEM with bins every 5 pN. Note catch bond characteristics of both with higher maxima peak for I195F-TCR system. **c**, Representative traces for I195F-TCR (left) at 17.9 pN and R175H-TCR3 (right) at 14.6 pN displaying conformational transitions (dotted boxes) of 11.9 nm and 5.2 nm, respectively, representative of force dependent αβTCR heterodimer extension. **d**, Representative traces showing reversible conformational transitions for I195F-TCR and R175H-TCR3 at 22.6 pN and 16.5 pN respectively, a feature of mechanosensing. **e**, Single cell activation requirement (SCAR) assay measuring TCR-triggered calcium flux upon ligand binding as an indicator of early T cell activation. Maximum normalised fluorescence and triggering percentage of R175H- and I195F-TCR-T cells stimulated with ∼20,000, 20, or 2 pMHC molecules during SCAR assay using R175H-TCR3-J76^CD8αβ^ and I195H-TCR-J76^CD8αβ^ cells. Each dot on the violin plot represents the maximum normalised fluorescence of a single cell during the SCAR assay. A normalised fluorescence around 1 is indicative of a non-triggering cell (dotted line), while higher levels indicate that the cell has been activated. The pie charts above the violin plot show the triggering percentage of each bin (20,000 R175H N = 31, 20 Force R175H N = 25, 2 Force R175H N=27, 20,000 I195F N = 32, 20 Force I195F N = 23, 2 Force I195F N = 37) where grey is the untriggered cell fraction. From this data, both cell lines appear to behave in a digital manner, where a significant fraction of T cells can activate at limiting peptide copy number (i.e., 2 pMHC) with the aid of force ranging from 6–9pN.

**Extended Data Table 1.**
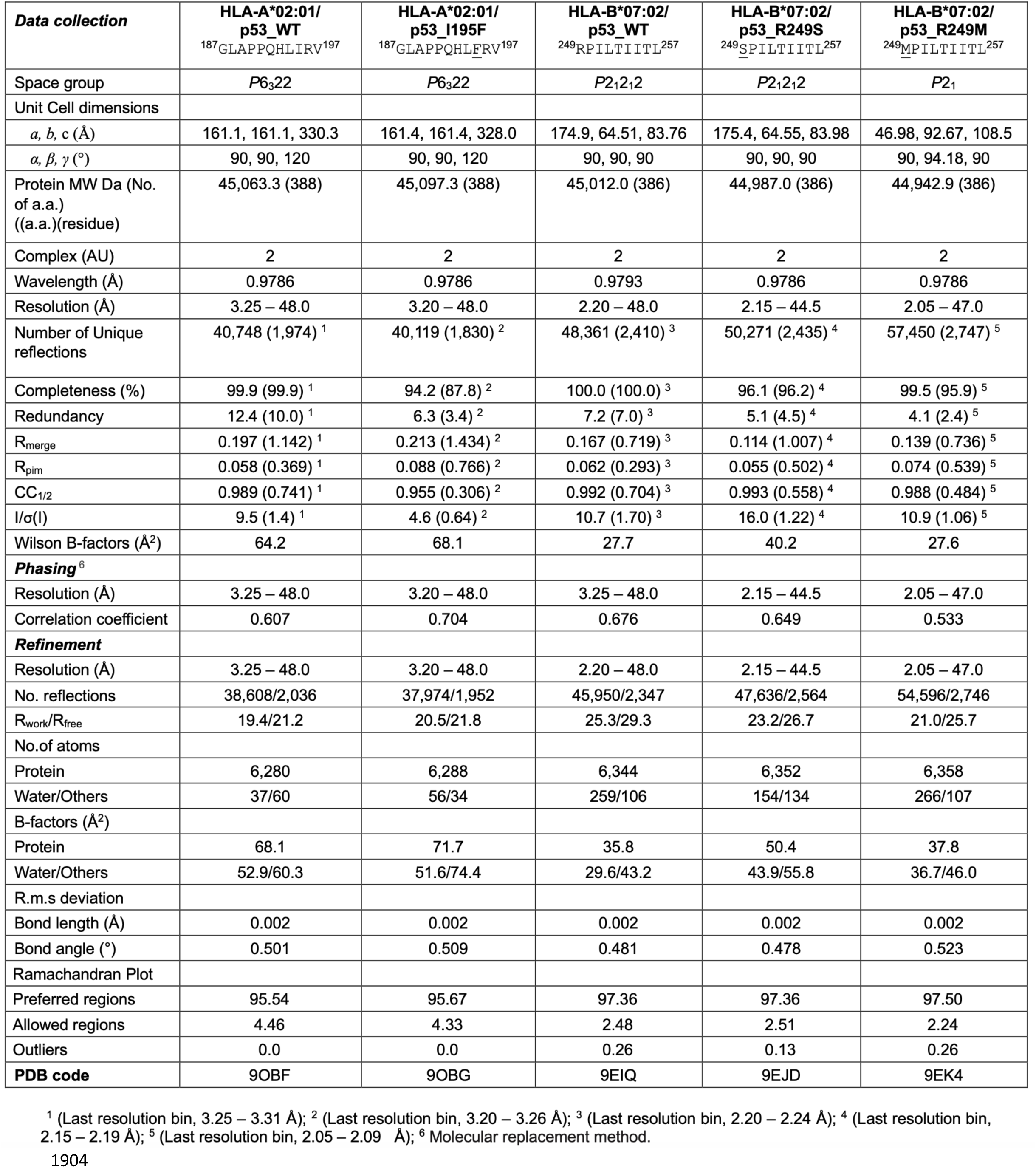
X-ray crystallographic data collection and refinement statistics.

