## Supplementary material for "Cancers modulate p53 truncal neoantigen display to evade T cell detection": Supplmentary Discussion 1

### Supplementary Discussion 1: <u>Detailed MS Methodologies, Concepts and Examples</u>

The following problem of interest in clinical oncology frames our approach to MS detection of tumour targets that characterize a cancer and may be actionable therapeutically. An immunohistology/pathology review has identified a tumour from needle biopsies, and both genomic and transcriptomic next generation sequence data have been acquired. Somatic mutations have been called, containing single nucleotide polymorphisms, deletions and frame errors. From those data, both known tumour associated antigens reflecting developmentally dysregulated or over-expressed genes as well as neoantigens have been identified. The data implicates hundreds of candidates for T-cell focused immunotherapy but very few of these candidates are expected to be presented as peptide-HLA complexes on the tumour cell surface. Taking the 'typical' mass of an epithelial cell as 2 ng, a biopsy mass of 10 mg wet tissue would contain 5 million cells. However, clinical biopsies often contain cellular stroma and extracellular matrix and fibrotic matter, so we set 1 million tumour cells as characterizing analytical scales. Taking 10 copies/cell as a relevant for CD8 T-cell recognition, one has 17 attomoles to detect and to do this in the setting of affinity purification, acid elution and peptide cleanup prior to LC-MS.

As a preamble to an outline of Poisson sampling, it is worth noting general directions with DIA MS. One path is to make DIA peptide spectrum matching (PSM) more like DDA PSM. Ion mobility front ends generate a time dimension to MS events, providing an additional isolation parameter for extracting a precursor's fragments in processing the data. timsTOF PASEF[1] further couples collisional cross section with DIA window selection by using a time-resolved extraction of ions initially immobilized in the z-direction of a stacked ring rf trap by opposed electric and collisional forces. In principle, PASEF timsTOF surveys ions during an extraction cycle and uses the (m/z, t) pairs to construct specific time-resolved DIA windows that are applied to subsequent ion extractions. For MHC class I peptides, a limited sequence length clusters both their m/z range and collision cross section into more compact domains than with proteomics. For these peptides a relation between ion mobility and mass for a charge state can be used as in a recent study[2] (Linear Scheduling, 5 DIA windows). In this setting how complex are the MS/MS windows? There are plots of inverse mobility versus m/z shown but for the set of all identifications in the run. The m/z distribution vs chromatographic time dimension is not shown. As the samples are monoallelic cells with 12 and 8 million cells ('two different concentrations'), here one has both a rather large sample and a comparatively simple immune peptidome. It's difficult to infer the signal levels in - and complexity of - the MS/MS windows behind the identifications and translate this into a clinical setting where samples are smaller, the allelic diversity more complex and the chemical background more pronounced. Another PASEF timsTOF study implements MS2Rescore to significantly improve identification depth[3]. Although MS2Rescore does not address feature extraction, it uses machine learning models to predict both fragmentation and elution to rescore PSMs. That rescoring with computer-predicted models gave significant improvement underscores a utility of starting with synthetic peptides where these features have been determined, even using latest generation instruments.

A second trend in DIA MS connects feature extraction and sequence identification not by a direct scoring of the feature set but by using the features for classification with machine learning algorithms that have been trained on supervised (known true, false) data[4,5]. For low level peptide signals embedded in DIA data the extraction of useful features associated with individual fragment traces is difficult. A current practice is to generate a high dimensional feature vector rather casually and let adaptive learning sift out what is useful for classification. A 2020 paper on DIA-NN[4] has '73 features' extracted for machine learning and subsequent inference. Example features: "*Pearson correlations of top 12 fragments' elution profiles with the smoothed elution profile of the "best" fragment (12 scores). Sum of these correlations for the top 6 fragments, calculated at different mass accuracies (3 scores). Sum of these correlations for the rest of the fragments, both without normalisation and normalised (2 scores). Cosine similarity measure (itself and to* *power 3) between the predicted and measured intensities of the top 6 fragments, weighted by the squared values of the smoothed "best" fragment elution curve (2 scores). Relative intensities of the top 6 fragments (6 scores). Pearson correlations of the smoothed elution profile of the "best" fragment with the MS1 elution profiles corresponding to the peptide featuring 1, 2 or 3 C13 (3 scores).*" A caution is not so much the redundancy in these features (e.g., the C12 peak almost always implicates a corresponding C13 peak), it's that such extracted features are obscured at low abundance with high background. For omics analyses, where identification number versus FDR is the training objective, it is reasonable to extract the familiar, observed features. They are the more prominent signals.

Regarding a future program of machine learning that digs into the noise, one anticipates data input to include less of the information-contracted processed data (e.g., Pearson correlations) and more primary data such as local ion maps. Sets of MS and MS/MS spectra covering the full m/z range over LC peak elution timescales are a logical extrapolation, but training and inference times are not trivial considerations. As it has evolved with LLMs, it's not hard to imagine a few institutes developing large scale models and then using distillation techniques (e.g., rank reduction) to distribute smaller models for general use including further training for site-specific instruments and tasks. Complementing this trend is the explosive development of AI assisted coding tools. Further development in MS data analysis is anticipated in the short term. Poisson analysis would serve as a preprocessing step to connect sequence information with raw data for more computationally intensive training and inference.

The current paper addresses key features of structural biology and immunology related to p53 antigen presentation where the MS is enabling rather than the primary focus. This paper does not provide a statistical study comparing our Poisson with existing workflows as the targeted approach doesn't immediately lend itself to FDR characterization given that true identifications are few. Validation of p53 detections relevant to the focus of the paper follows from successfully identifying peptide shifts when mutant TP53 transductions replace wtTP53 transduction (Extended Data 2) and from concordant identifications in tumours and normal cells (Extended Data 1). One could argue that p53 presentation is generally at a high level and comparatively easy to detect in general. This will be addressed following the outline of the theory but there is some relevant literature. A 2022 MS immunopeptidomics study reported on HLA-A*02:01 restricted p53 peptides[6]. This study was a significant effort by a well-established group. TP53 null osteosarcoma cells, SaOS-2, were transduced with R273H or R175H mutTP53 and immune peptidomes isolated from a half billion cells of each transduction were analyzed. The protocol involved sequential BB7.2 (A02-specific) then W632 affinity isolation, fractionation and DDA LC-MS of the fractions. At an FDR of 5% 10 to 20 thousand peptides were identified (Fig. 1). The detection of p53 peptides was characterized: "*Nine and seven unique p53-specific peptides were found in SaOS-R175H and SaOS-R273H, respectively (Supplementary Table S7). Of note, unique peptides proximal to the reciprocal mutation were found in the two transfectants, i.e., P53264–272 (LLGRNSFEV) was found only in SaOS-R175H transfectant, whilst peptide P53164–172 (KQSQHMTEV) was found only in SaOS-R273H transfectant.*" Two points are interesting. The loss of the LLGRNSFEV peptides was also seen in our data with the R273H mutation (Extended Data 2c). Supplementary Table S7 contained results only for the R273H mutation and BB7.2 affinity purification. There were six distinct p53 peptides identified with two additional peptides being methionine oxidized

**Table 1.** Number of HLA class I peptides (8–15 mers) identified across transfected SaOS cell lines.

| Condition | HLA-A2 (BB7.2) Peptides | Pan HLA (W632) Peptides | Total |
|---|---|---|---|
| No vector control | 2688 | 8982 | 11,670 |
| R175H | 9127 | 9814 | 18,941 |
| R273H | 2970 | 7145 | 10,115 |

*Figure 1. Published table showing deep coverage of immune peptidomes for SaOS-2 cells transfected with mutTP53.*

| Peptide | core | icore | EL-score | EL_Rank | BA-score | BA_Rank | Aff(nM) | NB |
|---|---|---|---|---|---|---|---|---|
| TYSPALNKM | TYSPALNKM | TYSPALNKM | 0.001 | 17.7226 | 0.0393 | 49.4795 | 32689.87 | 0 |
| EYLDDRNTF | EYLDDRNTF | EYLDDRNTF | 0.0004 | 25.8049 | 0.0208 | 72.2677 | 39917.71 | 0 |
| GLAPPQHLI | GLAPPQHLI | GLAPPQHLI | 0.8114 | 0.1003 | 0.551 | 1.2706 | 128.74 | 1 |
| KQSQHMTEV | KQSQHMTEV | KQSQHMTEV | 0.5573 | 0.3262 | 0.5214 | 1.5557 | 177.43 | 1 |
| TYSPALNKMF | TYSPALNKF | TYSPALNKMF | 0.0001 | 40.25 | 0.0295 | 59.8287 | 36323.75 | 0 |
| GLAPPQHLIRV | GLAPPLIRV | GLAPPQHLIRV | 0.7515 | 0.1452 | 0.5838 | 1.013 | 90.32 | 1 |

*Figure 2. netMHCpan 4.1b binding predictions for HLA-A*02:01*

forms. Of these six, only GLAPPQHLIRV was shared with our data. Binding affinity calculations for HLA-A*02:01 for these six (Fig. 2) predicted 3 peptides were unlikely to bind. The study was not compromised; it focused on proteomics and immunopeptidomics associated with the TP53 mutations. The p53 detections were incidental to an omics analysis with deep coverage; the three likely falsely assigned sequences, giving the calculated bindings, reflect local FDRs which are much higher than the global 5% FDR. There is no literature support that p53 peptides, transfected or endogenous, are easy to detect.

Our approach to neoantigen detection is not conventional immunopeptidomics. As the task is targeted, albeit with hundreds of targets, we exploit the commercial availability of pools of synthetic peptides at low cost (JPT). With synthetic pools, fragmentation and elution can be precisely characterized on our instruments and with our operating conditions. Given an imperative for sensitivity, we have synthesized 20 μm monolithic capillary columns with flow rates of ~5 nl/min using laser pulled, distal coated, 10 μm capillary electrospray tips. Fine tip geometry at low flow is critical for high sensitivity, and it has been our experience that tip performance degrades much faster than the columns. Laser pulling conditions alone have not reproducibly generated optimal tips; instead, after pulling and coating, the tip and column are coupled with a union (PCU) and connected to the LC. Running an open split at operational pressure, the flow is measured under a stereo microscope using a laser pulled 50 μm capillary to collect flow by capillary action. The flow is established by inserting the tip, still connected to the column, in 48% HF for 1-2 minutes and remeasuring. When an appropriate flow is obtained, this becomes the operational LC configuration. Sensitivity also compels an optimal use of the ion beam given our quadrupole TOF geometry. DIA window widths are varied over m/z 350 – 700 so that the ion current is evenly distributed, given the characteristic m/z distribution of MHC class I peptides. In contrast to timsTOF studies of immune peptidomes[3], we rarely see dominant singly-charged peptides either from cells or synthetics; hence we do not target MS/MS windows for single charge states. Our current use of 11 MS/MS minimally overlapped windows results in complex fragment backgrounds, motivating the interest in stochastic sampling measures. Where cells can be counted, the LC-DIA MS runs start with no more than 1 million cells that are lysed, peptide-HLA class I complexes captured on antibody (W6/32) coated beads, acid released and extracted on C18 tips. Eluted peptides are loaded on a 50- μm ID monolith pre-column in the same format as tumour samples are processed. Pointedly, data presented is not of 1 million cell equivalents from a large-scale affinity purification loaded directly on column.

**Metric and Probabilistic Similarity.** Presently, the dominant approach to identifying a reference MS/MS spectrum embedded in sample DIA data is to use a Euclidean framework. An XIC approach identifies fragment peaks in the appropriate DIA window that co-elute with a precursor ion. Fragment and precursor peaks are compared with each other by Pearson correlations which are the inner product (Euclidean metric) of the normed elution profile vectors. This approach might be most effective when a spectral library is not present and the XIC traces are clean enough to extract a sequence tag. Extracting peptide sequences with both a matching precursor m/z and subsequence tag from a protein database, followed with NN prediction of MS/MS spectra, translates the analysis into spectrum-spectrum matching. If, on the other hand, one already had a reference spectrum one could generate a detection chromatogram by spectrum-spectrum matching at every scan. Then the peaks in this chromatogram can be further qualified by XIC traces. This is very intuitive given clean XIC traces but not so clear when individual XICs are obscured. Consider spectrum-spectrum matching by inner product or a function of the inner product using MS/MS fragments shown in Figs 3 and 4. The reference MS/MS spectrum defines the dimensionality of the vector with the orthogonal basis or coordinate vectors identified with the fragment m/z's and the coordinate magnitudes as the peak amplitudes of the fragments ($y_1$, $y_2$, …, $y_6$) (see Fig. 3). The peak amplitudes at the corresponding m/z's in the DIA data (Fig.

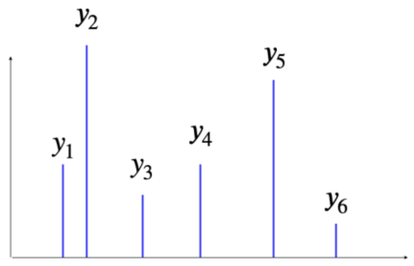


*Fig.3 Reference MS/MS spectrum*

4) generate the vector ($x_1$, $x_2$, …, $x_6$) which is then compared to the reference vector. The 'spectral angle' is defined by extension of a familiar geometric picture to arbitrary dimensions: $\left(\vec{x},\vec{y}\right) = \left|\vec{x}\right|\left|\vec{y}\right| cos(\theta)$. The direct use of the spectral angle $\theta(t) = \cos^{-1}\left[\frac{(\vec{x},\vec{y})}{|x||y|}\right]$ is favored for DDA because it compares 'shapes' without respect to data signal amplitude. However, generating a $\theta\left(t\right)$ detection chromatogram in a DIA setting with a complex fragment background, ignoring the overall fragment amplitudes comes at a cost in sensitivity/specificity. In these circumstances an inner product chromatogram $g(t) = \sum_{i=1}^{n}(x_i(t), y_i)$ with reference $\left|\vec{y}\right|$ = 1 is more effective. But not to belabor the details, it is a spatial intuition encapsulated in $\left(\vec{x},\vec{y}\right)$ that relates the patterns of the data and reference vectors in these detection algorithms. And this is a problem. Consider at a time point in the DIA data the MS/MS spectrum of Fig. 4 is extracted as a candidate for identification as the target characterized by Fig. 3. As suggested by the figures, the data spectrum is a scaled copy of the reference spectrum ($x_i = Ny_i$) except for the reference $x_3$ fragment whose amplitude in the data spectrum is zero. $\left(\vec{x},\vec{y}\right) = x_1y_1 + x_2y_2 + x_4y_4 + x_5y_5 + x_6y_6$ is the data spectrum's score which differs from the reference Fig. 3 by $x_3y_3$, a minor term relative to the sum of the other components. However, if one manually inspected these spectra the likelihood that the spectrum of Fig. 4 would be taken as the reference in Fig. 3 would depend on the signal amplitudes of the peaks in Fig. 4. With sufficient amplitude, one would conclude the absence of the expected $x_3$ peak invalidates the data spectrum as the target. The inner product does not identify this confident negative detection.

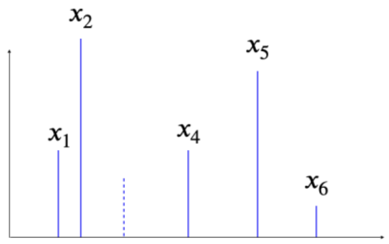


*Fig.4 Data MS/MS spectrum*

Broadly, pattern recognition in the setting of DIA detection is better treated as a stochastic sampling process and not as a time-dependent elution of shapes, although, as previously discussed, both feature sets can be generated and used in algorithmic classifiers. For counting ion events over a period that is short relative to the chromatographic time scale one expects the number of ion events in each m/z channel to be proportional to the counting period. In our experimental data the fragment ions are collected for 250 milliseconds while LC elution peaks are variable but in the range of 15 to 30 seconds. The arrival events are randomly but uniformly distributed in time over the counting period with an expected rate of ion arrivals. This stochastic event counting is sampling a Poisson process. In the MS/MS spectrum one has measured ions arriving in other m/z channels in this period, again arriving at random but uniformly distributed times, with each fragment channel characterized by an arrival rate. Sampling these channels would asymptotically converge on a shape (the reference spectrum) but the practical problem is not how similar the data and reference spectra as shapes are but to determine the likelihood that a measured distribution of events arises from a multi-channel Poisson process characterized by a set of arrival rates *given the events measured*.

Consider repeated measurement of a single channel Poisson process with an expected arrival rate L. The Central Limit Theorem states the distribution of these measured results converges on a normal distribution. For sampling the Poisson process the normal distribution of these measurements has both a mean *and a variance equal to L*. That is, high event counts in the measured MS/MS spectrum also have high variance. With the inner product between a measured and reference spectrum as a detection signal, highly fluctuating terms are summed with lower amplitude terms with less fluctuation, cancelling the significance of the low amplitude peaks. In applications using the inner product as a detection parameter, additional features such as the rank order of the individual fragment XIC elution profiles and the time correlation of the extracted ion chromatogram (XIC) traces with each other and with the precursor ion elution profile can be used in generating a feature-dependent scoring metric for qualifying detection candidates. As will be illustrated in the following, for sensitive detection with DIA data, the features of individual fragment XICs are largely obscured. For intense signals, tracing of the individual fragment XICs instead of the composite signal over all fragments (like the inner product) is self-evident.

**Poisson Detection Formulas.** Set the reference ion arrival rate in the ith channel as $\alpha_i$. To construct a normalized distribution { $p_i$ } over the D channels define an arrival period T such that $\sum_{i=1}^{D} T\alpha_i = \sum_{i=1}^{D} p_i = 1$. This rescales the unit of time so that in a period T, one ion fragment arrives and is distributed among the D channels with probabilities { $p_i$ }. In a collection period of NT, the expected number of ion fragments counted is N. The probability of measuring $n_i$ of the N events in the ith channel is given by the Poisson distribution P($n_i$) = $\frac{1}{n_i!}(Np_i)^{n_i}e^{-Np_i}$. Arrival events in the D channels are characterized as independent. Microscopically, charge conservation and/or the dynamics of molecular dissociation would have the appearance of one fragment at the expense of another. However, the period in which the instrument collects ion events for a m/z channel is far outside the time resolution required to measure such correlations between different m/z fragments. Each m/z channel is characterized only by the expected ion arrival rate. Under these conditions the probability of measuring a data MS/MS spectrum with events $\vec{n} = (n_1, n_2, \dots, n_D)$ that arise from a reference distribution $N\vec{p} = N(p_1, p_2, \dots, p_D)$ is given as the product of individual channel probabilities (Eqn. 1)

$$P(n_1, \dots, n_D \parallel N\vec{p}) = \prod_{i=1}^{D} \frac{1}{n_i!}(Np_i)^{n_i}e^{-Np_i} \qquad \text{(Eqn. 1)}$$

Previously it was noted that the inner product of Fig. 3 and Fig. 4 differed only in the $x_3y_3$ = 0 term from the inner product of Fig. 3 with itself (a perfect match). The difference was a small fraction of the inner products, irrespective of the total events in Fig. 3. For the Poisson detection from Eqn. 1 the difference would be a factor P($n_3$ = 0) = $\frac{1}{0!}(Np_3)^0e^{-Np_3} = e^{-Np_3}$. Here the expected reference events $y_3 = Np_3$ would determine the likelihood of the spectrum of Fig. 4. That is, the observed peaks in Fig. 4 are expected to follow the expected reference events $x_i = y_i = Np_i$. Suppose $x_2$ = 100 is observed. From the reference pattern of Fig. 3 one has (roughly) $p_3 = p_2/4$ so one would expect 25 = $Np_3$ events. With zero $x_3$ events the probability of this spectrum being a sampling of Fig. 3 drops by exp(-25). What the inner product scores as a nearly similar shape could be a very improbable match.

**Ion background and relative entropy.** Suppose in comparing a data spectrum with a reference spectrum, the measured events are greater than the expected number of events for a given channel, i.e., $n_i > Np_i$. For an isolated Poisson process this should decrease the probability P($n_i$). In real world DIA data, interpreting $n_i > Np_i$ as a sampling fluctuation is not workable due to the expected high level of background ions. Instead, the Poisson probability is calculated only for channels where $n_i < Np_i$. For $n_i > Np_i$, P( $n_i$ ) is set to 1. To detect a target $\vec{p}$, one increases the N that scales the reference spectrum and uses Eqn. 1 and the condition $n_i < Np_i$ to calculate if the data $\vec{n} = (n_1, n_2, \dots, n_D)$ can explain $N\vec{p} = N(p_1, p_2, \dots, p_D)$ events at a set cutoff probability ($P_c$). That is, as N increases more fragment channels have $n_i < Np_i$ and the the probability in Eqn. 1 decreases. The N such that $P(n_1, \dots, n_D \parallel N\vec{p}) < P_c$ is then the score of the 'Poisson' metric. The Poisson score for a reference pattern is then calculated as a function of elution time (scan number) to generate a chromatogram and this is plotted against the XIC of the precursor m/z.

As an algebraic exercise one can show $ln\left[P(n_1, \ldots, n_D \parallel N\vec{p})\right]$can be written in a form of a level-2 entropy (also known as a relative or Kullback-Leibler entropy). This derivation and the use of a Poisson or entropic measure for targeted detection with MS3 data has been published[7] (see Supplemental there).

**Automated processing of target sets and the hit file.** Synthetic retention time (RT) peptides are added to both the DDA and DIA runs of the synthetic target set and to the sample LC-DIA MS run. The DIA run of the synthetic set is used to validate the assignments of the DDA run. A polynomial mapping of elution times of the RT set from the synthetic run into the sample run is used to map the elution times of all the synthetic peptides in their DIA run into their expected elution times in the sample DIA run. For each targeted peptide (a candidate for detection), precursor XIC and Poisson chromatograms are generated. Chromatogram peak times (in scans) and amplitudes are extracted. The candidate's elution is associated with the highest Poisson prominence that has a nearby XIC prominence. The scan positions of the candidate peaks ('P sc', 'Xic sc'), the prominences (baseline corrected amplitudes, 'P pr', 'Xic pr'), the predicted elution position of the peptide ('pr sc'), the overall rank of the assigned Poisson peak ('rank') and the number of peaks in the Poisson chromatogram with an amplitude greater than a set fraction (here 1/2 or '#>0.5') are listed. For convenience in sorting, some derivative values are calculated; the positive distance between the predicted elution position and the position of the Poisson peak ('el dif'), as well as ratio of the Poisson to XIC prominence ('P/X pr') are listed. These features are sorted along columns and for those features that fall out of range, the corresponding rows are deleted. After the eliminations, a final sequence of sorting then clusters the best candidate detections near the top of the table. In Table 1 the final sort was by the Poisson significance

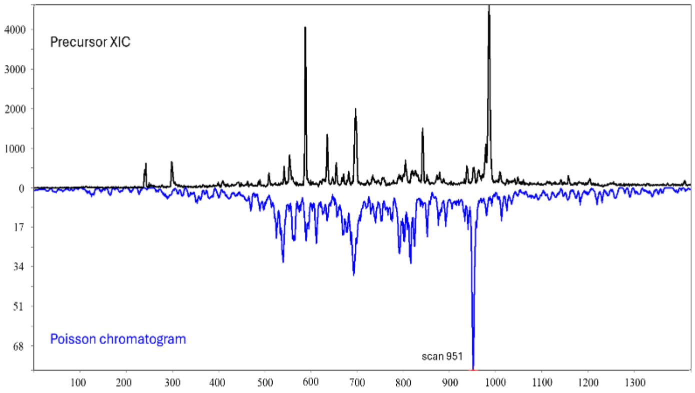


*Figure 5. Poisson Detection Pair*

| Label | peptide seq | m/z | P sc | P pr | Xic sc | Xic pr | pr sc | #>0.5 | el dif | P/X prom | rank |
|---|---|---|---|---|---|---|---|---|---|---|---|
| #49942 | GLAPPQHLIRV | 400.9119 | 581 | 4236 | 581 | 18515 | 593 | 1 | 12 | 0.229 | 1 |
| #49942 | KLLPENNVL | 520.3109 | 534 | 1163 | 535 | 4285 | 529 | 1 | 4 | 0.271 | 1 |
| #49942 | RPILTIITL | 520.3473 | 1005 | 1129 | 1006 | 4466 | 999 | 1 | 5 | 0.253 | 1 |
| #49942 | LLGRNSFEV | 517.7851 | 615 | 289 | 616 | 1864 | 615 | 1 | 0 | 0.155 | 1 |
| #49942 | GLAPPQHLIRV | 600.8642 | 580 | 152 | 581 | 473 | 593 | 1 | 13 | 0.321 | 1 |
| #SYN | KTYPVQLWV | 567.3213 | 951 | 91 | 952 | 317 | 954 | 1 | 3 | 0.287 | 1 |
| #49942 | APAPAAPTPA | 432.2347 | 221 | 263 | 220 | 162 | 263 | 2 | 42 | 1.623 | 1 |
| #49942 | LSQETFSDL | 520.2508 | 597 | 102 | 596 | 1905 | 632 | 2 | 35 | 0.054 | 2 |
| #49942Mox | TRVRAmAIY | 366.2026 | 316 | 219 | 316 | 343 | 277 | 6 | 38 | 0.638 | 1 |
| #49942Mox | VTRVRAmAIY | 399.2254 | 371 | 196 | 368 | 118 | 324 | 6 | 46 | 1.661 | 1 |
| #49942 | TPPPGTRV | 412.7349 | 224 | 90 | 223 | 216 | 187 | 6 | 36 | 0.417 | 2 |
| #49942 | FRHSVVVPY | 552.3035 | 535 | 80 | 534 | 2824 | 503 | 6 | 31 | 0.028 | 2 |
| #49942 | SDGLAPPQHLI | 574.3089 | 569 | 47 | 570 | 811 | 604 | 6 | 35 | 0.058 | 2 |
| #49942 | APAAPTPAAPA | 467.7533 | 310 | 105 | 311 | 387 | 305 | 7 | 4 | 0.271 | 1 |
| #49942 | RVLAMAIYKQ | 398.2339 | 519 | 77 | 519 | 119 | 508 | 7 | 10 | 0.647 | 1 |
| #49942 | STPPPGTRVR | 356.5367 | 218 | 173 | 218 | 146 | 171 | 8 | 46 | 1.185 | 2 |

*Table 1. Top-ranking peptides after sorting and elimination by XIC and Poisson chromatogram features.*

(‘>0.5’) column. The distance off the RT elution line is an important out of bounds eliminator, but the scatter of elution prediction inside a characteristic range is not a good top rank predictor. Typically (and here) hundreds of candidates are listed, but only the top rankings are relevant and shown.

**Inner product and Poisson chromatograms.** Table 1 lists the top-ranking candidates from an LC-DIA MS run of $10^6$ H2228 cells transduced with a p53 variant in which the cysteine at position 141 was replaced with tyrosine ($p53^{C141Y}$). Figure 6 shows the chromatogram picture for the top three ranking detections.

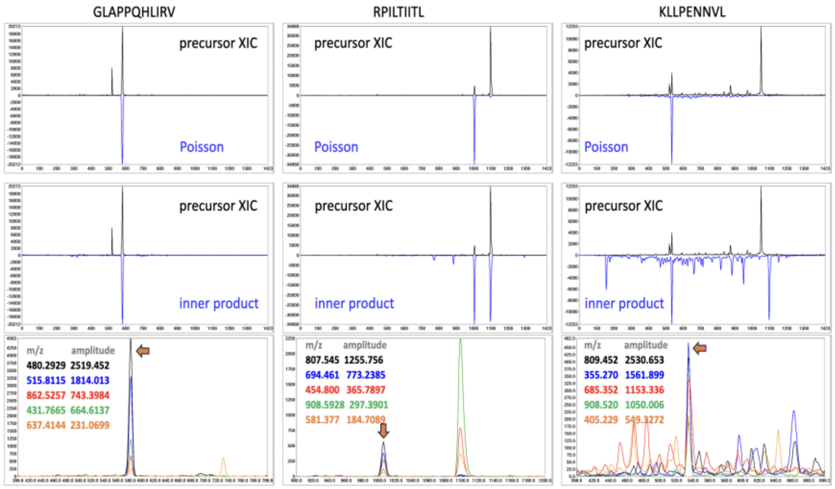


*Fig. 6 Comparing Poisson, inner product, and fragment XICs for top 3 rows of Table 1*

GLAPPQHLIRV, RPILTIITL, and KLLPENNVL are confidently detected by either Poisson, inner product, or individual fragment traces. While RPILTIITL has multiple peaks in the inner product assignment, detection can be confidently made by the set of individual fragment traces (Fig. 6, bottom panel row, marked with arrows). KLLPENNVL has a singular Poisson signature but multiple inner product peaks although its elution at scan 534 remains compelling by examining individual fragment XICs (Fig. 6). For KTYPVQLWV the Poisson chromatogram peak at scan 951 (Fig. 7, top left panel) is the largest peak but the total number of events that can be embedded (as a sampling of KTYPVQLWV’s fragmentation) at scan 951 is substantially smaller compared with LLGRNSFEV (Table 1, P pr’ column). The individual fragment XICs do not present correlated elution profiles but the Poisson chromatogram peak at 951 shows a significant sampling of KTYPVQLWV’s fragmentation profile is probable.

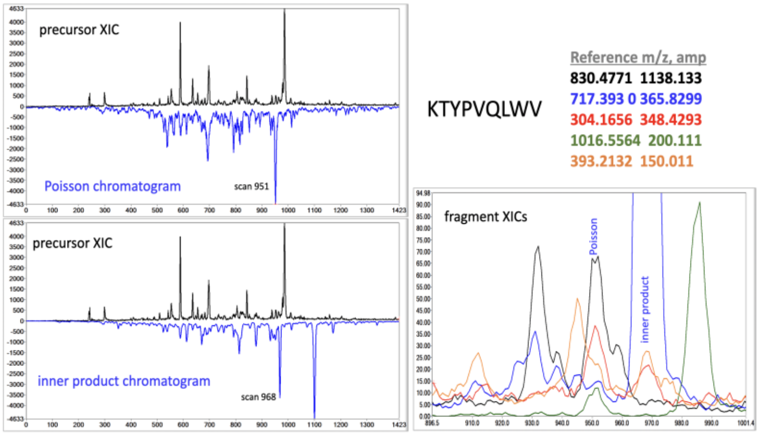


*Fig. 7 Comparing Poisson, inner product, and fragment XICs for detection of KTYPVQLWV, the $p53^{C141Y}$ peptide. The Poisson detection is at scan 951, the nearby inner product detection is at scan 968.*

As target abundance decreases relative to other fragment ions, the elution profiles of individual fragments are easily obscured. The inner product chromatogram peaks no longer effectively discriminate true and false detections and adding elution profile correlations between fragments doesn’t easily rescue this. Poisson detection is more discriminating than the inner product. It determines how many total arrival events can be embedded at an elution point if the relative arrivals are distributed among the fragment channels with a known distribution. As the reference distribution increases, the data channels in the measured spectrum will have too few events to be a probable (a cutoff parameter) fluctuation of the reference arrival rates. The Poisson amplitude, the highest number of embedding events for this channel set (a fragment spectrum) is this number. Rephrasing this, the inner product of the data channels on a normalized vector (the reference) is a geometric picture. Although weighted, events measured in one channel can compensate for a lack of events in another channel. To compensate, one needs to turn to individual fragment traces. As the signal-to-background ratio decreases, traces are obscured, and the inner product loses discrimination.

**False positives and false negatives.** False negatives may arise if there are chromatographic or chemical problems with the peptide. However, problems intrinsic to the peptide are also flagged by the synthetic set and here one draws not a negative detection but instead an inability to measure conclusion (Supplementary Discussion 2). If distributed events are embedded in the MS/MS data, there will be a corresponding Poisson amplitude. As target events decrease, background fragment ions appear in all the channels and these events compete with the true events for ranking in the hit list. The Poisson signature with its physically grounded requirements for event coincidence is more discriminating, but there are always detection limits. However, the dominant practical problem with Poisson detection is not false negatives, but false positives. Many of these are associated with partial overlaps due to fragment proximity in m/z and elution. To address this one turns to ion maps instead of XICs. We examine the top ranked subset of detections, operating with the premise that the Poisson chromatograms have eliminated most targets, and one is filtering out false positives.

**Three-dimensional Poisson detection.** Extending the framework of sampling a stochastic process in D channels where each channel has a known arrival rate, one can consider the elution and fragmentation of a target molecule as a stochastic process defined on a two-dimensional [m/z, t] domain. That is, the mass spectrometer records arrival events in multiple m/z channels at different scans. A domain covering the m/z and scan channels for a practical assessment of possible overlapping ion signals is defined, and for each peptide in a list (e.g., Table 1), ion maps for the precursor and each of the fragments are extracted from the LC-DIA MS data. The time (scan) coordinate is already common to all the maps. The m/z points of each fragment map are translated so that the center m/z of the fragment map is equal to the center m/z of the precursor ion map. Implicit in these translations and alignments is that the raw MS and MS/MS data points have been assigned to a uniformly spaced lattice. For each of the points in the [m/z, scan] domain the number of events that can be embedded in the D ion maps, given the reference distribution $N\vec{p} = N(p_1, p_2, \ldots, p_D)$, is calculated. The calculation is the same as with the Poisson chromatogram calculated over a set of XIC traces, only now one has a Poisson surface calculated over a set of ion fragment surfaces. That is, for each of the (m/$z_i$, $scan_j$) coordinate points, $\vec{n} = (n_1, n_2, \ldots, n_D)$ describes the measured events on the D fragment surfaces and one increases $N$ until $P(n_1, \ldots, n_D \parallel N\vec{p}) < P_c$ as before. This assigns N to (m/$z_i$, $scan_j$) and extended to all domain points creates an event amplitude or Poisson surface. The precursor ion map and Poisson surface are then compared for alignment over a two-dimensional [m/z, scan] domain by a surface Pearson correlation.

**Application to Primary T cells**. Extended Data Figure 1 shows the p53 detection data for stimulated T cells with HLA A*02:01, A*02:01, B*07:02, B*40:02, C*02:02 and C*07:02. Individual Poisson and XIC chromatograms for the peptides we identify as detected are shown in ED Fig. 1b. From all p53 reference patterns in this detection set, features are extracted and sorted with cutoffs and the top-ranking peptides by chromatogram features are listed in ED Fig. 1a, left panel. The blue shaded rows correspond to the detected p53 peptides. The three-dimensional analysis by Pearson correlation for the sequences in the top-ranked chromatogram list is shown in ED Fig. 1a, right panel, with illustrations of some of the surfaces in ED Fig. 1c and 1d. The resorting of the chromatographic list by surface Pearson correlations is automated but the software leaves a trail for insight as a manual evaluation. Figure 8 illustrates both chromatographic features and surface geometry. There are two yellow dotted lines embedded on the semi-transparent red surface. Cut at a scan, it is a local mass spectrum. Cut at an m/z it is a local precursor XIC trace. The Poisson chromatogram used in features ranking (ED Fig1a, left panel) is, in the local map, the solid white line embedded on the Poisson surface at the precursor's m/z. As with the inner product, there is no m/z associated with the Poisson chromatogram; it is an amplitude plotted along the scan direction, i.e., a chromatogram. Peaks extracted from the dotted yellow and solid white chromatograms nicely coincide. The surface perspective breaks this coincidence, and the comparatively poor Pearson surface correlation demotes VLAMAIYK's ranking (ED Fig1a, right panel). In extracting ion surfaces a domain is chosen that extends outside the m/z and elution intervals characteristic of the mass spectrometer and chromatography. Figure 9 links to a movie showing the precursor and Poisson detection surfaces for LLGRNSFEV. Ion peaks not in the relevant region around the center need to be suppressed for the Pearson correlation to accurately compare the precursor and Poisson surfaces. Instead of extracting a smaller region, an edge suppression function (roughly cylindrical with smoothed edges) defines the region over which the correlation is calculated (Figure 10). Keeping the extended domain as in the example of Fig. 9 provides better objects for manual inspection without impacting the ranking.

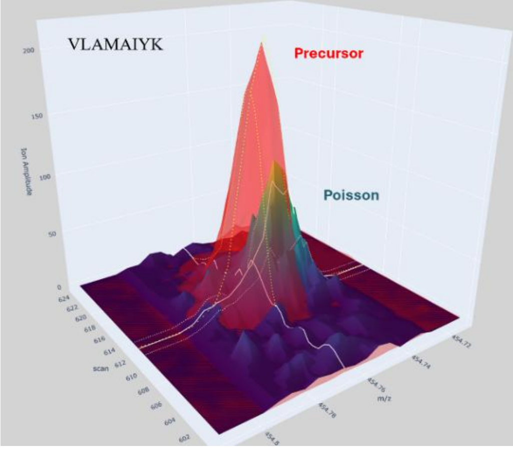


*Figure 8. Unlike chromatograms, the three-dimensional surfaces illustrate the discordance between precursor and Poisson surfaces.*

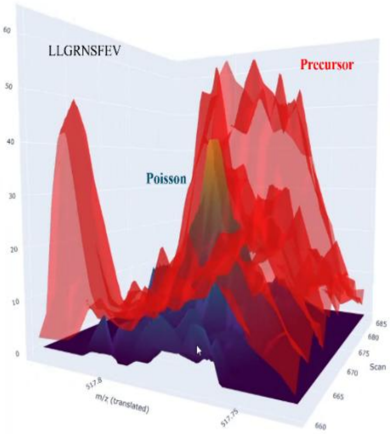


*Figure 9. Link to three-dimensional precursor and Poisson surfaces for LLGRNSFEV (Sup. Movie 2).*

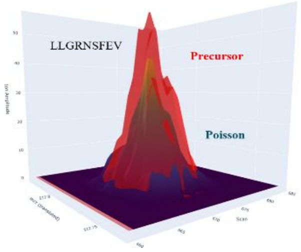


*Figure 10. Link to edge suppressed surfaces of Fig. 9 (Sup. Movie 3).*

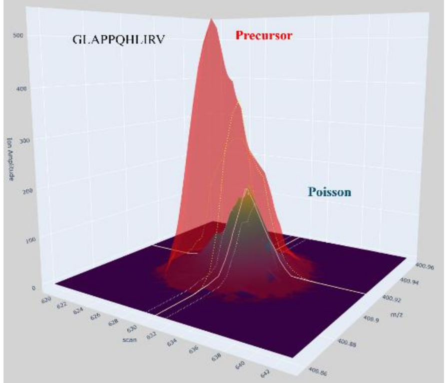


*Figure 11. Precursor and Poisson surfaces the dominant GLAPPQHLIRV p53 peptide. This is also a link to a movie (Sup. Movie 4) showing the precursor decomposed into two Gaussians.*

A three-dimensional perspective also allows decomposition of overlapping ion peaks which would be difficult to discern from chromatograms. ED Fig. 1a shows the relative rank of GLAPPQHLIRV is reduced when surface features (right panel) are used for ranking as opposed to chromatographic features (left panel). As a component of the surface Poisson analysis, software also decomposes the precursor ion surfaces in the chromatographic list file into two Gaussian surfaces by constrained nonlinear fits. Figure 11 shows the precursor and Poisson surfaces and it links to a movie showing the decomposition into two Gaussian surfaces (red and green, both partially transparent) and the Poisson surface, solid with yellow-to-blue shading. By visual inspection the Poisson surface is clearly correlated with the minor Gaussian (red) component and the Pearson calulation confirms this (Fig. 12). The decomposition is most interesting if the minor component shows good correlation where the major component does not. When the precursor is not well modeled as a composite of two Gaussian peaks the decomposition either generates a major component that correlates with the Poisson surface or fits a pair of Gaussians, neither of which is well correlated with the Poisson surface. KTZPVQLWV ranks highly by Pearson correlation with the precursor peak but poorly with either of the fit Gaussians. Figure 12 also shows KLLPENNVL, like GLAPPQHLIRV, having a better correlation with the minor component of a two Gaussian fit. Again, by visual inspection (Figure 13), an assymmetric peak in the precursor ion map (in semi-transparent red) is decomposed into two components, and the minor Gaussian from the two-component fit (in semi-transparent green) correlates with the Poisson surface (in solid yellow-to-blue shading). Figure 13 also links to a movie showing the spatial relations dynamically. These surfaces were not incorporated into the ranking of ED Fig. 1a, right panel, because the connections in the data don't work with the primitive sort and reject ranking. A utility is anticipated in the context of input features for adaptive classifiers that could be developed broadly in the MS community.

| Peptide | Precursor_mz | Pearson_Major | Pearson_Minor |
|---|---|---|---|
| RPILTIITL | 520.3473 | 0.911 | 0.4268 |
| RELNEALEL | 543.7931 | 0.898 | 0.5692 |
| GLAPPQHLIRV | 400.9119 | 0.416 | 0.889 |
| VLAMAIYK | 454.7673 | 0.6655 | 0.6673 |
| VLSPLPSQA | 456.2635 | 0.8245 | 0.3116 |
| LLGRNSFEV | 517.7851 | 0.8248 | 0.3947 |
| KTZPVQLWV | 565.805 | 0.3451 | 0.4471 |
| PPGTRVLAM | 471.2655 | 0.2705 | 0.1519 |
| KLLPENNVL | 520.3109 | 0.5038 | 0.7191 |

*Figure 12. Pearson correlations between Poisson surface and each of the nonlinear and constrained two-component Gaussian fits to the precursor ion surface.*

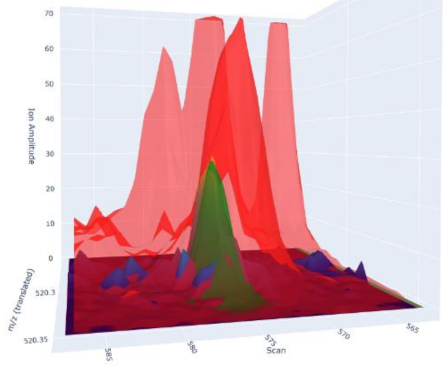


*Figure 13. Precursor ion map, minor Gaussian component and Poisson surface for KLLPENNVL (see text) also serving as a link to a movie (Sup. Movie 5) for a more intuitive perspective.*

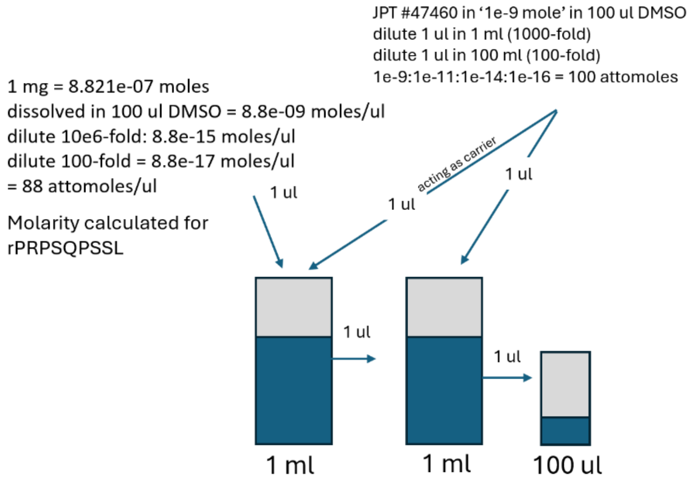


*Figure 14. Dilution protocol for sensitivity assay.*

Operating detection sensitivity combines sample processing, LC and MS instrumentation and data analysis. The mass spectrometer is a Sciex tripleTOF 6600 is a quadrupole-oTOF. Effective sample processing is evidenced by the main paper's central MS result, p53 detections across multiple 1 million cell inputs, and contrasting this detection with sample scales reported in the literature as discussed above. Although MS detection with purified synthetic peptides does not determine the sensitivity of detection framed as 'copy number per cell - the immunologically relevant target - it does parmetrize a significant scale. A less common aspect is the nano LC electrospray. From a prior ALK[8] study some years ago a set of 4 isotopically labelled and purified peptides (close to 1 mg each) had been stored at -20$^{o}$ C. A crude synthetic set of 265 ordered peptides was used as a carrier for the dilution protocol as shown in Figure 14. The crude peptides are roughly characterized at 100 attomole/ul concentration per peptide after dilution and, specifically for rPRPSQPSSL, 88 attomoles/ul. The other peptides were similar, correcting for differences in molecular weight. 1 ul was loaded by pressure bomb onto the precolumn, the columns reconnected to the LC and the gradient started. 43 minutes later the first peptides elute and data DIA MS data was acquired for 58 minutes (1127 scan cycles). Figure 15 shows the XIC and Poisson chromatograms. The LC resolution at the top of the gradient is comparatively poor for the alkane-modified, polystyrene-divinylbenzene monoliths as shown by the rPRPSQPSSL peptide and also addressed in Supplementary Discussion 2. Since the background is simple here, one can track just the precursor XICs. Overall, ion intensities are as expected, except for AMLDL(L*)HVA, which is substantially lower. Whether this reflects storage, synthesis or oxidation of the imidazole side chain or something else, is unknown.

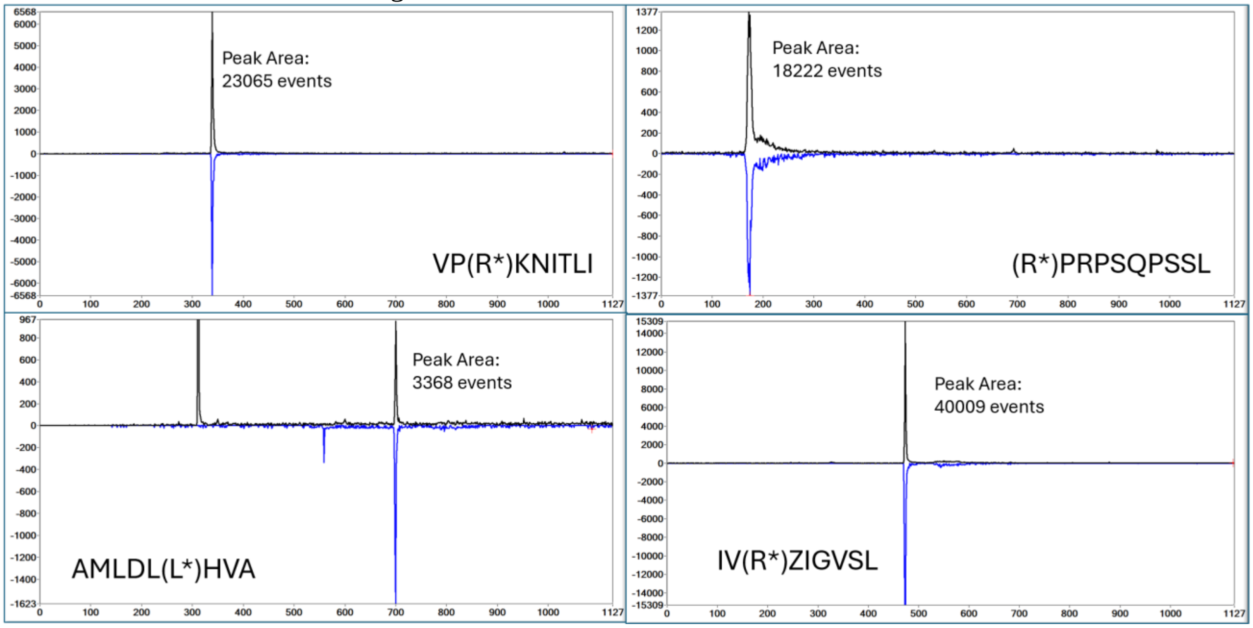


*Figure 15. Extracted ion and Poisson chromatograms for 4 peptides loaded at 88 - 100 attomoles each. '*' identifies isotope-labeled residue, 'Z' is alkylated cysteine.*
