## Supplementary material for "Cancers modulate p53 truncal neoantigen display to evade T cell detection": Supplmentary Discussion 2

**Supplementary Discussion 2.** Detection using synthetic peptides also identifies properties that preclude their sensitive detection.

One of the advantages of having crude synthetic peptide pools is a better understanding of the analytical MS methodology and its limitations. MS analyses can uncover signatures that reveal mechanistic reasons behind low sensitivity for detection. For example, one such signature is the dynamic oxidation of methionine in some peptides during the LC run. DDA/DIA analyses of synthetic pools containing GMNLRPILTI identified the unoxidized peptide eluting at scan 1050 but at comparatively low levels (**Fig. SD2a**). The oxidized peptide was also identified in the DDA run at much higher abundance and its fragmentation pattern gave an unusual (extracted precursor ion, Poisson detection chromatogram) pair. In particular, the oxidized peptide was observed at scan 853, broadly over scans 900 to 1000 and then a sharp scan coeluting with the unoxidized peptide (**Fig. SD2b**). The individual fragment ions, and their rank order in the reference pattern (black, blue, red, green) mark very high confidence detection (**Fig. SD2c**). The interpretation is that positive mode electrospray generates a cationized aerosol flux out of the needle tip and also solvated anions that must transfer their negative charge at the metal-coated distil end of the needle in our LC configuration. In the mobile phase, the excess of negative charge corresponds to a highly basic environment that promotes oxidation of methionine. We construe the triplet of GMoxNLRPILTI peaks in the following way. The 853-scan elution corresponds to GMoxNLRPILTI that was formed prior to LC-MS. The 1050 peak corresponds to the GMNLRPILTI being oxidized at the electrospray needle where the anion density is highest. The broad diffuse peak at scan 922 corresponds to oxidation at different positions in the column during the elution of GMNLRPILTI. Once oxidized, the peptide is more hydrophilic and migrates at a higher rate through the analytical column. Such facile oxidation of methionine during analysis is unusual in general for methionine containing peptides but was observed with all the p53 R248X synthetic peptides tested and hence diminishing sensitivity of peptide detection.

Poor LC focusing at the top of the gradient can also limit operating sensitivity (Fig. SD2d). Poisson detection chromatograms for the peptides HMTEVVRRZ and HMTEVVRHZ illustrate the poor LC elution profile for hydrophilic peptides using the alkane modified, polystyrenedivinylbenzene monolithic columns of our ultra-low flow (5 nl/min) chromatography. A scan here is 3 seconds, so the 100-scan elution profile corresponds to roughly 5 minutes peak width. This precludes achieving a relevant sensitivity for these two p53 peptides R175 and R175H, respectively.

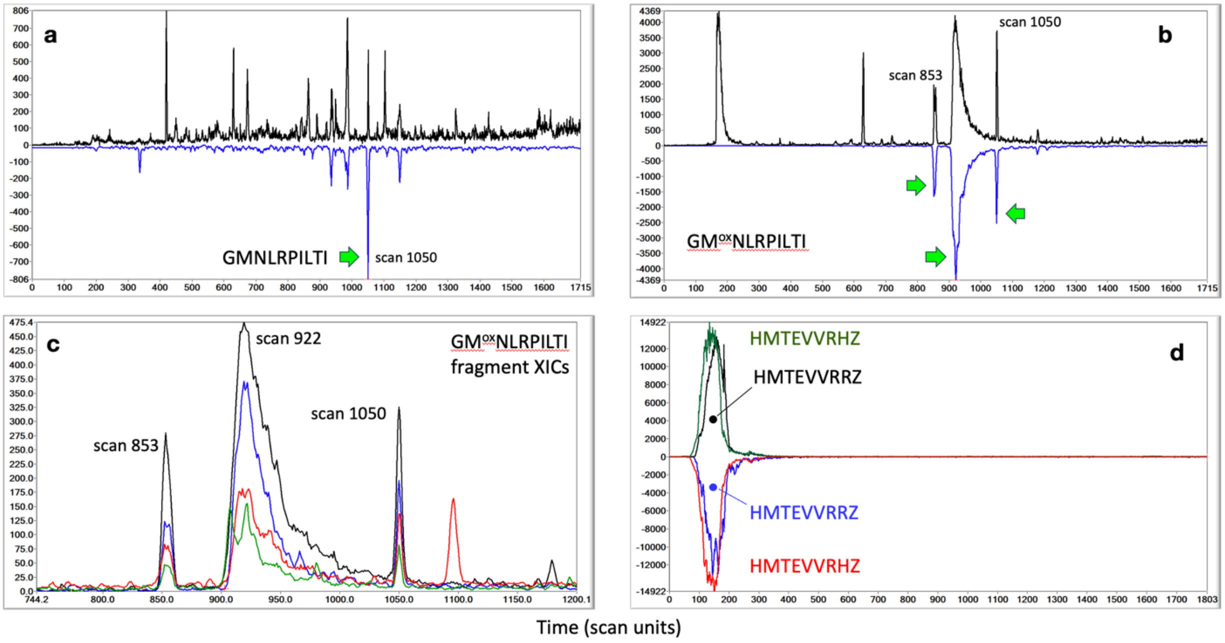


Figure **SD2a-d**. Poisson LC-DIAMS chromatograms for a set of synthetic peptides. **a**. Poisson detection chromatogram pair for the unoxidized GMNLRPILTI peptide showing a sharp elution at scan 1050. **b**. Poisson detection chromatogram pair for GMoxNLRPILTI. Here a sharp elution is observed early at scan 853, a broad diffuse and asymmetric elution profile covers scans 900-1000 and another sharp elution is observed at scan 1050. **c**. The 4 most abundant fragment XICs for GMoxNLRPILTI are plotted, validating the Poisson identification in panel b. **d**. Poisson detection pairs for the peptides HMTEVVRHZ and HMTEVVRRZ illustrate the poor LC elution profile and consequent poor sensitivity for these hydrophilic peptides
