## Supplementary material for "Cancers modulate p53 truncal neoantigen display to evade T cell detection": Supplmentary Discussion 3

**Supplementary Discussion 3.** Crystal Structures of p53 peptide and MHC complexes

1. WT p53 $^{187}$GLAPPQHLIRV$^{197}$/HLA-A*02:01 complex

There are two WT peptide/HLA-A*02:01 (HLA-A2) pMHC complexes per asymmetric unit that are nearly identical in structure. The LSQ (Least Squares Quadratic) alignments (Cα only) of their corresponding α-chains, β2m domains and peptides yields RMSD (Root Mean Square Deviation) values of 0.404Å, 0.324Å and 0.109Å, respectively. Thus, in the following description of the pMHC structure, only one complex (ABC chains of PDB 9OBF) is presented.

The 11mer WT p53 peptide $^{187}$GLAPPQHLIRV$^{197}$, has well-defined electron density maps and is bound to the HLA-A2 antigen-binding cleft using p2L and p11V acting as primary anchors. These are favored anchoring residues for A and F pockets of HLA-A2, respectively. The three N-terminal peptide residues p1G, p2L and p3A form multiple hydrogen bonds with neighboring MHC residues through their mainchain amide and carbonyl groups in addition to hydrophobic and Van de Waals interactions. The two consecutive prolines p4P and p5P are rare in the antigenic peptides displayed by HLA-A2 based on existing pMHC structures in the PDB. The peptide bulges up and out of the HLA antigen presenting platform groove from p4P to p8L. There are no specific interactions observed between p4P, p5P, p6Q, p7H and p8L and the HLA-A2 molecule, even including the p6Q that points to α2 helix. Water-bridged hydrogen bonds between these bulged residues and HLA-A2 may exist, however, since water molecules are not resolved in the structure at 3.2 Å resolution limit. Meanwhile, the five bulged residues, notably including two prolines that lack sidechains, leave the B, C and D pockets of HLA-A2 empty, thereby creating a cavity between the elevated peptide in this part and the peptide binding cleft below. The pocket below p5P and p6Q, seems to be occupied by a solvent molecule or a mixture of solvent molecules derived from buffers. The crystallization buffer includes a mixture of organic acids including malonic acid, citrate, succinic acid, DL-malic acid, acetate, format, and tartrate. In the structure, an ethylene glycol molecule was built into density for refinement purpose. The p9I points downward to the E pocket of HLA-A2 and serves as a secondary anchor of the peptide. The p10R projects upward, forming three hydrogen bonds to MHC including two contributed by its guanidinium group. Overall, p4P and p5P form a relatively flat featureless surface on the N-terminal region of peptide while p7H, p8L and p10R are the well-exposed, prominent residues of the peptide at its C-terminal region.

2. Mutant p53 $^{187}$GLAPPQHLFRV$^{197}$/HLA-A*02:01 complex

The mutant peptide/HLA-A*02:01 pMHC crystallized under the same condition as its WT with the same space group and nearly identical asymmetric unit cell parameters. The HLA-A2 molecule in the mutant structure has minimal structural change compared to WT. A LSQ alignment of WT and I195F complex α-chain resulted in a RMSD value of 0.242 Å (Cα atoms only) or 0.525 Å (all atoms). Even the residues that form the antigen binding cleft remain in the same conformation for the most part. However, the mutant peptide manifests striking conformational changes in comparison to its WT counterpart in several respects. While the five N-terminal peptide residues generally remain unchanged except p5P, which shifts slightly in response to the dramatic change of p6Q noted below, this N-terminal region conformational stability is not surprising due to the anchoring p2L residue and the rigidity of two consecutive prolines at P4 and P5. In contrast, the p6Q that points to the α2-helix in the WT complex shifts toward the α1-helix with its sidechain points upward and becoming well exposed. The p7H that is on α1-helix side rotates to α2-helix side and forces a large rotamer change of the sidechain of Q155 of HLA-A2 on the α2-helix to avoid a clash and to form a hydrogen bond between p7H and Q155. The new conformation of Q155 represents one of only a few significant conformational changes of HLA-A2 in the binding to the mutant peptide. The p8L that is fully exposed in the WT structure becomes completed buried, forming a secondary anchor in the mutant structure. From p5P to p8L a $3_{10}$ αhelical motif forms, which appears to help stabilize the conformation of mutant peptide in its bulged region. The residue p9F rotates upward in contrast to the buried p9I in the WT peptide complex to become exposed in the I195F mutant form. Interestingly, a likely chlorine ion bridges the amide group of p9F to two positively charged HLA-A2 residues, R97 and H114 from the peptide binding groove, thereby neutralizing the local charge and stabilizing peptide binding. The p10R largely remains in the same conformation as found in WT with its sidechain conformation locked by hydrogen bonding to the α1-helix of HLA-A2.

Overall, in the mutant peptide/HLA-A2 pMHC complex, the N-terminal region of the peptide remains largely the same as in the WT complex while its C-terminal region shows a completely different pattern with p6Q, p7H, and p9F and p10R forming a new cluster of exposed peptide residues. This dramatic antigenicity change can explain the differential immunogenicity of I195F/HLA-A*02:01. Given the locale of the change, it is likely to be prominently sensed by the β-chain of TCRs that respond to mutant I195F neoantigen and not the WT peptide.

3. WT p53 $^{249}$RPILTIITL$^{257}$/HLA-B*07:02 complex

There are two nearly identical WT peptide/HLA-B*07:02 (HLA-B7) pMHC complexes per asymmetric unit in structure. The LSQ alignments (Cα only) of their corresponding α-chains, β2m domains and peptides yields RMSD values of 0.294Å, 0.352Å and 0.165Å, respectively. Therefore, only one complex (ABC chains of PDB 9EIQ) is used for the presentation in this report.

In the WT structure, the p53 $^{249}$RPILTIITL$^{257}$ peptide is well defined. From its N-terminal region to C-terminal region, the 9mer peptide runs largely following the curvature of the peptide binding groove. The sidechain of the N-terminal p1R is sandwiched by the sidechains of R62 from α1-helix and W167 from α2-helix. The stacking of two parallel guanidinium groups and a guanidinium to an aromatic sidechain are commonly observed in protein structures and generally regarded as structurally stabilizing factors. The p1R also forms a bidentate salt bridge with E163 from the α2-helix. The p2P is a conserved residue in all HLA-B7-binding peptides based on known crystal structures deposited in PDB. The sidechain of p2P, a pyrrolidine ring is stacked onto the sidechain of Y7 from the bottom of the peptide-binding groove (not shown in figure). Overall, the N-terminal end of the peptide is tightly bounded to HLA-B7. The residue p3I points shallowly to the opening of D pocket. The exposed peptide residues are p4L, p6I and p8T while p5T points downwards and p7I sidewards to the E pocket. Both the B and C pockets on the α1-helix side are largely empty. The anchoring residue p9L is mostly favored for F-pocket binding of HLA-B7 based the known structures in PDB.

4. Mutant p53 $^{249}$MPILTIITL$^{257}$/HLA-B*07:02 complex

Like WT p53 $^{249}$RPILTIITL$^{257}$/HLA-B*07:02, the mutant p53 $^{249}$MPILTIITL$^{257}$/HLA-B*07:02 crystallized in the same space group and with practically equal cell parameters. Similarly, two nearly identical mutant peptide p1M/HLA-B7 complexes were found in one asymmetric unit. Though a few differences between two complexes are going to be discussed below, the complex comprised of ABC chains of PDB 9EJD will be used for general description and comparison.

The mutant structure is nearly the same as the WT. The LSQ alignments (Cα only/all atoms) of their corresponding α-chains, β2m domains and peptides yields RMSD values of 0.299/0.763 Å, 0.283/0.793 Å and 0.177/0.466 Å, respectively. These numbers indicate that the substitution of p1 residue on peptide causes only minor overall conformational changes. A few exceptional variations resulting from the mutation will be examined below.

In p1M/HLA-B7 structure, the replacing p1M dislodges the arginine pairing between R62 and p1R, and the sandwich-like stacking of p1R by R62 and W167 as well as the salt bridge between p1R and E163, all contributed by p1R in the WT structure. In the mutant p1R/M structure, only p1M is packed against W167 while R62 and E163 are seemingly unable to find their binding partners, showing two different unpaired conformations in two different p1M/HLA-B7 complexes. The alternative conformations indicate a destabilized binding of the N-terminal region of the mutant peptide in comparison to WT structure.

5. Mutant p53 $^{249}$SPILTIITL$^{257}$/HLA-B*07:02 complex

The crystals of the second mutant peptide p1S/HLA-B7 complex initially were obtained under the same condition as WT p1R/HLA-B7 and mutant p1M/HLA-B7 complexes. However, these crystals were small and poorly diffracted. Diffraction quality crystals were only obtained in an additive screening from a condition with an addition of 10% of 0.1M $ZnCl_2$ into initial crystal growth formulation. These crystals have a different space group. Inter-molecular bridging through

$Zn^{2+}$ ion present in the buffer seemingly helped form a more stable molecular packing for the improved diffraction limit.

There are also two nearly identical p1S/HLA-B7 complexes in one asymmetric unit of the new crystal. The LSQ alignments (Cα only/all atoms) of their α-chains, β2m domains and peptides results in RMSD values of 0.300/0.905Å, 0.175/0.699 Å and 0.100/0.245 Å, respectively. Therefore, the first complex (ABC chains of PDB 9EK4) is used for representation.

Additionally, the structure of the mutant p1S/HLA-B7 is almost identical to that of WT p1R/HLA-B7, the LSQ alignments (Cα only/all atoms) of their corresponding α-chains, β2m domains and peptides results in RMSD values of 0.240/0.727Å, 0.181/0.765 Å and 0.158/0.262 Å, respectively. The only notable conformational variation is associated with the p1R/S substitution. The sidechain of p1S forms an additional hydrogen bond to Y171. The amide group of the N-terminal residue of HLA-B7-binding peptide always forms a hydrogen bond to Y171 (not shown in the figures of WT and mutant p1M structures). The sidechain of W167 apparently tilts towards to the emptied space given that the long sidechain of the WT p1R has been replaced by p1S and forms a hydrogen bond to E163. Interestingly, E163 and R62 forms a long bidentate salt bridge (4.6-4.8 Å) in addition. The long salt bridge as well as other interactions around the p1S site described above is conserved in two p1S/HLA-B7 complexes in asymmetrical unit. The local structure surrounding P1 site seemingly reaches conformational stability through the adjustment described above. Furthermore, the absence of a long sidechain of p1R in WT creates a pocket with a small polar residue serine (p1S in the mutant) sitting at its bottom. This pocket presents an opportunity to host a long sidechain of a charged or a polar residue likely derived from CDR3α of an approaching TCR for ligand recognition.
