## Supplementary Data File 6 for "Cancers modulate p53 truncal neoantigen display to evade T cell detection"

**Supplementary Data File 6. p53$^{I195X}$ expression and HLA-A*02 expression in PANFR0583 and COV318 cells.**

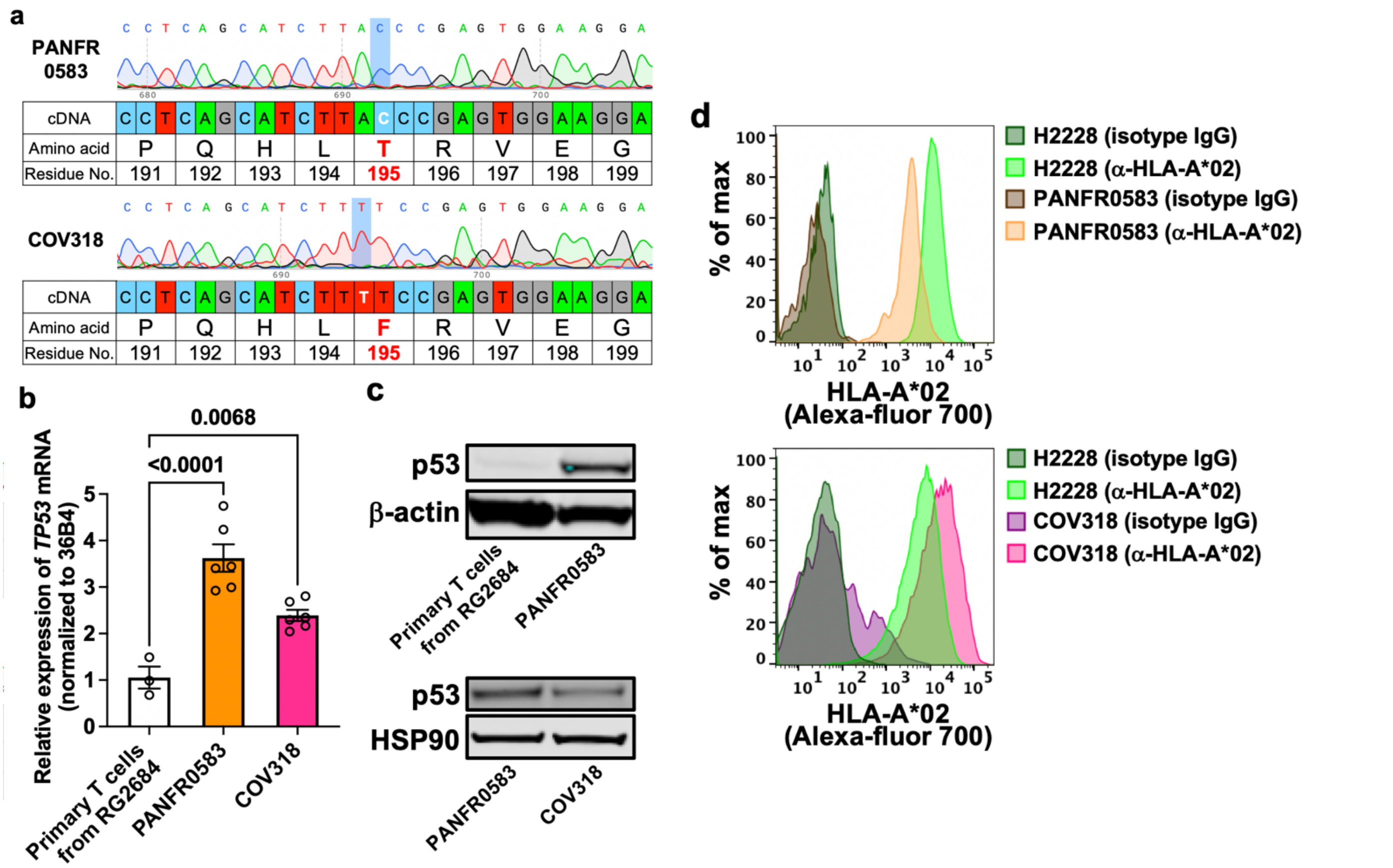


**a,** Sanger sequencing results of genomic DNA extracted from PANFR0583 organoid cells and COV318 cells.
**b,** *TP53* expression levels in PANFR0583 organoid cells and COV318 cells in comparison with those in activated primary circulating T cells from (a donor RG2684) in which wt $^{187}$GLAPPQHLIRV$^{197}$ peptides were clearly detected. Bars indicated the mean value and SEM. *P* values were calculated by Dunnett's multiple comparisons test. Data indicate technical replicates from one (RG2684) or two (PANFR0583 and COV318) independent biological replicates.
**c,** Immunoblotting results of p53 protein in PANFR0583 organoid cells and COV318 cells. p53 protein expression was significantly higher in PANFR0583 than in activated primary circulating T cells (RG2684) (top). p53 protein expression level in COV318 cells was comparable to that in PANFR0583 organoid cells. β-actin was used as internal control. Data are a representative of two biological replicates.
**d,** Flow cytometry data of HLA-A*02 cell surface protein in PANFR0583 organoid cells and COV318 cells. H2228 cells were used as positive control of HLA-A*02 expression. Isotype control antibodies were used for negative control. Data are a representative of two biological replicates.
