## Supplementary Data File 7 for "Cancers modulate p53 truncal neoantigen display to evade T cell detection"

**Supplementary Data File 7. Transduced HLA-B*07:02 cell surface expression in BT549$^{HLA\text{-}B*07:02}$ cells.**

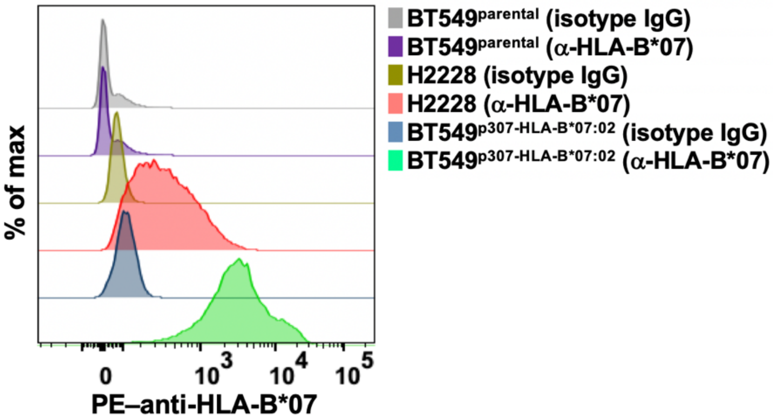


Flow cytometry data of HLA-B*07 cell surface protein on BT549 cells or BT549 transduced with HLA-B*07:02 using lentivirus system (p307 vector plasmid). H2228 cells were used as positive control of HLA-B*07 expression. Isotype control antibodies were used for negative control. Data are a representative of two biological replicates.
