## Supplementary Data File 9 for "Cancers modulate p53 truncal neoantigen display to evade T cell detection"

**Supplementary Data File 9. Flow cytometry data of transduced I195F-TCR on human pCD8 cells and J76 cells.**

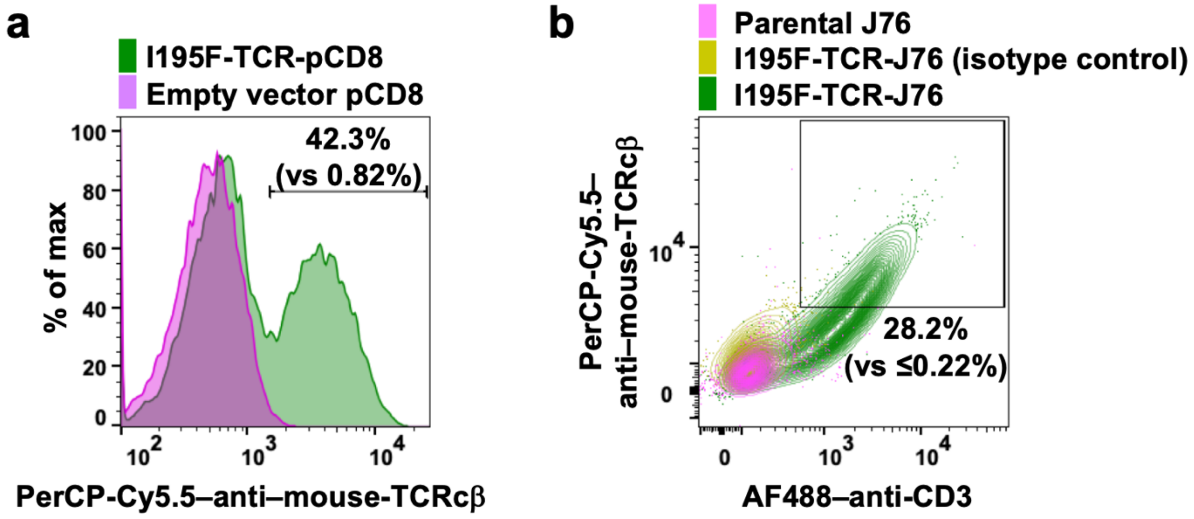


**a**,**b**, Flow cytometry data of mouse TCR β constant (mTCRcβ) region (**a**,**b**) and CD3 (**b**) on human pCD8 cells (**a**) and J76 cells (**b**). Transduced TCRs was engineered to have mouse TCR constant region, and mTCRcβ was utilized to label transduced TCR. Empty-vector–transduced pCD8 cells, parental J76 cells, and isotype control antibodies were used for negative control. Note that parental J76 cells cannot express CD3 on cell surface unless TCR is successfully transduced. Frequency (%) of mTCRcβ$^+$ (**a**) and mTCRcβ$^+$/CD3$^+$ (**b**) cells in overall I195F-TCR transduced cells are shown in comparison with the negative control. Data are representatives of two biological replicates.
