## Supplementary Data File 10 for "Cancers modulate p53 truncal neoantigen display to evade T cell detection"

**Supplementary Data File 10. ERAP1 allotypes in PANFRO0583 and COV318 cells.**

**a** (PANFR0583)

**Exon 2–9**

**Forward primer: GAAGATGGTGTTTCTGCCCC**

**Reverse primer: AAGGGATGGATGGCTTTTGC**

TTTCTACTTTCCTCACTGTTGGCTCTCTTAACTGTGTCCACTCCTTCATGGTGTCAGAGCACTGA
AGCATCTCCAAAACGTAGTGATGGGACACCATTTCCTTGGAATAAAATACGACTTCCTGAGTA
CGTCATCCCAGTTCATTATGATCTCTTGATCCATGCAAACCTTACCACGCTGACCTTCTGGGGA
ACCACGAAAGTAGAAATCACAGCCAGTCAGCCCACCAGCACCATCATCCTGCATAGTCACCAC
CTGCAGATATCTAGGGCCACCCTCAGGAAGGGAGCTGGAGAGAGGCTATCGGAAGAACCCCT
GCAGGTCCTGGAACACCCCCGTCAGGAGCAAATTGCACTGCTGGCTCCCGAGCCCCTCCTTGTC
GGGCTCCCGTACACAGTTGTCATTCACTATGCTGGCAATCTTTCGGAGACTTTCCACGGATTTT
ACAAAAGCACCTACAGAACCAAGGAAGGGGAACTGAGGATACTAGCATCAACACAATTTGAA
CCCACTGCAGCTAGAATGGCCTTTCCCTGCTTTGATGAACCTGCCTTCAAAGCAAGTTTCTCAA
TCAAAATTAGAAGAGAGCCAAGGCACCTAGCCATCTCCAATATGCCATTGGTGAAATCTGTGA
CTGTTGCTGAAGGACTCATAGAAGACCATTTTGATGTCACTGTGAAGATGAGCACCTATCTGGT
GGCCTTCATCATTTCAGATTTTGAGTCTGTCAGCAAGATAACCAAGAGTGGAGTCAAGGTTTCT
GTTTATGCTGTGCCAGACAAGATAAATCAAGCAGATTATGCACTGGATGCTGCGGTGACTCTTC
TAGAATTTTATGAGGATTATTTCAGCATACCGTATCCCCTACCCAAACAAGATCTTGCTGCTAT
TCCCGACTTTCAGTCTGGTGCTATGGAAAACTGGGGACTGACAACATATAGAGAATCTGCTCTG
TTGTTTGATGCAGAAAAGTCTTCTGCATCAAGTAAGCTTGGCATCACAATGACTGTGGCCCATG
AACTGGCTCACCAGTGGTTTGGGAACCTGGTCACTATGGAATGGTGGAATGATCTTTGGCTAA
ATGAAGGATTTGCCAAATTTATGGAGTTTGTGTCTGTCAGTGTGACCCATCCTGAACTGAAAGT
TGGAGATTATTTCTTTGGCAAATGTTTTGACGCAATGGAGGTAGATGCTTTAAATTCCTCACAC
CCTGTGTCTACACCTGTGGAAAATCCTGCTCAGATCCGGGAGATGTTTGATGATGTTTCTTATG
ATAAGGGAGCTTGTATTCTGAATATGCTAAGGGAGTATCTTAGTGCTGACGCATTTAAAAGTG
GTATTGTACAGTATCTCCAGAAGCATAGCTATAAAAATACAAAAAACGAGGACCTGTGGGATA
GT

**Amino acid sequence**

$^{15}$FLLSSLLALLTVSTPSWCQSTEASPKRSDGTPFPWNKIRLP**E**YVIPVHYDLLIHANLTTLTFWGTTK
VEITASQPTSTIILHSHHLQISRATLRKGAGERLSEEPLQVLEHP**R**QEQIALLAPEPLLVGLPYTVVIH
YAGNLSETFHGFYKSTYRTKEGELRILASTQFEPTAARMAFPCFDEPAFKASFSIKIRREPRHLAISN
MPLVKSVTVAEGLIEDHFDVTVKMSTYLVAFIISDFESVSKITKSGVKVSVYAVPDK**I**NQADYALD
AAVTLLEFYEDYFSIPYPLPKQDLAAIPDFQSGAMENWGLTTYRESALLFDAEKSSASSKL**G**IT**M**TV
AHELAHQWFGNLVTMEWWNDLWLNEGFAKFMEFVSVSVTHPELKVGDYFFGKCFDAMEVDAL
NSSHPVSTPVENPAQIREMFDDVSYDKGACILNMLREYLSADAFKSGIVQYLQKHSYKNTKNEDL
WDS$^{481}$

**Exon 11–19**

**Forward primer: TCATCCTCACATTGGCATCAG**

**Reverse primer: GGAGCCAAAACAGCCATCTC**

TGGACACTGCAGAAGGGTTTTCCCCTAATAACCATCACAGTGAGGGGGAGGAATGTACACATG
AAGCAAGAGCACTACATGAAGGGCTCTGACGGCGCCCCGGACACTGGGTACCTGTGGCATGTT
CCATTGACATTCATCACCAGCAAATCCGACATGGTCCATCGATTTTTGCTAAAAACAAAAACA
GATGTGCTCATCCTCCCAGAAGAGGTGGAATGGATCAAATTTAATGTGGGCATGAATGGCTAT
TACATTGTGCATTACGAGGATGATGGATGGGACTCTTTGACTGGCCTTTTAAAAGGAACACAC
ACAGCAGTCAGCAGTAATGATCGGGCGAGTCTCATTAACAATGCATTTCAGCTCGTCAGCATT
GGGAAGCTGTCCATTGAAAAGGCCTTGGATTTATCCCTGTACTTGAAACATGAAACTGAAATT
ATGCCCGTGTTTCAAGGTTTGAATGAGCTGATTCCTATGTATAAGTTAATGGAGAAAAGAGAT
ATGAATGAAGTGGAAACTCAATTCAAGGCCTTCCTCATCAGGCTGCTAAGGGACCTCATTGAT
AAGCAGACATGGACAGACGAGGGCTCAGTCTCAGAGCGAATGCTGCGGAGTCAACTACTACTC
CTCGCCTGTGTGCACAACTATCAGCCGTGCGTACAGAGGGCAGAAGGCTATTTCAGAAAGTGG
AAGGAATCCAATGGAAACTTGAGCCTGCCTGTCGACGTGACCTTGGCAGTGTTTGCTGTGGGG

GCCCAGAGCACAGAAGGCTGGGATTTTCTTTATAGTAAATATCAGTTTTCTTTGTCCAGTACTG
AGAAAAGCCAAATTGAATTTGCCCTCTGCAGAACCCAAAATAAGGAAAAGCTTCAATGGCTAC
TAGATGAAAGCTTTAAGGGAGATAAAATAAAAACTCAGGAGTTTCCACAAATTCTTACACTCA
TTGGCAGGAACCCAGTAGGATACCCACTGGCCTGGCAATTTCTGAGGAAAAACTGGAACAAAC
TTGTACAAAAGTTTGAACTTGGCTCATCTTCCATAGCCCACATGGTAATGGGTACAACAAATCA
ATTCTCCACAAGAACACGGCTTGAAGAGGTAAAAGGATTCTTCAGCTCTTTGAAAGAAAATGG
TTCTCAGCTCCGTTGTGTC

**Amino acid sequence**
$^{524}$WTLQ**K**GFPLITITVRGRNVHMKQEHYMKGSDGAPDTGYLWHVPLTFITSKS**D**MVHRFLLKTKT
DVLILPEEVEWIKFNVGMNGYYIVHYEDDGWDSLTGLLKGTHTAVSSNDRASLINNAFQLVSIGKL
SIEKALDLSLYLKHETEIMPVFQGLNELIPMYKLMEKRDMNEVETQFKAFLIRLLRDLIDKQTWTDE
GSVSE**R**MLRS**Q**LLLLACVHNYQPCVQRAEGYFRKWKESNGNLSLPVDVTLAVFAVGAQSTEGWD
FLYSKYQFSLSSTEKSQIEFALCRTQNKEKLQWLLDESFKGDKIKTQEFPQILTLIGRNPVGYPLAWQ
FLRKNWNKLVQKFELGSSSIAHMVMGTTNQFSTRTRLEEVKGFFSSLKENGSQLRCV$^{909}$

**b** (COV318)
**Exon 2–9**
**Forward primer: GAAGATGGTGTTTCTGCCCC**
**Reverse primer: AAGGGATGGATGGCTTTTGC**
CTACTTTCCTCACTGTTGGCTCTCTTAACTGTGTCCACTCCTTCATGGTGTCAGAGCACTGAAGC
ATCTCCAAAACGTAGTGATGGGACACCATTTCCTTGGAATAAAATACGACTTCCTGAGTACGTC
ATCCCAGTTCATTATGATCTCTTGATCCATGCAAACCTTACCACGCTGACCTTCTGGGGAACCA
CGAAAGTAGAAATCACAGCCAGTCAGCCCACCAGCACCATCATCCTGCATAGTCACCACCTGC
AGATATCTAGGGCCACCCTCAGGAAGGGAGCTGGAGAGAGGCTATCGGAAGAACCCCTGCAG
GTCCTGGAACACCCCCCTCAGGAGCAAATTGCACTGCTGGCTCCCGAGCCCCTCCTTGTCGGGC
TCCCGTACACAGTTGTCATTCACTATGCTGGCAATCTTTCGGAGACTTTCCACGGATTTTACAA
AAGCACCTACAGAACCAAGGAAGGGGAACTGAGGATACTAGCATCAACACAATTTGAACCCA
CTGCAGCTAGAATGGCCTTTCCCTGCTTTGATGAACCTGCCTTCAAAGCAAGTTTCTCAATCAA
AATTAGAAGAGAGCCAAGGCACCTAGCCATCTCCAATATGCCATTGGTGAAATCTGTGACTGT
TGCTGAAGGACTCATAGAAGACCATTTTGATGTCACTGTGAAGATGAGCACCTATCTGGTGGC
CTTCATCATTTCAGATTTTGAGTCTGTCAGCAAGATAACCAAGAGTGGAGTCAAGGTTTCTGTT
TATGCTGTGCCAGACAAGATAAATCAAGCAGATTATGCACTGGATGCTGCGGTGACTCTTCTA
GAATTTTATGAGGATTATTTCAGCATACCGTATCCCCTACCCAAACAAGATCTTGCTGCTATTC
CCGACTTTCAGTCTGGTGCTATGGAAAACTGGGGACTGACAACATATAGAGAATCTGCTCTGTT
GTTTGATGCAGAAAAGTCTTCTGCATCAAGTAAGCTTGGCATCACAATGACTGTGGCCCATGA
ACTGGCTCACCAGTGGTTTGGGAACCTGGTCACTATGGAATGGTGGAATGATCTTTGGCTAAAT
GAAGGATTTGCCAAATTTATGGAGTTTGTGTCTGTCAGTGTGACCCATCCTGAACTGAAAGTTG
GAGATTATTTCTTTGGCAAATGTTTTGACGCAATGGAGGTAGATGCTTTAAATTCCTCACACCC
TGTGTCTACACCTGTGGAAAATCCTGCTCAGATCCGGGAGATGTTTGATGATGTTTCTTATGAT
AAGGGAGCTTGTATTCTGAATATGCTAAGGGAGTATCTTAGTGCTGACGCATTTAAAAGTGGT
ATTGTACAGTATCTCCAGAAGCATAGCTATAAAAATACAAAAAACGAGGACCTGTGGGATAGT

**Amino acid sequence**
$^{16}$LLSSLLALLTVSTPSWCQSTEASPKRSDGTPFPWNKIRLP**E**YVIPVHYDLLIHANLTTLTFWGTTKV
EITASQPTSTIILHSHHLQISRATLRKGAGERLSEEPLQVLEHP**P**QEQIALLAPEPLLVGLPYTVVIHYA
GNLSETFHGFYKSTYRTKEGELRILASTQFEPTAARMAFPCFDEPAFKASFSIKIRREPRHLAISNMPL
VKSVTVAEGLIEDHFDVTVKMSTYLVAFIISDFESVSKITKSGVKVSVYAVPDK**I**NQADYALDAAV
TLLEFYEDYFSIPYPLPKQDLAAIPDFQSGAMENWGLTTYRESALLFDAEKSSASSKL**G**IT**M**TVAHE
LAHQWFGNLVTMEWWNDLWLNEGFAKFMEFVSVSVTHPELKVGDYFFGKCFDAMEVDALNSSH
PVSTPVENPAQIREMFDDVSYDKGACILNMLREYLSADAFKSGIVQYLQKHSYKNTKNEDLWDS$^{481}$

**Exon 11–19**
**Forward primer: TCATCCTCACATTGGCATCAG**
**Reverse primer: GGAGCCAAAACAGCCATCTC**

CAGAAGGGTTTTCCCCTAATAACCATCACAGTGAGGGGGAGGAATGTACACATGAAGCAAGA
GCACTACATGAAGGGCTCTGACGGCGCCCCGGACACTGGGTACCTGTGGCATGTTCCATTGAC
ATTCATCACCAGCAAATCCGACATGGTCCATCGATTTTTGCTAAAAACAAAAACAGATGTGCTC
ATCCTCCCAGAAGAGGTGGAATGGATCAAATTTAATGTGGGCATGAATGGCTATTACATTGTG
CATTACGAGGATGATGGATGGGACTCTTTGACTGGCCTTTTAAAAGGAACACACACAGCAGTC
AGCAGTAATGATCGGGCGAGTCTCATTAACAATGCATTTCAGCTCGTCAGCATTGGGAAGCTG
TCCATTGAAAAGGCCTTGGATTTATCCCTGTACTTGAAACATGAAACTGAAATTATGCCCGTGT
TTCAAGGTTTGAATGAGCTGATTCCTATGTATAAGTTAATGGAGAAAAGAGATATGAATGAAG
TGGAAACTCAATTCAAGGCCTTCCTCATCAGGCTGCTAAGGGACCTCATTGATAAGCAGACAT
GGACAGACGAGGGCTCAGTCTCAGAGCGAATGCTGCGGAGTCAACTACTACTCCTCGCCTGTG
TGCACAACTATCAGCCGTGCGTACAGAGGGCAGAAGGCTATTTCAGAAAGTGGAAGGAATCCA
ATGGAAACTTGAGCCTGCCTGTCGACGTGACCTTGGCAGTGTTTGCTGTGGGGGCCCAGAGCA
CAGAAGGCTGGGATTTTCTTTATAGTAAATATCAGTTTTCTTTGTCCAGTACTGAGAAAAGCCA
AATTGAATTTGCCCTCTGCAGAACCCAAAATAAGGAAAAGCTTCAATGGCTACTAGATGAAAG
CTTTAAGGGAGATAAAATAAAAACTCAGGAGTTTCCACAAATTCTTACACTCATTGGCAGGAA
CCCAGTAGGATACCCACTGGCCTGGCAATTTCTGAGGAAAAACTGGAACAAACTTGTACAAAA
GTTTGAACTTGGCTCATCTTCCATAGCCCACATGGTAATGGGTACAACAAATCAATTCTCCACA
AGAACACGGCTTGAAGAGGTAAAAGGATTCTTCAGCTCTTTGAAAGAAAATGGTTCTCAGCTC
CGTTGTGTCCAACAGACAATTGAAACCATTGAAGAAAACATCGGTTGGATGGATAAGAATTTT
GATAAAATCAGAGTGTGGCTGCAAAGTGAAAAGCTTGAACGTATGTAAAAATTCCTCCCTTGC
CAGGTTCCTGTTATCTCTA

**Amino acid sequence**
$^{527}$Q**K**GFPLITITVRGRNVHMKQEHYMKGSDGAPDTGYLWHVPLTFITSKS**D**MVHRFLLKTKTDVLI
LPEEVEWIKFNVGMNGYYIVHYEDDGWDSLTGLLKGTHTAVSSNDRASLINNAFQLVSIGKLSIEK
ALDLSLYLKHETEIMPVFQGLNELIPMYKLMEKRDMNEVETQFKAFLIRLLRDLIDKQTWTDEGSV
SE**R**MLRS**Q**LLLLACVHNYQPCVQRAEGYFRKWKESNGNLSLPVDVTLAVFAVGAQSTEGWDFLY
SKYQFSLSSTEKSQIEFALCRTQNKEKLQWLLDESFKGDKIKTQEFPQILTLIGRNPVGYPLAWQFLR
KNWNKLVQKFELGSSSIAHMVMGTTNQFSTRTRLEEVKGFFSSLKENGSQLRCVQQTIETIEENIGW
MDKNFDKIRVWLQSEKLERM*

**a,b,** Sanger sequencing results of cDNA extracted from PANFR0583 organoid cells (**a**) and COV318 cells (**b**) using primers shown. Homozygote was assumed for both samples based on the sanger sequencing electropherograms. Amino acid sequences are given according to their cDNA sequences. Codons and residues which affect ERAP1 function are highlighted in blue. PANFR0583 had the most active variant (allotype 2), and COV318 cells had the second most active variant (allotype 1) among 10 variants phenotyped by Hutchinson JP et al. (PMID: 33617882).
