## Supplementary Data File 12 for "Cancers modulate p53 truncal neoantigen display to evade T cell detection"

### Supplementary Data File 12: Original source data for immunoblotting

#### a For Fig. 5a

1 H2228[parental]

2 H2228[p307-p53-I195F]

3 COV318

4 PANFR0583

5 Jurkat E6-1 (as a positive control)

* Irrelevant samples

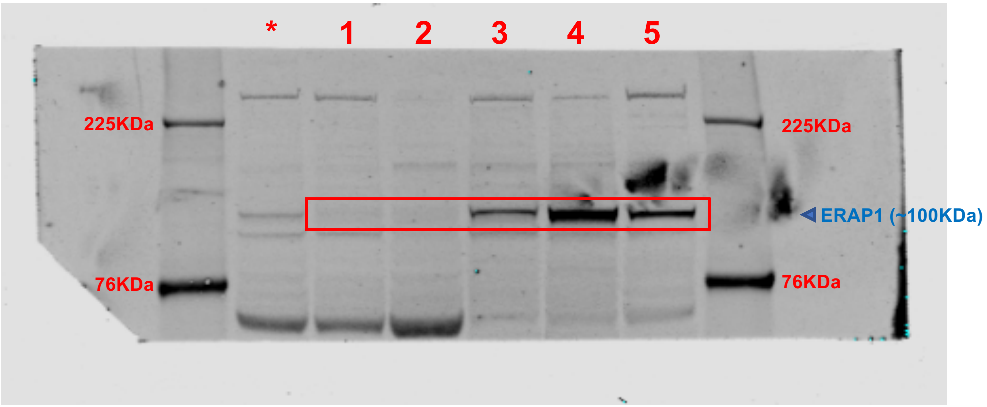


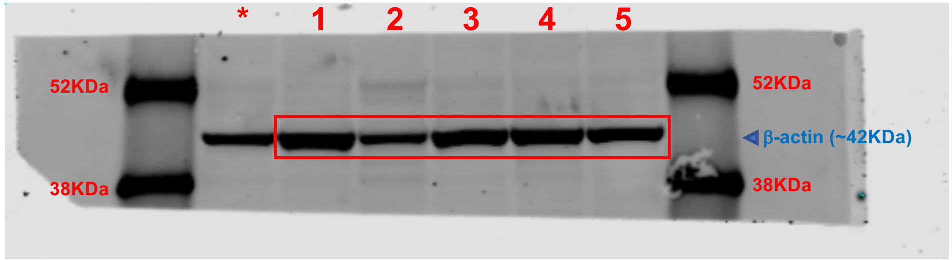


**Supporting data for Fig. 5a.**
Original source images for immunoblotting data used in Fig. 5a.
Two gels were derived from the same parent gel.
Red rectangles indicate where images were cropped for Fig. 5a.

Supplementary Data File 12: Original source data for immunoblotting

**b** For **Fig. 5b**

1 COV318sg-control

2 COV318sg1-ERAP1

3 COV318sg2-ERAP1

4 COV318sg4-ERAP1

5 COV318parental

* Irrelevant samples

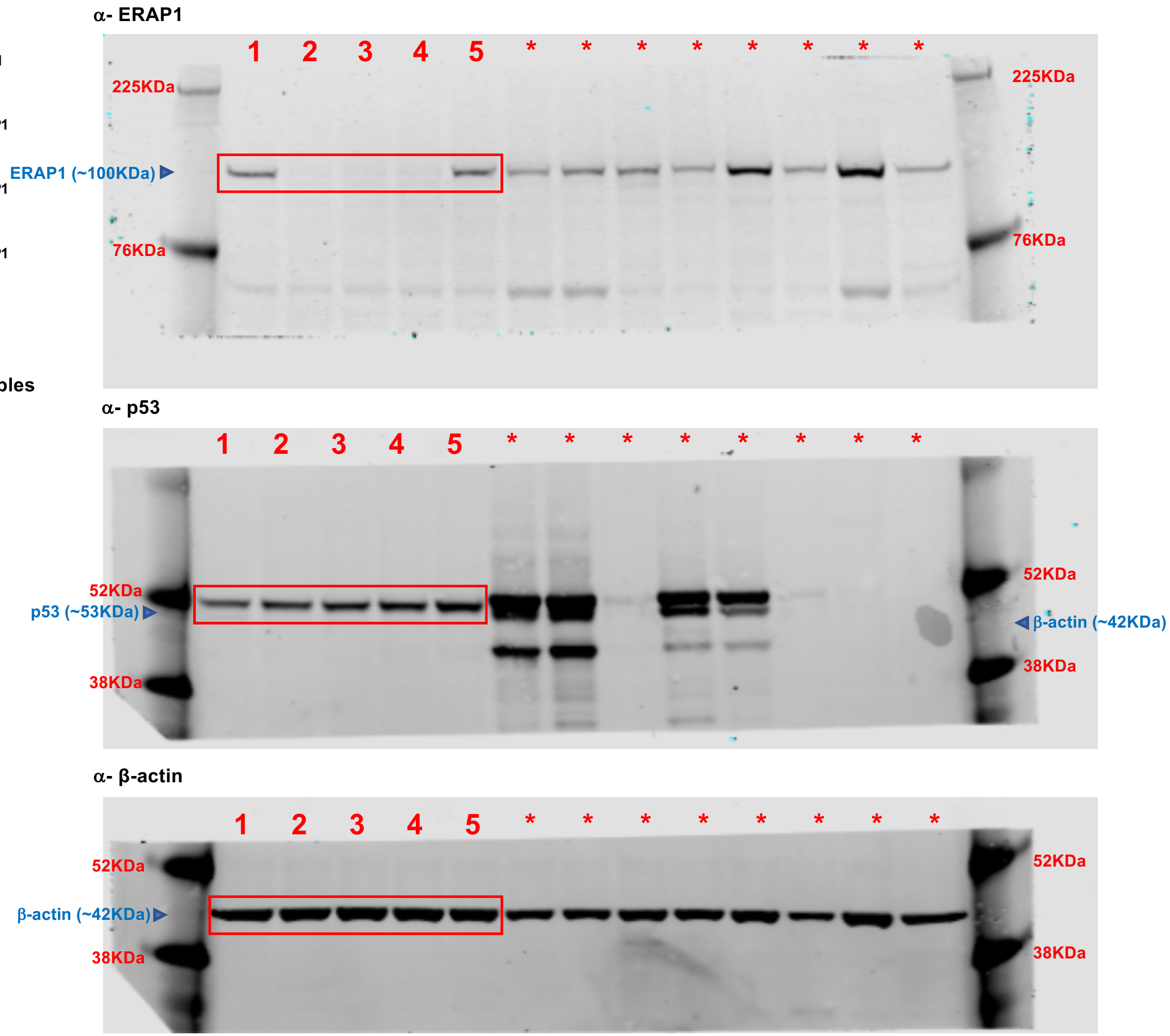


**Supporting data for Fig. 5b.**
Original source images for immunoblotting data used in Fig. 5b.
Gels for ERAP1 and β-actin were derived from the same parent gel.
The same cell lysates were used for p53 in a different gel.
Red rectangles indicate where images were cropped for Fig. 5b.

Supplementary Data File 12: Original source data for immunoblotting

#### c For **Extended Data Fig. 4a**

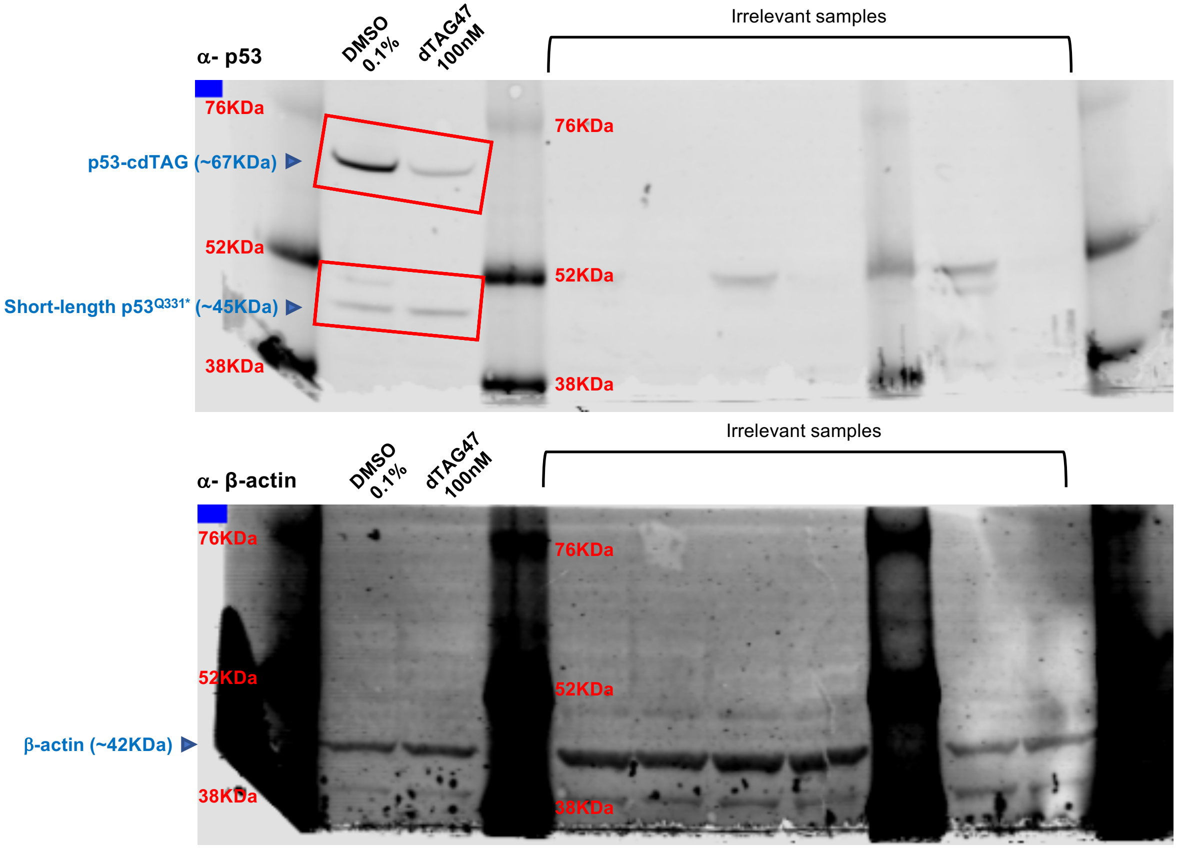


**Supporting data for Extended Data Fig. 4a.**
Original source images for immunoblotting data used in Extended Data Fig. 4a.
Red rectangles indicate where images were cropped for Extended Data Fig. 4a.
Immunoblotting of β-actin using the same cell lysates are also shown for reference.

#### d For **Extended Data Fig. 4f**

1 H2228 treated with DMSO (16 h)

2 H2228 treated with 50 nM of TL-13-122 (16 h)

3 H2228 treated with 100 nM of TL-13-122 (16 h)

4 H2228 treated with 250 nM of TL-13-122 (16 h)

5 H2228 treated with 500 nM of TL-13-122 (16 h)

6 H2228 treated with 1000 nM of TL-13-122 (16 h)

7 H2228 treated with DMSO (1 h)

8 H2228 treated with 300 nM of TL-13-122 (1 h)

9 H2228 treated with DMSO (3 h)

10 H2228 treated with 300 nM of TL-13-122 (3 h)

11 H2228 treated with DMSO (5 h)

12 H2228 treated with 300 nM of TL-13-122 (5 h)

13 H2228 treated with DMSO (7 h)

14 H2228 treated with 300 nM of TL-13-122 (7 h)

15 H2228 treated with DMSO (9 h)

16 H2228 treated with 300 nM of TL-13-122 (9 h)

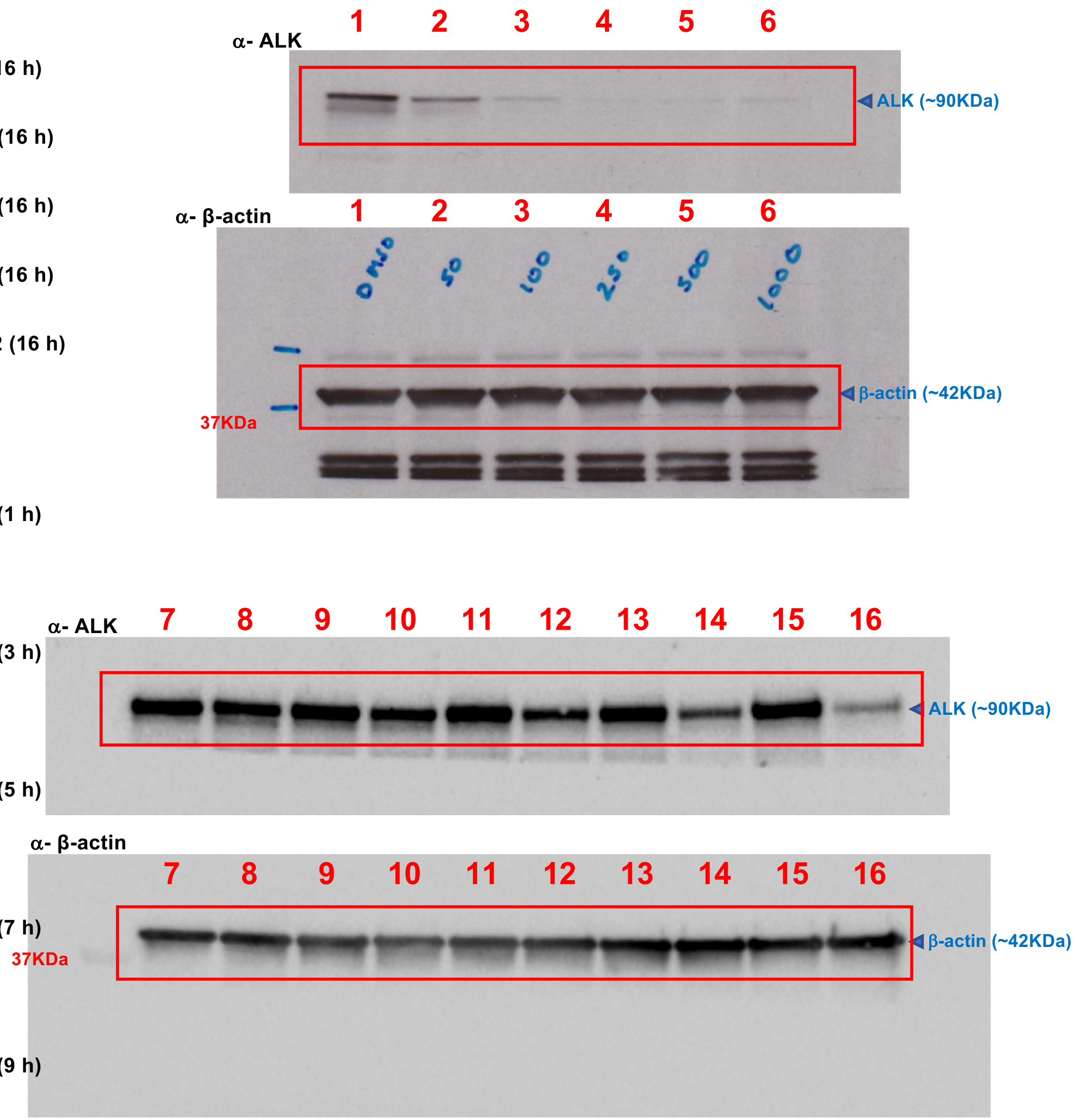


**Supporting data for Extended Data Fig. 4f.**
Original source images for immunoblotting data used in Extended Data Fig. 4f.
Gels for ALK and β-actin were derived from the same parent gel.
Red rectangles indicate where images were cropped for Extended Data Fig. 4f.

Supplementary Data File 12: Original source data for immunoblotting

e For **Extended Data Fig. 7c**

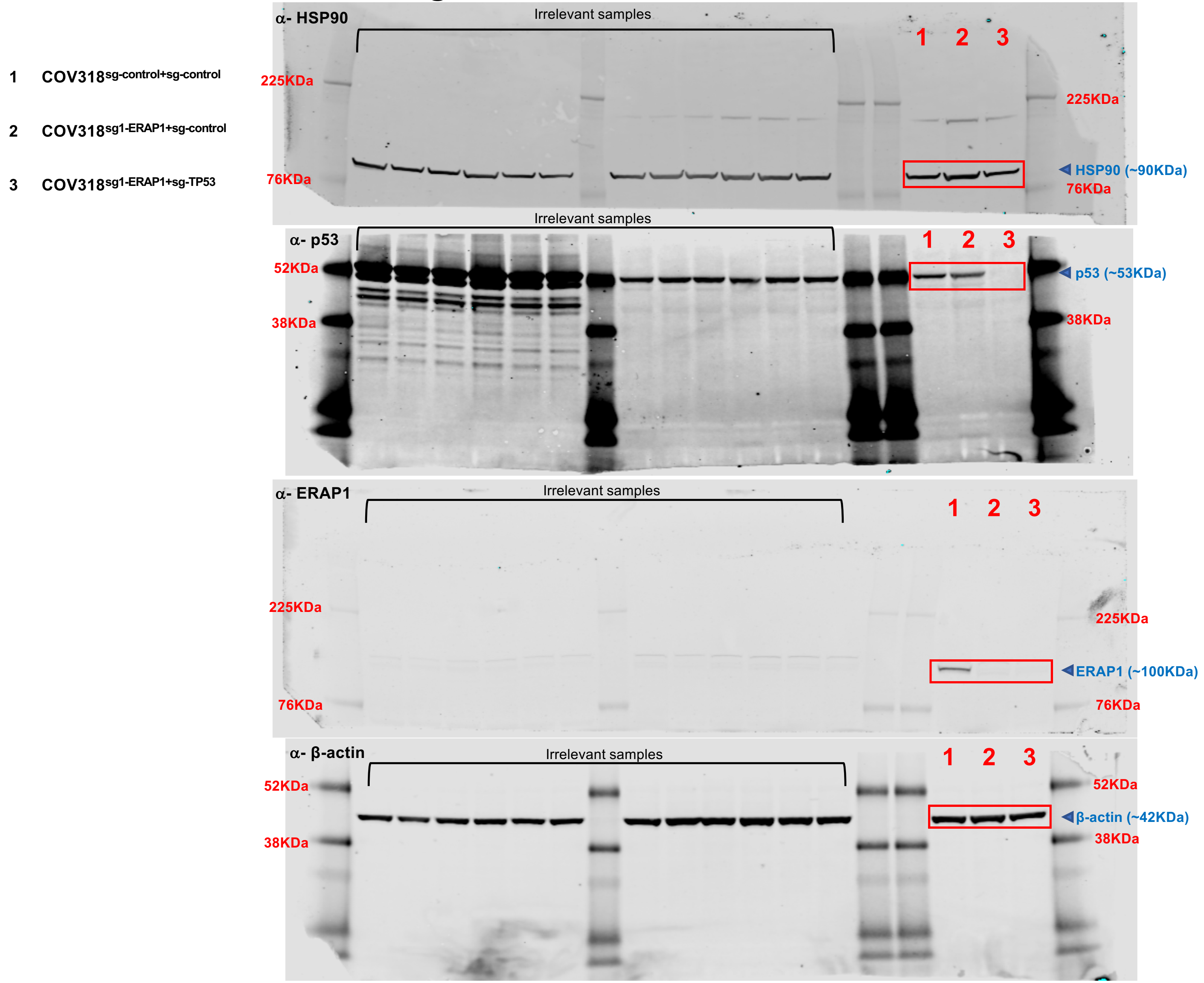


**Supporting data for Extended Data Fig. 7c.** Original source images for immunoblotting data used in Extended Data Fig. 7c. Gels for HSP90 and p53, and those for ERAP1 and β-actin were derived from the same gels, respectively. The same cell lysates were used across all gels. Red rectangles indicate where images were cropped for Extended Data Fig. 7c.

### f For **Extended Data Fig. 7e**

1 CT26$^{sg\text{-}control}$

2 CT26$^{Erap1\text{-}sg1}$

3 CT26$^{Erap1\text{-}sg2}$

4 CT26$^{Erap1\text{-}sg3}$

5 CT26$^{Erap1\text{-}sg4}$

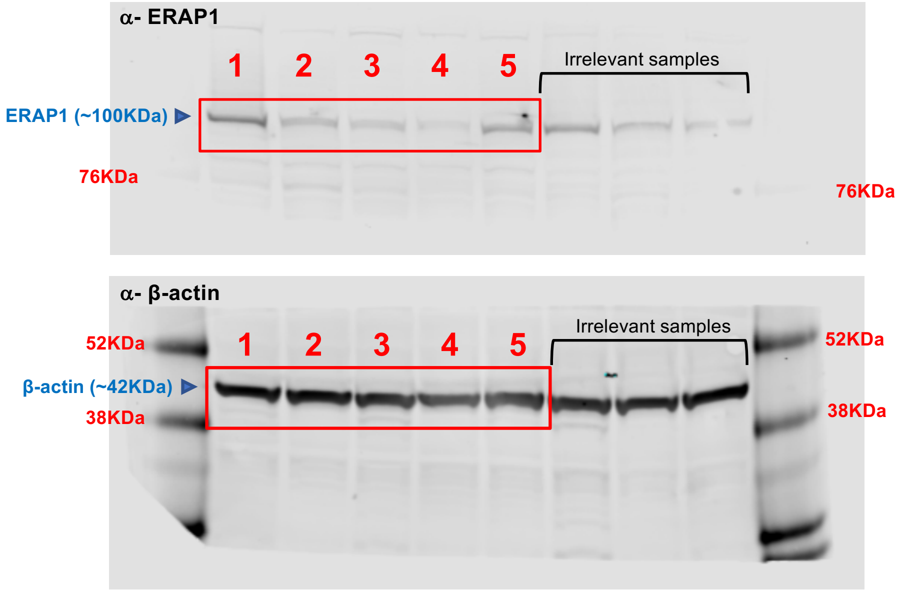


**Supporting data for Extended Data Fig. 7e.**

Original source images for immunoblotting data used in Extended Data Fig. 7e.

Gels for HSP90 and p53, and those for ERAP1 and β-actin were derived from the same gels, respectively.

The same cell lysates were used across all gels.

Red rectangles indicate where images were cropped for Extended Data Fig. 7e.

#### g For **Extended Data Fig. 8c**

1 **H2228**$^{p304\text{-}ffLuc}$

2 **H2228**$^{p304\text{-}ffLuc\text{-}p307\text{-}p53\text{-}R175H}$

3 **H2228**$^{p304\text{-}ffLuc\text{-}p307\text{-}p53\text{-}I195F}$

4 **H2228**$^{p304\text{-}ffLuc\text{-}p307\text{-}p53\text{-}I195F}$

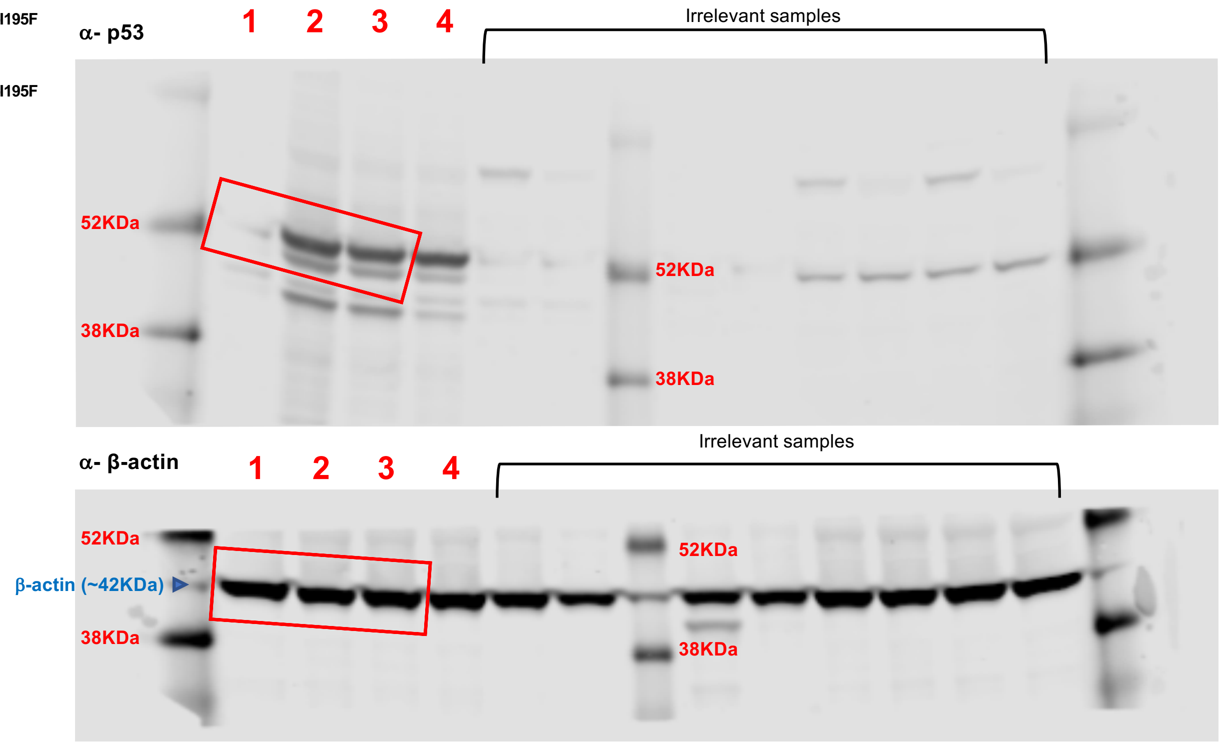


**Supporting data for Extended Data Fig. 8c.**
Original source images for immunoblotting data used in Extended Data Fig. 8c.
The same cell lysates were used in two different gels.
Red rectangles indicate where images were cropped for Extended Data Fig. 8c.

### Supplementary Data File 12: Original source data for immunoblotting

#### h For **Supplementary Data File 6c**

1 Primary T cells from the RG2684 donor

2 PANFR0583 (1st biological replicate)

3 PANFR0583 (2nd biological replicate)

4 H2228p305cdTAG-p53 + DMSO 0.1% (as a positive control)

5 H2228p305cdTAG-p53 + dTAG47 100mM (as a positive control)

* Irrelevant samples

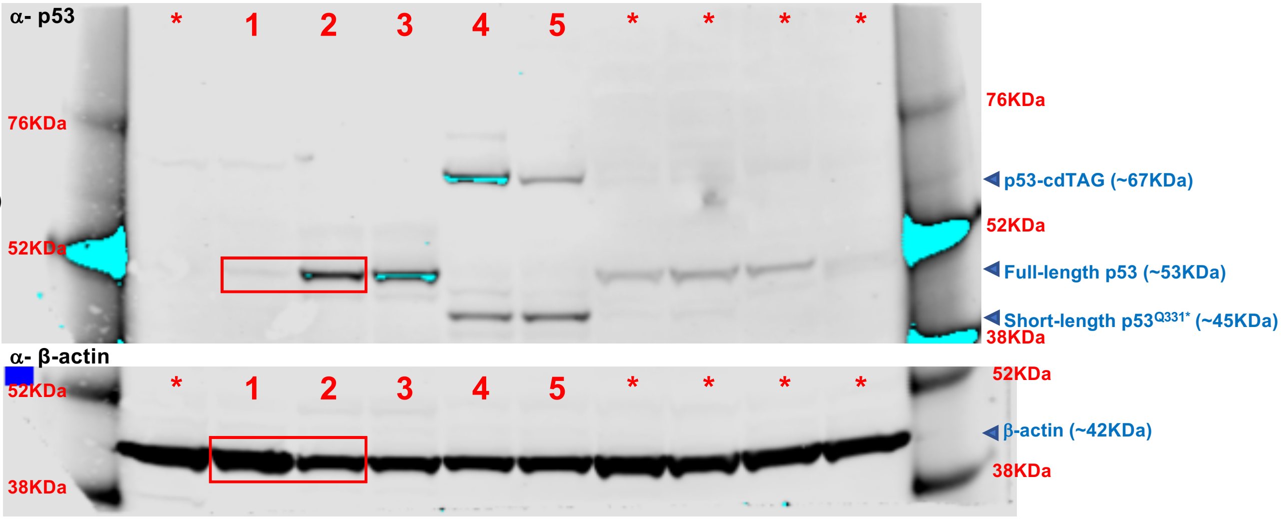


6 PANFR0583 (2nd biological replicate)

7 COV318

8 H2228p305cdTAG-p53

* Irrelevant samples

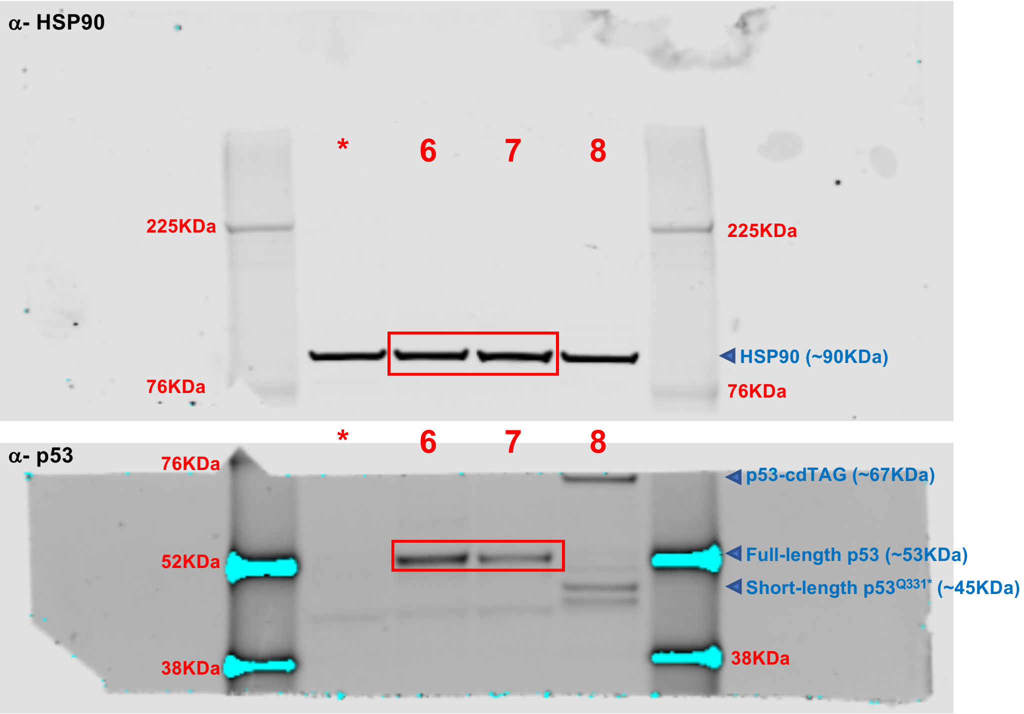


**Supporting data for Extended Data Fig. 6c.** Original source images for immunoblotting data used in Extended Data Fig. 6c. The same lysates were used in the two different top gels for samples 1–5. The two bottom gels for HSP90 and p53 in samples 6–8 were derived from the same gels. Samples 2 and 6 were the same lysates. Red rectangles indicate where images were cropped for Extended Data Fig. 6c.
