## Supplementary Data File 13 for "Cancers modulate p53 truncal neoantigen display to evade T cell detection"

### Supplementary Data File 13: Flow cytometry gating strategy

**a** For **Fig. 4b**

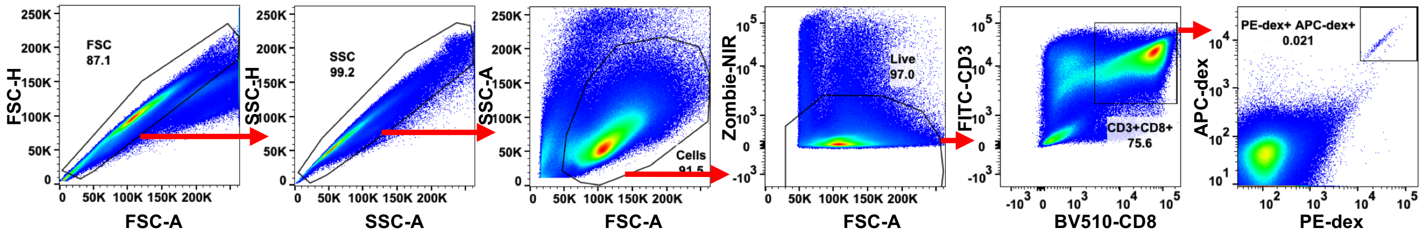

### Supplementary Data File 13: Flow cytometry gating strategy

**b** For **Fig. 4d**, and **Supplementary Data File 9**

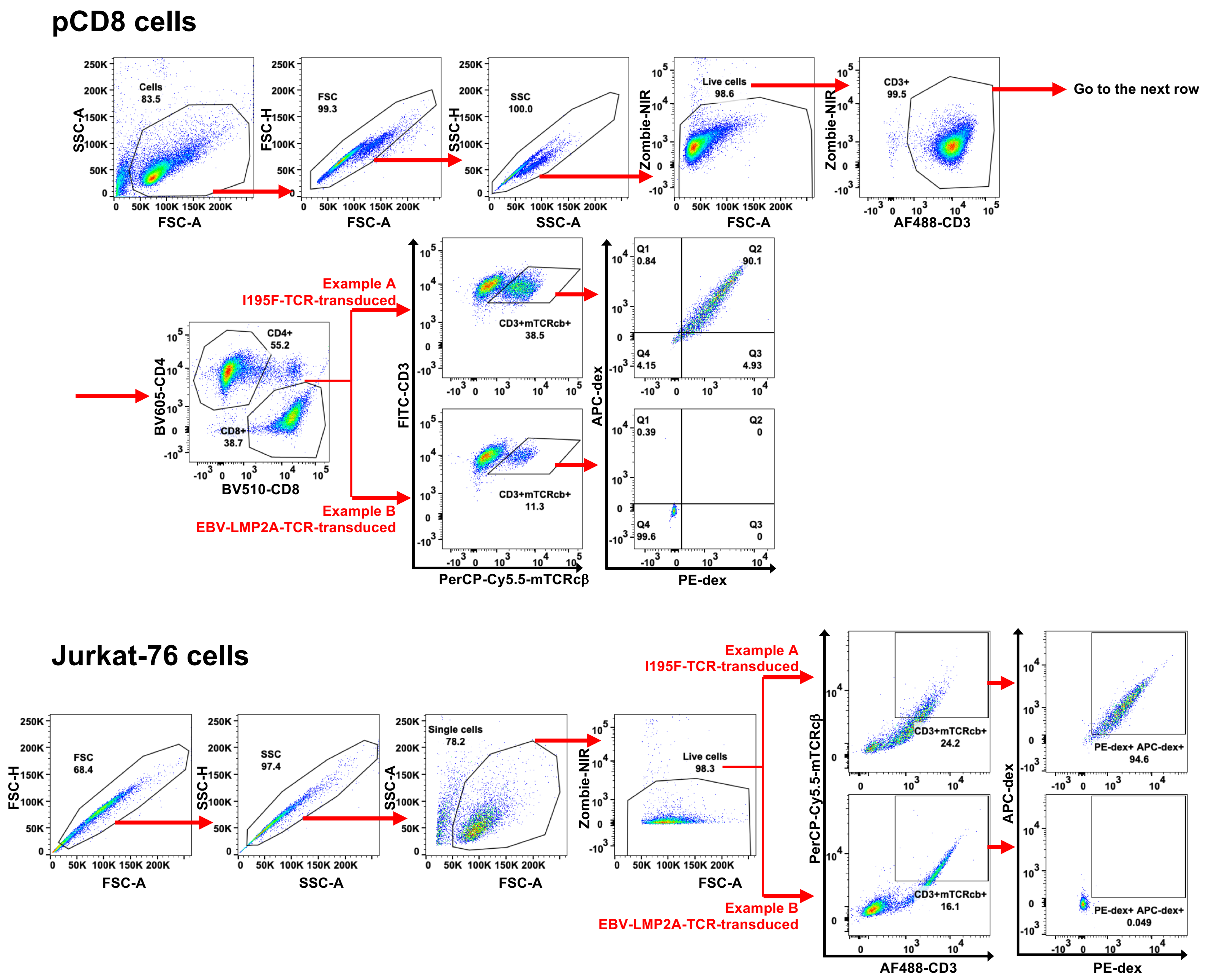

#### Supplementary Data File 13: Flow cytometry gating strategy

**c** For **Fig. 4e**, **Fig. 5**, **Extended Data Fig. 7a**, and **Extended Data Fig. 8a/d**

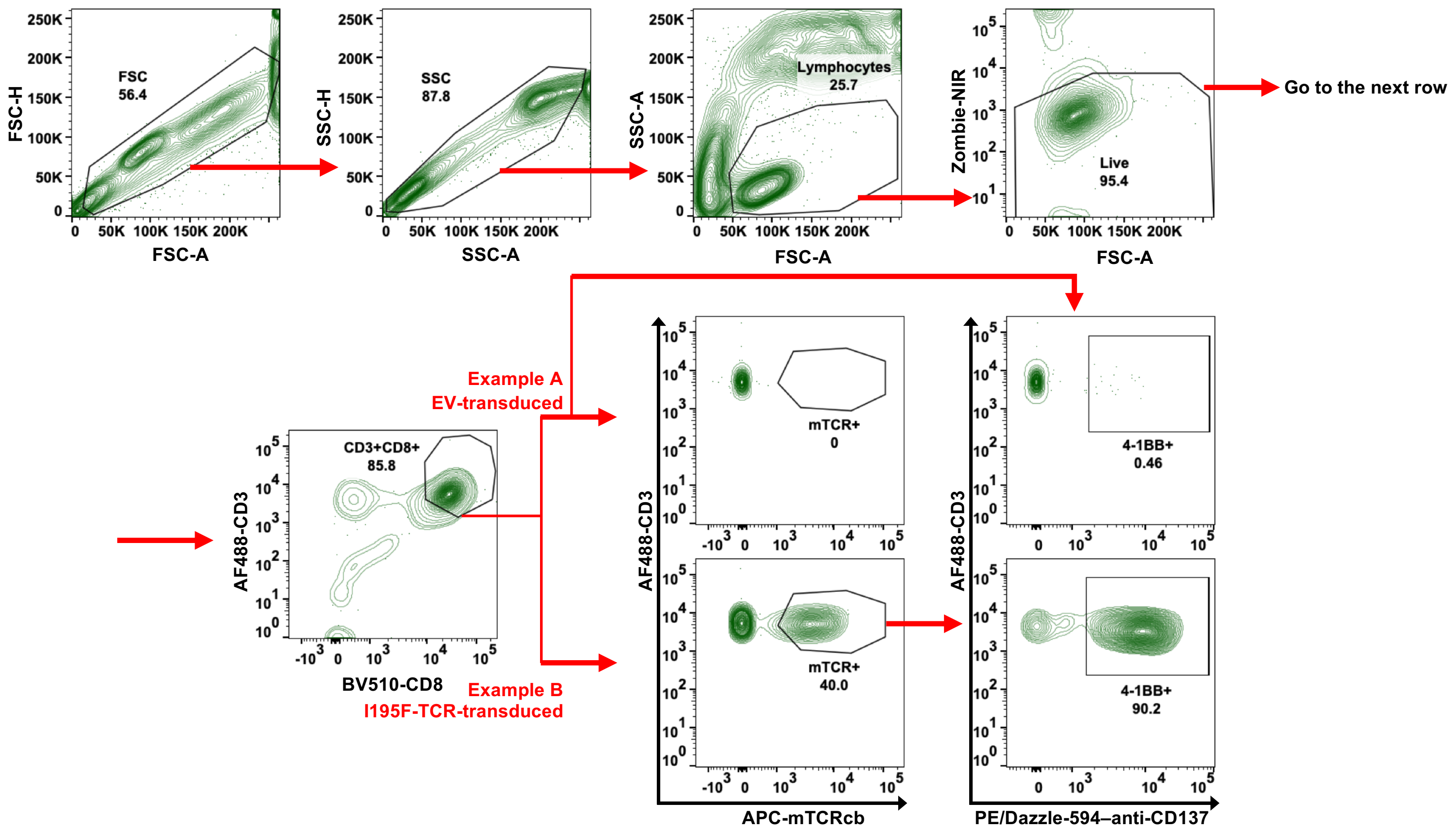

**d** For **Fig. 6e**, **Extended Data Fig. 7d**, **Extended Data Fig. 8c**, **Supplementary Data File 3b**, **Supplementary Data File 6d**, and **Supplementary Data File 7**

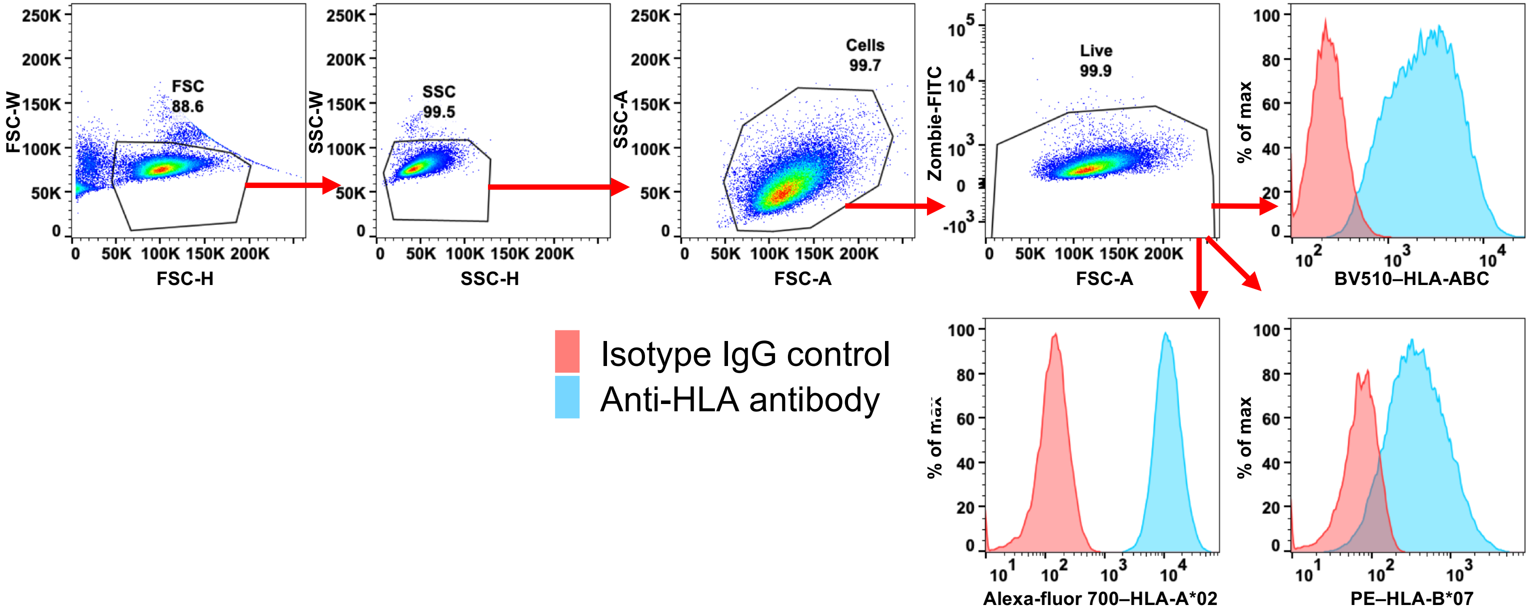

**e** For **Extended Data Fig. 7f**

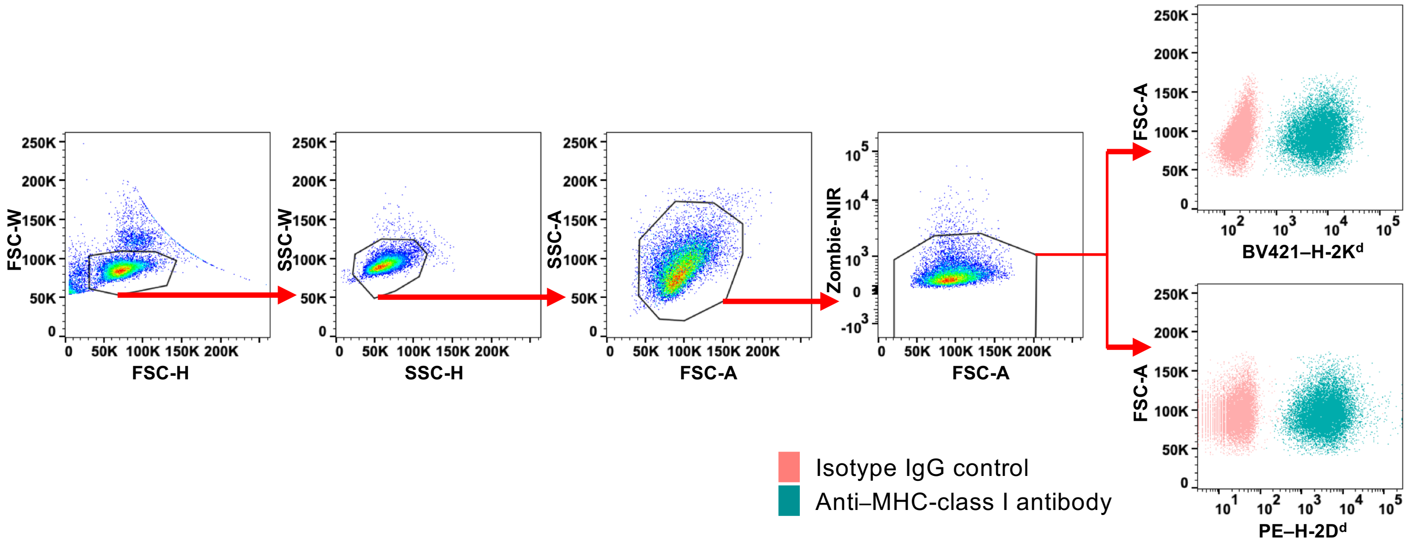

### Supplementary Data File 13: Flow cytometry gating strategy

**f** For **Extended Data Fig. 8e**

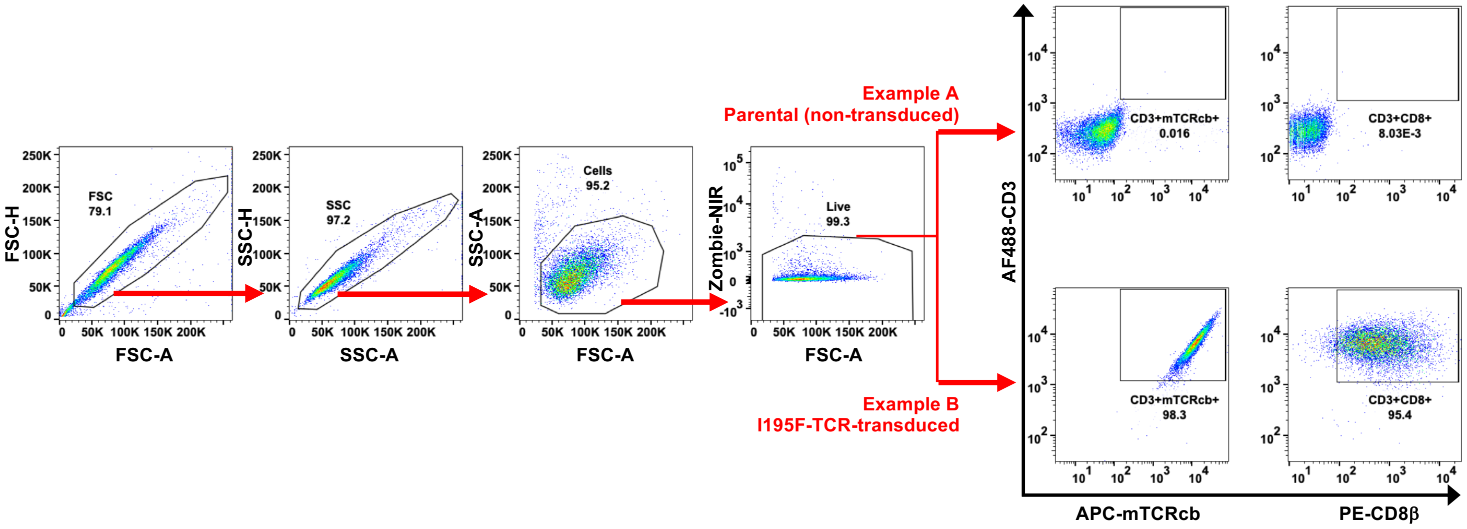
